# Hierarchical and dynamic regulation of defense-responsive specialized metabolism by WRKY and MYB transcription factors

**DOI:** 10.1101/700583

**Authors:** Brenden Barco, Nicole K. Clay

## Abstract

The plant kingdom produces hundreds of thousands of specialized bioactive metabolites, some with pharmaceutical and biotechnological importance. Their biosynthesis and function have been studied for decades, but comparatively less is known about how transcription factors with overlapping functions and contrasting regulatory activities coordinately control the dynamics and output of plant specialized metabolism. Here, we performed temporal studies on pathogen-infected intact host plants with perturbed transcription factors. We identified WRKY33 as the condition-dependent master regulator and MYB51 as the dual functional regulator in a hierarchical gene network likely responsible for the gene expression dynamics and metabolic fluxes in the camalexin and 4-hydroxy-indole-3-carbonylnitrile (4OH-ICN) pathways. This network may have also facilitated the regulatory capture of the newly evolved 4OH-ICN pathway in *Arabidopsis thaliana* by the more-conserved transcription factor MYB51. It has long been held that the plasticity of plant specialized metabolism and the canalization of development (Waddington, 1942) should be differently regulated; our findings imply a common hierarchical regulatory architecture orchestrated by transcription factors for specialized metabolism and development, making it an attractive target for metabolic engineering.

## Introduction

Plants are engaged in a continuous co-evolutionary struggle for survival with their pathogens. Although they lack mobile defender cells and an adaptive immune system, they rely on their innate immune system to collectively synthesize hundreds of thousands of ecologically specialized, mostly lineage-specific, preformed and pathogen-inducible metabolites at sites of infection (Chae *et al.,* 2014; Weng *et al.,* 2012; Dixon and Strack, 2003; Wink, 2003). Pathogen-inducible specialized metabolites are synthesized under two primary modes of plant innate immunity – pattern- and effector-triggered immunity (PTI and ETI). PTI depends on signaling networks that identify the non-self microbial invader via its conserved microbe-associated molecular pattern molecules (MAMPs), whereas ETI utilizes pathogen-specific virulence effector proteins for pathogen detection (Jones and Dangl, 2006). Specialized metabolism is further dependent on gene regulatory networks (GRNs) that respond to perceived threats by activating defense-responsive transcription factors (TFs) (Clay *et al.,* 2009; Chezem *et al.,* 2017; Barco *et al.,* 2019b) and suppressing TFs involved in growth and development (Lozano-Durán *et al.,* 2013; Fan *et al.,* 2014; Malinovsky *et al.,* 2014; Lewis *et al.,* 2015).

TFs are ultimately responsible for controlling the dynamics and output of gene expression in plant specialized metabolism, and genes encoding specialized metabolic enzymes are often organized into regulons, whereby they come under the control of a limited set of TFs for optimal timing, amplitude, and tissue/pathway-specific expression and subsequent metabolite accumulation (Grotewold, 2005; Hartmann, 2007; Martin *et al.,* 2010; Tohge and Fernie, 2012; Omranian *et al.,* 2015). However, transcription networks that are responsive to external perturbations often contain many TFs with overlapping functions and contrasting regulatory activities, as well as regulons that often include diverse targets (e.g. genes encoding other TFs, metabolic enzymes for multiple pathways, and non-enzymatic proteins). GRNs are thus elaborate, supercoordinated forms of organization which connect primary and secondary metabolism, environmental signals and physiological responses such as growth and defense (Aharoni and Galili, 2011; Baghalian *et al*., 2014). Subsequently, the ability to engineer novel plant specialized metabolism more often than not produces a frustrating array of unanticipated and undesirable outcomes to the system (Colón et al., 2010; Bonawitz and Chapple, 2013).

Much progress has been made in understanding the finer details of GRN architecture. Central to GRN organization are small sets of recurring regulatory circuits called network motifs (Milo *et al*., 2002; Shen-Off *et al.,* 2002). Each motif has been experimentally found to perform specific dynamical functions in gene expression and is wired into the network in such a way that preserves its autonomous functions in natural contexts, thus predictions of network dynamics can be made with simple network motifs of core components without precise knowledge of all of the underlying parameters (Alon, 2007; Gutenkunst *et al.,* 2007). One of the most prevalent network motifs in the GRNs of *Escherichia coli* (Shen-Orr *et al.,* 2002; Ma *et al.,* 2004), *Saccharomycetes cerevisiae* (Lee *et al.,* 2002; Mangan *et al.,* 2006), mammalian cells (Ma’ayan *et al.,* 2005; Odom *et al.,* 2004; Boyer *et al.,* 2005), and *Arabidopsis thaliana* (*A. thaliana*) (Jin *et al.,* 2015; Defoort *et al.,* 2018) is the three-component feed-forward loop (FFL), which is composed of two cascaded TFs that interact at a target promoter and jointly determine its rate of transcription (Milo *et al.,* 2002; Mangan and Alon, 2003; Alon, 2007). Depending on the nature of the feed-forward regulation of the target gene (e.g., activation and/or repression, AND- and/or OR-logic gating), the FFL architecture has been shown to exhibit four types of expression dynamics: 1) memory effects of input signals, such as persistence detection (noise filtering), fold-change detection, and dynamics detection (Mangan and Alon, 2003; de Ronde *et al.,* 2012; Goentoro *et al.,* 2009; Lee *et al.,* 2014; Chepyala *et al.,* 2016; Gao *et al.,* 2018); 2) temporal effects of target gene responses such as fast or delayed activation and inhibition, oscillations, and (near-)perfect adaptive pulses (Mangan and Alon, 2003; Mangan *et al.,* 2003; Mangan *et al*., 2006; Basu *et al.,* 2004; Basu *et al.,* 2005; Cournac and Sepulchre, 2009; Ma *et al.,* 2009; Sontag *et al.,* 2009; Takeda *et al.,* 2012); 3) amplitude- and pulse-filtering of target gene responses (Shen-Orr *et al.,* 2002; Mangan and Alon, 2003; Mangan *et al.,* 2003; Kaplan *et al.,* 2008); and 4) irreversible switches and transitions (Jaeger *et al.,* 2013; Pullen *et al.,* 2013; Lavenus *et al.,* 2015). FFL circuits can also exhibit two dynamical functions in a network, for example, noise-filtering and irreversibility by using OR-logic gating of target gene responses for transcriptional activation at a reduced level and AND-logic gating for maximal transcriptional activation (Pullen *et al.,* 2013).

GRNs for growth and development are defined by interlinked or clustered FFLs in animals (Gerstein *et al.,* 2010; modEncode Consortium *et al.,* 2010; Cheng *et al.,* 2011; Niu *et al.,* 2011), plants (Lin *et al.,* 2013; Jaeger *et al.,* 2013; Lavenus *et al.,* 2015; Taylor-Teeples *et al.,* 2015; Joanito *et al.,* 2018; Zhan *et al.,* 2018; Chen *et al.,* 2019), and *E. coli* (Semsey *et al.,* 2007). Nonetheless, such networks for stress-responsive plant specialized metabolism are still largely defined by individual TFs and their overlapping regulons (Li *et al.,* 2014; James *et al.,* 2017; Yang *et al.,* 2017). Little is known about the hierarchical network motifs that enable multiple TFs with activating and repressive functions to coordinately control the dynamics and output of gene expression and metabolic flux in this context.

The best-studied defense-responsive specialized metabolites in *A. thaliana* with demonstrated immune functions against fungal and bacterial pathogens are the tryptophan (Trp)-derived camalexin, 4-methoxyindol-3-ylmethyl glucosinolate (4M-I3M), and 4-hydroxyindole-3-carbonylnitriles (4OH-ICN) (Bohman *et al.,* 2004; Bednarek *et al.,* 2009; Clay *et al.,* 2009; Consonni *et al.,* 2010; Ferrari *et al.,* 2003; Hiruma *et al.,* 2010; Lipka *et al*., 2005; Pandey *et al.,* 2010; Rajniak *et al.,* 2015; Sanchez-Vallet *et al.,* 2010; Schlaeppi *et al.,* 2010; Thomma *et al.,* 1999). 4M-I3M, its immediate precursor 4-hydroxy-I3M (4OH-I3M), and sister metabolite 1-methoxy-I3M, are all derived from the parent molecule I3M, and are collectively known as indole glucosinolates (indole GSLs). The biosynthetic pathways of 4M-I3M, camalexin and 4OH-ICN share an early Trp-to-indole-3-acetaldoxime (IAOx) biosynthetic step, courtesy of the genetically redundant cytochrome P450 monooxygenases (CYPs) CYP79B2–3 (Mikkelsen *et al.,* 2000; Glawischnig *et al.,* 2004; Rajniak *et al.,* 2015). CYP71 clade enzymes CYP83B1 and partially redundant CYP71A12/13 respectively convert IAOx to short-lived *aci*-nitro intermediates (ANI) and indole-3-cyanohydrin (ICY) (Figure 1A) (Bak *et al.,* 2001; Nelson and Werck-Reichhart, 2011; Klein *et al.,* 2013; Rajniak *et al.,* 2015; Barco *et al.,* 2019c). CYP71A13 and CYP71B15/PAD3 convert ICY to camalexin, while flavin-dependent oxidase FOX1/AtBBE3 and 4-hydroxylase CYP82C2 convert ICY to 4OH-ICN (Figure 1A) (Nafisi *et al.,* 2007; Böttcher *et al.,* 2009; Rajniak *et al.,* 2015). 4M-I3M is synthesized from ANI via glutathione-*S*-transferases GSTF9–10, γ-glutamyl peptidase GGP1, *S*-alkyl-thiohydroximaste lyase SUR1, UDP-glycosyltransferase UGT74B1, sulfotransferase SOT16, 4-hydroxylases CYP81F1–3, and I3M methyltransferases IGMT1–2 (Figure 1A) (Chezem and Clay, 2016; Barco *et al.,* 2019a).

**Figure 1.**
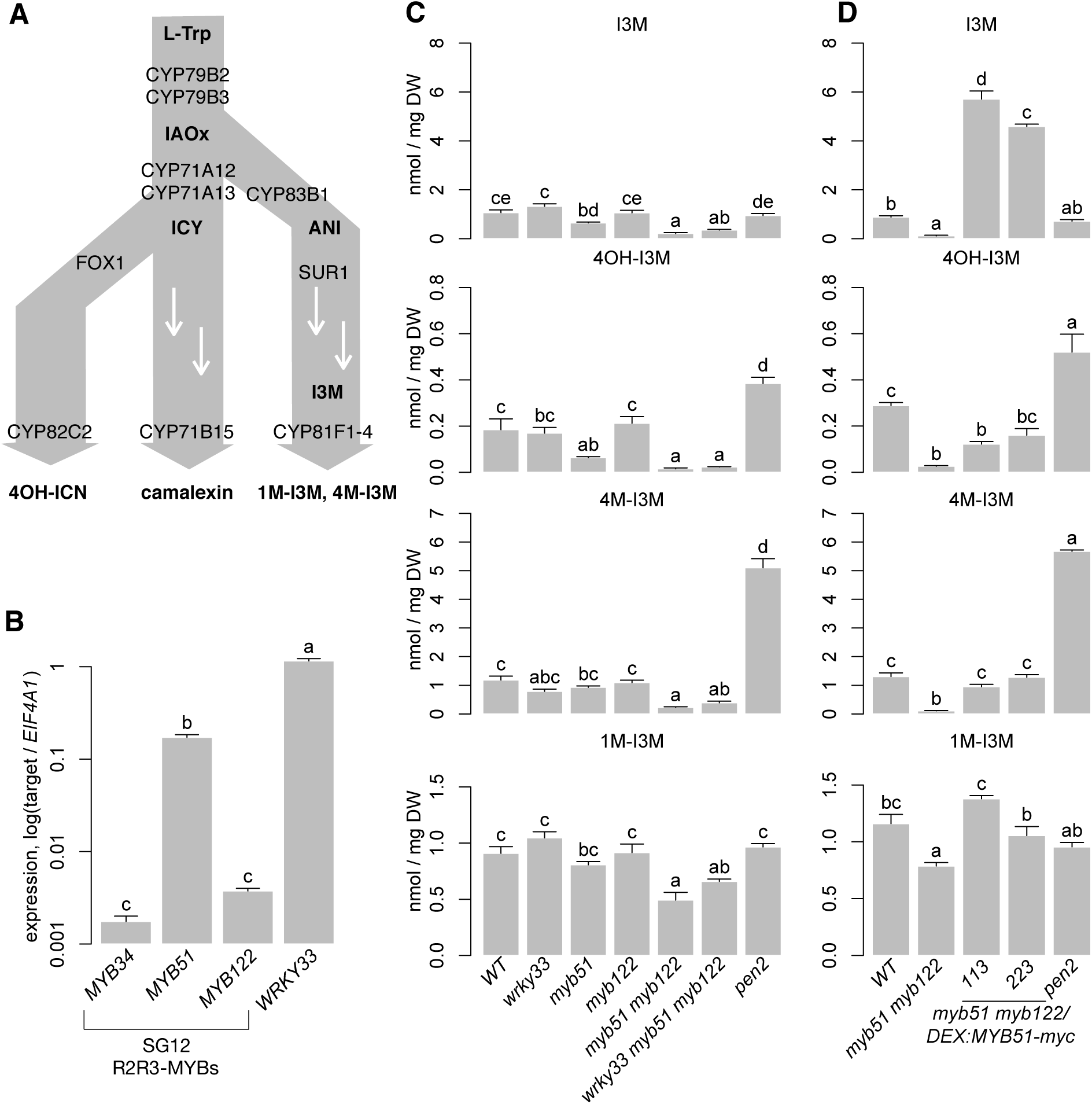
*MYB51/MYB122* are necessary for 4OH-I3M and 4M-I3M biosynthesis in ETI. **(A)** Schematic of tryptophan (L-Trp)-derived secondary metabolite pathways in Arabidopsis. 1M-I3M, 1-methoxyindol-3-ylmethyl glucosinolate; 4M-I3M, 4-methoxyindol-3-ylmethyl glucosinolate; 4OH-ICN, 4-hydroxyindole-3-carbonylnitrile; ANI, aci-nitro indole; I3M, indol-3-ylmethyl glucosinolate; IAOx, indole-3-acetaldoxime; ICY, indole-3-cyanohydrin.
**(B)** qPCR analysis of indole GSL regulatory genes and WRKY33 in 9-day-old WT seedlings co-elicited with 20 μM Dex and *Psta* for 12 hr. Data represent mean ± SE of four replicates of 13-17 seedlings each. Expression values were normalized to that of the housekeeping gene *EIF4A1*.
**(C)** and **(D)** HPLC-DAD analysis of I3M, 4OH-I3M, 4M-I3M, and 1M-I3M (top to bottom) in 9-day-old seedlings elicited with *Psta* for 24 hr **(C)** or co-elicited with 20 μM Dex and *Psta* for 24 hr **(D)**. DW, dry weight. The *pen2* mutant cannot hydrolyze indole GSLs and thus over-accumulates defense-induced 4OH-I3M and 4M-I3M (Bednarek *et al.,* 2009; Clay *et al.,* 2009). Data represent the mean ± SE of four replicates of 13-17 seedlings each. Different letters in (**B**-**D**) denote statistically significant differences (*P* < 0.05, one-factor ANOVA coupled to Tukey’s test). Experiments were performed twice, producing similar results.

The SG12-type R2R3-MYB TFs MYB51 and MYB122 and the Group I WRKY TF WRKY33 are among the best-characterized defense-responsive TFs in *A. thaliana*. MYB51 and MYB122 are activators of 4M-I3M biosynthesis and are required for basal resistance to a variety of bacterial and fungal pathogens (Gigolashvili *et al.,* 2007a; Malitsky *et al.,* 2008; Clay *et al.,* 2009; Humphry *et al.,* 2010; Frerigmann and Gigolashvili, 2014a; Lahrmann *et al.,* 2015; Frerigmann *et al.,* 2016). MYB51 and MYB122 also contribute to camalexin biosynthesis in response to UV stress and the fungal necrotroph *Plectosphaerella cucumerina* through *trans*-activation of *CYP79B2* and *CYP79B3* promoters (Frerigmann *et al.,* 2015, 2016). WRKY33 is an activator of camalexin and 4OH-ICN biosynthesis in response to the ETI-eliciting bacterial pathogen *Pseudomonas syringae* (*Pst*) *avrRpm1* and the fungal necrotroph *Botrytis cinerea* (*B. cinerea*) (Qiu *et al.,* 2008; Birkenbihl *et al*., 2012; Liu *et al.,* 2015; Birkenbihl *et al*., 2017; Barco *et al.,* 2019b) and is required for basal resistance to *Pst* and *B. cinerea* (Zheng *et al.,* 2006; Barco *et al.,* 2019b).

To understand how TFs with variable functions and activities coordinately and dynamically govern plant specialized metabolism, we performed temporal studies employing an ETI-eliciting pathogen on host plants exhibiting gain or loss of TF expression. Hydroponically- and sterilely-grown naïve (unprimed) seedlings were tested to better synchronize the infection process and reduce stress memory effects. We identified a composite hierarchical network motif with WRKY33 as the condition-dependent master regulator and MYB51 as the dual functional regulator that is likely responsible for the gene expression dynamics and metabolic fluxes through the CYP79B2/B3- and CYP82C2-catalyzed steps in the camalexin and/or 4OH-ICN pathways. The characterization of these TF activities in hierarchical gene circuits - in particular how targets are dynamically and coordinately controlled - will better inform how new biosynthetic pathways can be engineered or evolved.

## Results

### MYB51 and MYB122 are necessary for 4OH-I3M and 4M-I3M biosynthesis in ETI

Previous studies have shown that MYB34, MYB51, and MYB122 distinctly regulate indole GSL biosynthesis in response to plant hormones. MYB51 is the central regulator of indole GSL synthesis upon salicylic acid and ethylene (ET) signaling, MYB34 is the key regulator upon abscisic acid and jasmonic acid (JA) signaling, and MYB122 has a minor role in JA/ET-induced indole GSL biosynthesis (Frerigmann and Gigolashvili, 2014a). In addition, MYB51 is the major regulator of pathogen-/MAMP-induced 4M-I3M biosynthesis, with MYB122 having a minor role (Clay *et al.,* 2009; Frerigmann *et al.,* 2016). To identify the *A. thaliana* SG12-type R2R3-MYB regulator(s) of 4M-I3M biosynthesis in ETI, we compared the host transcriptional response to PTI-eliciting bacterial MAMP flagellin epitope flg22 with that to ETI-eliciting bacterial pathogen *Pst avrRpm1* (*Psta*) under similar conditions as those of previous studies (Denoux *et al.,* 2008; Clay *et al.,* 2009; Rajniak *et al.,* 2015). *MYB51* and *MYB122* were induced in response to flg22, with *MYB51* expression increasing as high as 50-fold (Supplemental Table 1) (Denoux *et al.,* 2008). Similarly, *MYB51* was about 100-fold induced in response to *Psta* (Figure 1B). The observed expression of *MYB51* and *MYB122* and quantitative differences in transcriptional responses between flg22 and *Psta* are consistent with previous transcriptional studies of other PTI and ETI elicitors (Supplemental Table 1; Tao *et al.,* 2003; Navarro *et al.,* 2004; Toufighi *et al.,* 2005; Austin *et al.,* 2016; Frerigmann *et al.,* 2016).

To assess functional redundancy between MYB51 and MYB122 in activating 4M-I3M biosynthesis in ETI, we compared the host metabolic response to *Psta* in wild-type (WT), a loss-of-function *myb51* transposon insertion mutant, a newly isolated loss-of-function *myb122-3* T-DNA insertion mutant, and the *myb51 myb122-3* double mutant (Supplemental Figure 1A). We previously have shown that 4M-I3M and its immediate precursor 4-hydroxy-I3M (4OH-I3M) levels were increased at the expense of the parent metabolite I3M in *Psta*-infected WT plants compared to uninfected WT or *Psta*-infected *rpm1* mutant, which is ETI-deficient when elicited with *Psta* (Bisgrove *et al.,* 1994; Barco *et al.,* 2019b). By contrast, I3M and 4OH-I3M levels were reduced in the *Psta*-infected *myb51* mutant relative to *Psta*-infected WT (Figure 1C), consistent with a previous report of reduced flg22-elicited indole GSL biosynthesis in *myb51* (Clay *et al.,* 2009). *Myb122-3* contains a T-DNA insertion in a region that encodes the DNA-binding R2R3 domain of MYB122. This T-DNA insertion is further upstream than that of the loss-of-function *myb122-2* mutant (Supplemental Figure 1A) (Frerigmann *et al.,* 2014a) and *myb122-3* (hereafter referred to as *myb122*) resembles *myb122-2* in exhibiting WT levels of both *MYB122* transcription upstream of the T-DNA insertion and indole GSL metabolism (Supplemental Figure 1B; Figure 1C) (Frerigmann *et al.,* 2014a). In the *Psta*-infected *myb51 myb122-3* mutant (hereafter referred to as *myb51 myb122*), severe reductions in all indole GSLs – including 1-methoxy-I3M (1M-I3M) – were observed (Figure 1C-D). Consistent with these results, transcript levels of indole GSL core biosynthetic genes *CYP83B1* and *SUR1* were also reduced in *myb51 myb122* (Figure 2A).

**Figure 2.**
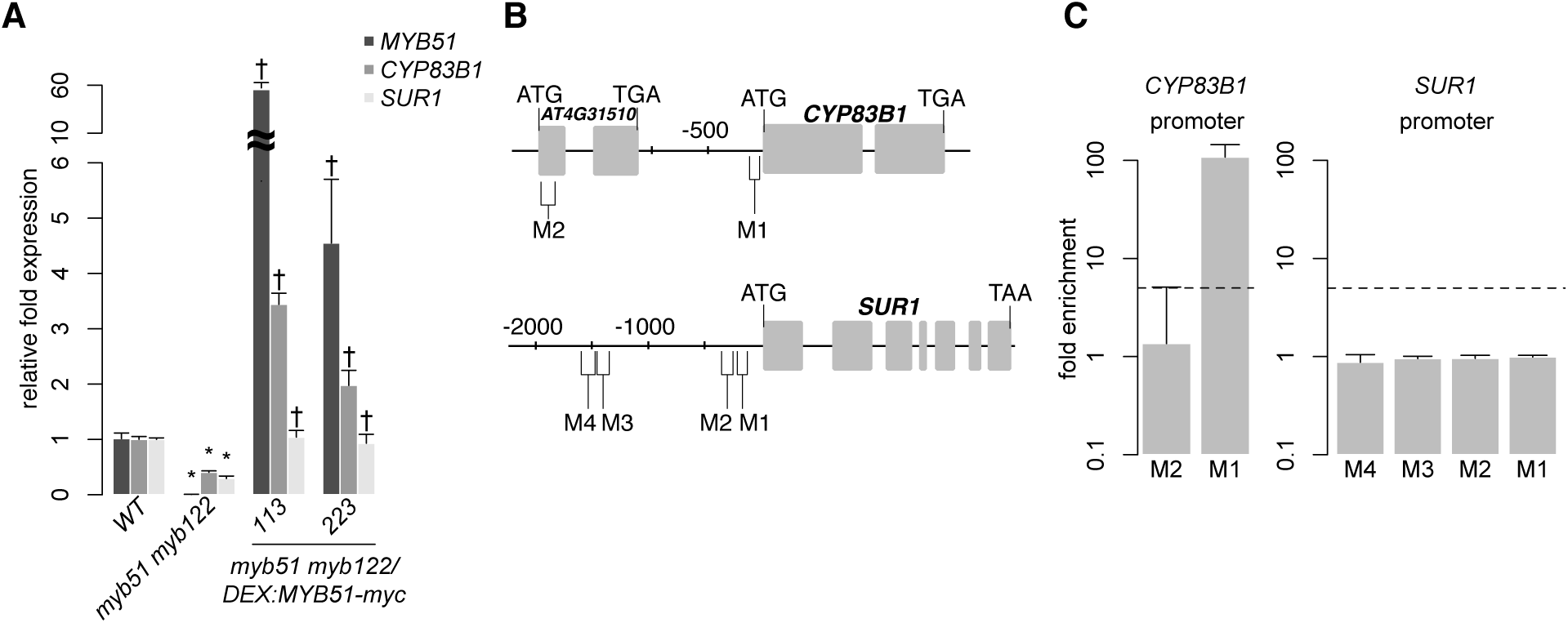
MYB51 directly activates SMRE-containing *CYP83B1* promoter. **(A)** qPCR analysis of *MYB51* and indole GSL biosynthetic genes *CYP83B1* and *SUR1* in 9-day-old seedlings co-elicited with 20 μM Dex and *Psta* for 12 hr. Data represent mean ± SE of four replicates of 13-17 seedlings each. Expression values were normalized to that of the housekeeping gene *EIF4A1*. Asterisks and daggers denote statistically significant differences compared to wild-type and *myb51 myb122*, respectively (*P* < 0.05, two-tailed *t* test).
**(B)** and **(C)** Nucleotide positions **(B)** and ChIP-PCR analysis **(C)** of SMRE (M)-containing promoter regions (pro) bound by MYB51-myc in 9-day-old seedlings co-treated with 20 μM Dex or mock solution (0.5% DMSO) and *Psta* for 9 hr. Boxes in **(B)** indicate exons. Dashed lines represent the 5-fold cutoff between weak and strong TF-DNA interactions. Data in **(C)** represent median ± SE of 4 biological replicates, each containing approximately 200 seedlings. Fold enrichment was determined by calculating the ratio of PCR product intensities in ChIP Dex/Mock to Input Dex/Mock. Experiments were performed twice, producing similar results.

To confirm that MYB51 is sufficient to activate 4OH-I3M and 4M-I3M biosynthesis in ETI, we utilized in the *myb51 myb122* background the two-component glucocorticoid-inducible system (Aoyama and Chua, 1997) to generate plants that in the presence of the glucocorticoid hormone dexamethasone (Dex) express a wild-type copy of the *MYB51* gene with a C-terminal fusion to *6x c-Myc* (*myb51 myb122/DEX:MYB51- myc*) (Figure 2A; Supplemental Figure 1C). Induced expression of *MYB51-myc* increased I3M biosynthesis in the *myb51 myb122* mutant to greater than WT levels by more than four-fold in two independent transgenic lines (Figure 1D), enough to fully restore 4M-I3M and 1M-I3M biosynthesis to WT levels (Figure 1D). Collectively, these results indicate partial functional redundancy between MYB51 and MYB122, with a predominant role for MYB51 in 4OH-I3M and 4M-I3M biosynthesis in ETI.

### MYB51 directly activates SMRE-containing *CYP79B2, CYP79B3*, and *CYP83B1* promoters

The mechanism for the transcriptional regulation of camalexin biosynthesis by MYB51 and MYB122 was previously shown to be centered on *CYP79B2* and *CYP79B3*, which encode two enzymes redundantly involved in an early step in the pathway (Gigolashvilli *et al*., 2007b; Frerigmann *et al*., 2015). Previous transient *trans*-activation assay studies with target promoter-*GUS* reporter genes have shown that indole GSL biosynthetic regulators MYB34 and MYB51 (and MYB122 in the case of *CYP83B1*) directly target secondary indole metabolic genes *CYP79B2* and its functionally redundant homolog *CYP79B3* (which are shared by indole GSL/camalexin/ICN pathways) and indole GSL-specific pathway gene *CYP83B1*. By contrast, aliphatic GSL biosynthetic regulators (MYB28, MYB29 and MYB76) directly target aliphatic GSL biosynthetic genes as well as the biosynthetic gene *SUR1* which is shared by both aliphatic and indole GSL pathways (Gigolashvili *et al.,* 2007b, 2008; Frerigmann *et al.,* 2016).

To confirm that MYB51 *trans*-activates *CYP79B2, CYP79B3*, *CYP83B1* and *SUR1* expression in ETI, we compared host transcriptional response to *Psta* in WT, *myb51 myb122* and *myb51 myb122/DEX:MYB51-myc*. Transcript levels of *CYP79B2*, *CYP79B3,* and *CYP83B1* were reduced in *myb51 myb122* relative to WT but restored to greater than WT levels upon induced expression of *MYB51-myc* (Figures 2A, 3A). Interestingly, *SUR1* transcript level was also reduced in *myb51 myb122* and restored to WT level upon induced expression of *MYB51-myc* (Figure 2A).

SG12-type R2R3-MYBs are closely related to R2R3-MYBs that bind to type IIG Myb recognition sequences [(T/C)ACC(A/T)A(A/C)C] in electrophoretic mobility shift assays (Romero *et al.,* 1998) and to a shorter 7-bp secondary wall MYB-responsive element (SMRE) consensus sequence [ACC(A/T)A(A/C)(T/C)] within the type IIG Myb recognition sequence in *trans*-activation and chromatin immunoprecipitation (ChIP) assays (Zhou *et al.,* 2009; Zhong and Ye, 2012; Chezem *et al.,* 2017). Moreover, all indole GSL pathway gene promoters contain one or more SMREs (Figures 2B, 3B).

To determine whether MYB51 directly binds to SMRE-containing promoter regions of indole GSL pathway genes, we performed ChIP on 9-hr *Psta*-infected mock- and Dex-treated *myb51 myb122/DEX:MYB51-myc* seedlings using antibodies specific to c-Myc (Supplemental Figure 1D). We then PCR-amplified SMRE-containing regions within 2000 nt upstream of the translational start site (TSS) of genes encoding the first three enzymes in the indole GSL pathway: *CYP79B2, CYP79B3, CYP83B1* and *SUR1*. Consistent with a previous report which utilized transactivation assays (Gigolashvili *et al.,* 2007a), MYB51 bound strongly (>10-fold enrichment) to the proximal SMRE-containing regions of *CYP79B2* (M1)*, CYP79B3* (M1) and *CYP83B1* (M1) promoters (Figures 2B-2C, 3B-3C; Supplemental Figures 2, 3A-B; Supplemental File 1).

Interestingly, all three proximal SMRE-containing regions contain SMRE motifs that are identical to the three AC elements [AC-I (ACCTACC), AC-II (ACCAACC), and AC-III (ACCTAAC)], which are present in nearly all phenylalanine-derived monolignol pathway gene promoters (Raes *et al.,* 2003). By contrast, MYB51 did not bind to any of the four SMRE-containing regions of the core GSL biosynthetic *SUR1* promoter, three of which contained AC elements (Figure 2B-2C; Supplemental Figure 2B; Supplemental File 1). Since no indole GSL biosynthetic regulator has been shown to directly target core GSL biosynthetic genes (Gigolashvili *et al.,* 2008), our results suggest that MYB51 binds to proximal SMRE-containing promoter regions to directly activate *CYP79B2, CYP79B3* and *CYP83B1* expression for 4M-I3M biosynthesis in ETI.

### Regulation of camalexin and ICN flux by MYB51/MYB122

Metabolic flux though the indole GSL pathway is primarily controlled by enzyme activities at the CYP79B2/CYP79B3-catalyzed step, consistent with theoretical predictions of flux control by the first biosynthetic step (Mikkelsen *et al.,* 2000; Zhao *et al.,* 2002; Sugawara *et al.,* 2009; Wright and Rausher, 2010). In addition, flux though the indole GSL pathway is also likely regulated by SG12-type R2R3-MYBs through changes in *CYP79B2/CYP79B3* gene expression (Celenza *et al.,* 2005; Gigolashvili *et al.,* 2007a; Frerigmann *et al.,* 2014a). Recently, MYB51 and MYB122 have been shown to also regulate camalexin biosynthesis in response to flg22 and UV stress (Frerigmann *et al.,* 2015, 2016). Since MYB51 can directly activate *CYP79B2* and *CYP79B3* promoters in response to *Psta* (Figure 3), we hypothesized that MYB51 and MYB122 may also regulate flux through the camalexin and 4OH-ICN pathways through changes in *CYP79B2/CYP79B3* gene expression. To test this hypothesis, we compared the host metabolic and transcriptional response to *Psta* in WT, *myb51, myb122, myb51 myb122*, and *myb51 myb122/DEX:MYB51-myc*. Camalexin levels were largely unchanged in *myb51* and *myb122* single mutants relative to WT, and reduced in the *myb51 myb122* double mutant (Figure 4A-4B). Similarly, the level of ICN, the immediate precursor to 4OH-ICN, was unchanged in single mutants relative to WT, and nearly abolished in the double mutant, comparable to the ETI-deficient *rpm1* mutant (Figure 4A-4B).

**Figure 3.**
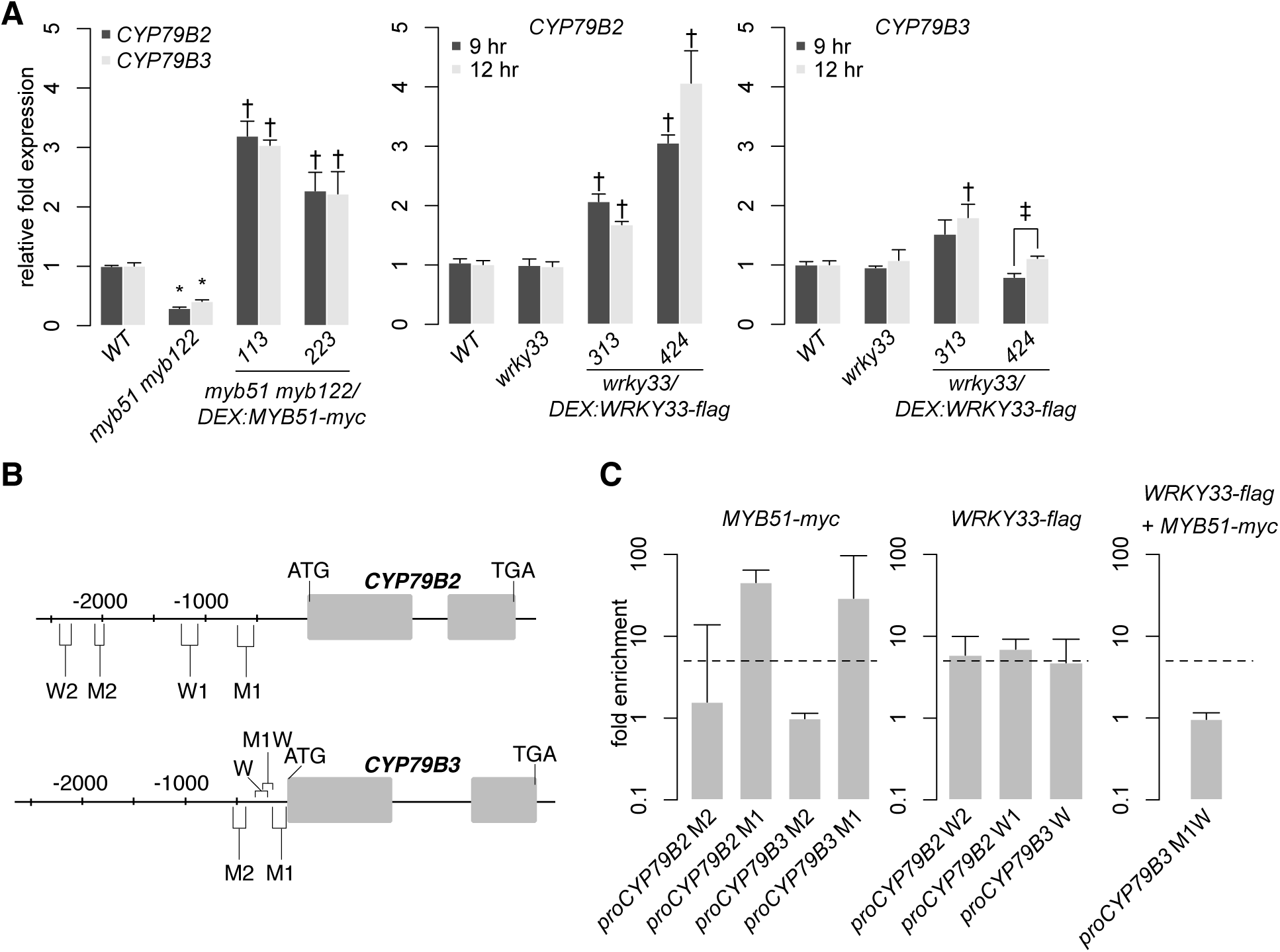
MYB51 and WRKY33 directly activate SMRE- and W-box-containing *CYP79B2* and *CYP79B3* promoters. **(A)** qPCR analysis of *CYP79B2,* and *CYP79B3* in 9-day-old seedlings co-elicited with 20 μM Dex and *Psta* for 12 hr (left) or for 9 and 12 hr (middle, right). Data represent the mean ± SE of four replicates of 13-17 seedlings each. Expression values were normalized to that of the housekeeping gene *EIF4A1* and relative to those of 9-hr elicited WT plants. Asterisks and single daggers denote statistically significant differences compared to wild-type and *wrky33* (A) or *myb51 myb122* (B), respectively and double daggers denote statistically significant differences between 9 and 12 hr time points (*P* < 0.05, two-tailed *t* test).
**(B-C)** Nucleotide positions **(B)**, ChIP-PCR analysis **(C**, left and middle), and sequential ChIP-PCR analysis **(C**, right) of SMRE (M), W-box (W), and SMRE and W-box (MW)-containing promoter (pro) regions bound by MYB51-myc and/or WRKY33-flag in 9-day-old seedlings co-elicited with 20 μM Dex or mock solution (0.5% DMSO) and *Psta* for 9 hr. Dashed lines in **(C)** represent the 5-fold cutoff between weak and strong TF-DNA interactions. Data in **(C)** represent the median ± SE of 3 (right) and 4 (left and middle) biological replicates, each containing approximately 200 seedlings. Fold enrichment was determined by calculating the ratio of PCR product intensities in ChIP Dex/Mock to Input Dex/Mock. Experiments were performed twice, producing similar results.

**Figure 4.**
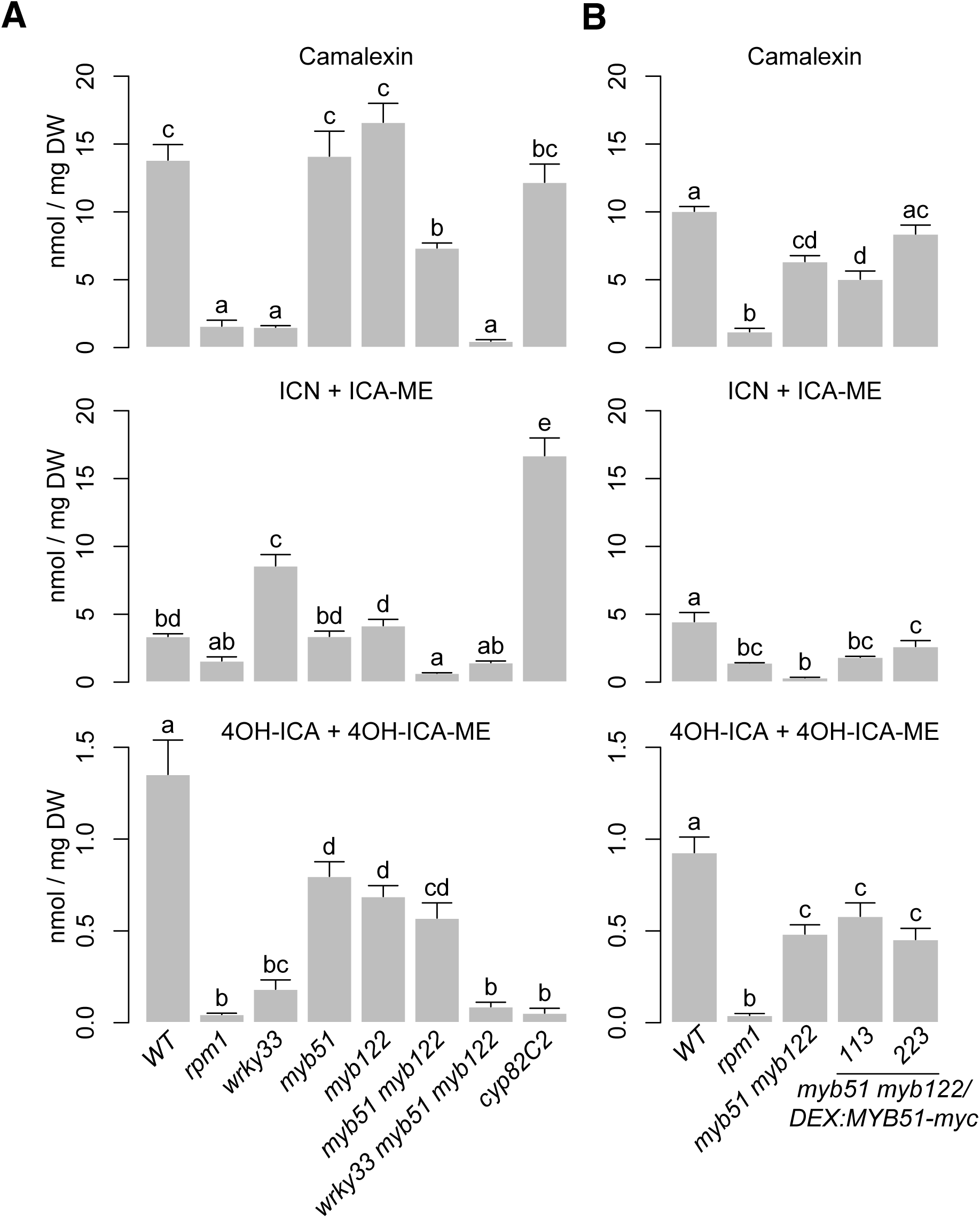
MYB51, MYB122 and WRKY33 regulate metabolic flux through camalexin and 4OH-ICN pathways. **(A)** and **(B)** LC-DAD analysis of camalexin (top), ICN (center), and 4OH-ICN (bottom) in 9-day-old seedlings elicited with *Psta* (**A**) or co-elicited with 20 μM Dex and *Psta* (**B**) for 24 hr. Data represent mean ± SE of four replicates of 13-17 seedlings each. The *rpm1* mutant is ETI-deficient when elicited with *Psta* (Bisgrove *et al.,* 1994; Barco *et al*., 2019b). The *cyp82C2* mutant is impaired in 4-hydroxylation of ICN (Rajniak *et al*., 2015). DW, dry weight. Different letters denote statistically significant differences (*P* < 0.05, one-factor ANOVA coupled to Tukey’s test). Experiments in (**A**) were performed twice, producing similar results. Experiments in (**B**) were performed three times, producing a range of results (Supplemental Figure 4), the average outcome of which is shown.

Interestingly, impairments in 4OH-ICN were observed in both *myb51* and *myb122* single and double mutants (Figure 4A-4B). Induced expression of *MYB51-myc* in at least one line restored ICN biosynthesis on average to WT levels relative to the *myb51 myb122* background (Figure 4B), By contrast, camalexin and 4OH-ICN levels were on average not modulated by induction of *MYB51-myc*.

Interestingly, a wide range of responses were observed with respect to ICN and 4OH-ICN induction in *MYB51-myc* lines (Figure 4B, Supplementary Figure 4). Therefore we further investigated the effect of MYB51-myc on expression of genes downstream from *CYP79B2*/*CYP79B3*. In agreement with the reported inability of MYB51 and MYB122 to *trans*-activate the promoters of camalexin biosynthetic genes *CYP71A13* and *CYP71B15* (also known as *PAD3*) (Frerigmann *et al.,* 2015), transcript levels of *CYP71A13* and *CYP71B15* as well as 4OH-ICN biosynthetic genes *CYP71A12* and *FOX1* were repeatedly unchanged or slightly elevated in *myb51 myb122* relative to WT (Supplemental Figure 5). Furthermore, induced expression of *MYB51-myc* had no effect on *CYP71A12* and *CYP71A13* expression and increased *CYP71B15* and *FOX1* expression in *myb51 myb122* by only 1.5-fold (Supplemental Figure 5). These results suggest that in ETI, MYB51 and MYB122 regulate biosynthetic flux to camalexin, ICN, and 4OH-ICN through gene expression changes to *CYP79B2/CYP79B3* and not *CYP71A13, CYP71B15, FOX1,* or *CYP71B15*.

### Flux regulation of camalexin and 4OH-ICN pathways by WRKY33

The transcription factor WRKY33 was recently shown to be a major regulator of camalexin and 4OH-ICN biosynthesis in ETI (Qiu *et al.,* 2008; Barco *et al.,* 2019b), directly activating nearly all associated biosynthetic genes pathways in response to *Psta* (Figure 3A) (Barco *et al.,* 2019b). To determine WRKY33’s contribution to flux regulation of camalexin and 4OH-ICN pathways in ETI, we compared host metabolic responses to *Psta* in *wrky33*, *myb51 myb122*, and the newly generated *wrky33 myb51 myb122* triple mutant (Supplemental Figure 1A). Further reductions in indole glucosinolate levels were not observed in the *wrky33 myb51 myb122* mutant relative to *myb51 myb122* (Figure 1C), indicating WRKY33 does not contribute towards flux regulation of 4M-I3M biosynthesis in ETI. In contrast, (4OH-)ICN and camalexin levels, which were reduced to varying degrees in *myb51 myb122* and *wrky33,* were collectively abolished in the *wrky33 myb51 myb122* mutant to levels comparable to the ETI-deficient *rpm1* mutant (Figure 4A). These results suggest that WRKY33 also contributes to flux regulation of the camalexin and 4OH-ICN pathways in ETI at the point of CYP79B2/CYP79B3 catalysis.

### WRKY33 directly activates W-box-containing *CYP79B2* and *CYP79B3* promoters

MYB51 is necessary and sufficient for *Psta*-induced 4OH-I3M and 4M-I3M biosynthesis (Figure 1C-D), whereas WRKY33 appears to have no role in their synthesis (Figure 1C). On the other hand, WRKY33 is necessary and sufficient for *Psta*-induced camalexin and 4OH-ICN biosynthesis (Figure 4A) (Barco *et al.,* 2019b), and MYB51 has a supporting role in their synthesis (Figure 4A-4B). MYB51 is also necessary and sufficient for *Psta*-induced *CYP79B2* and *CYP79B3* expression (Figure 3A). Since all Trp-derived defense metabolites require *CYP79B2* and/or *CYP79B3* for their synthesis, it is likely that WRKY33, like MYB51, also directly activates *CYP79B2* and *CYP79B3* expression in ETI. To test this, we first compared *Psta*-induced *CYP79B2* and *CYP79B3* expression in WT, *wrky33,* and previously characterized *wrky33/DEX:WRKY33-flag* lines (Barco *et al.,* 2019b), which express in the *wrky33* mutant background a Dex-inducible wild-type copy of *WRKY33* with a C-terminal fusion to FLAG tag. *CYP79B2* and *CYP79B3* expression in *wrky33* was mostly unchanged and occasionally reduced compared to WT in response to *Psta* (Figure 3A), but increased 2- to 4-fold upon induced expression of *WRKY33-flag* (Figure 3A). This result is consistent with a previous report of unchanged *CYP79B2* and reduced *CYP79B3* expression in *wrky33* in response to the fungal pathogen *Botrytis cinerea* (Liu *et al.,* 2015) and indicates that although WRKY33 activates *CYP79B2* and *CYP79B3* expression under ETI, MYB51 is the predominant player.

WRKY TFs specifically bind to W-box core sequences [TTGAC(T/C)] (Rushton *et al.,* 2010), and WRKY33 preferentially binds W-boxes that are within 500 nt of the ‘WRKY33-specific’ motif [(T/G)TTGAAT]) (Liu *et al.,* 2015). WRKY33 has been previously shown to bind to the distal W-box-containing promoter region of *CYP79B2* (W2 in Figure 3B) in response to flg22 (Birkenbihl *et al.,* 2017). To test whether WRKY33 binds *CYP79B2* or *CYP79B3* under ETI, we performed ChIP on 9-hr *Psta*-infected mock and Dex-treated *wrky33/DEX:WRKY33-flag* seedlings using antibodies specific to FLAG (Barco *et al.,* 2019b), and PCR-amplified W-box/WRKY33 motif-containing regions within 2500 nt upstream of the *CYP79B2* and *CYP79B3* TSS. WRKY33 bound moderately well (approximately 5-fold enrichment) to two W-box/WRKY33 motif-containing promoter regions of *CYP79B2* (W1 and W2) and *CYP79B3* (W) genes (Figure 3C; Supplemental Figure 3A-B), including the previously reported W2 region (Birkenbihl *et al.,* 2017). Since WRKY33 does not contribute to 4M-I3M biosynthesis (Figure 1C), these results suggest that WRKY33 directly activates *CYP79B2* and *CYP79B3* expression to increase metabolic flux to camalexin and 4OH-ICN biosynthetic pathways in ETI.

### MYB51 directly represses SMRE-containing *CYP82C2* promoter

SG12-type R2R3-MYBs have thus far been characterized as transcriptional activators. For example, MYB51 dramatically increases IAOx flux to indole GLSs (Figures 1C-1D) by direct activation of *CYP79B2* and *CYP79B3* expression (Figure 3; Supplemental Figure 3A). However, induced expression of *MYB51-myc* leads to variable effects on (4OH-)ICN biosynthesis in *myb51 myb122* (Figure 4B, Supplemental Figure 4). This result led us to hypothesize that additional complex flux regulation of the 4OH-ICN pathway may exist. Consistent with our hypothesis, the *CYP82C2* gene, which encodes an enzyme responsible for the synthesis of 4OH-ICN from ICN (Rajniak *et al.,* 2015), was upregulated 3.5-fold in *myb51 myb122* relative to WT in response to *Psta* (Figure 5A). Moreover, induced expression of *MYB51-myc* in *myb51 myb122* decreased *CYP82C2* expression to WT levels (Figure 5A), indicating MYB51 represses *CYP82C2* expression. We also observed that MYB51 binds strongly (>10-fold enrichment) to two SMRE-containing promoter regions of *CYP82C2* (M1 and MW) (Figure 5B-5C; Supplemental Figure 6), indicating *CYP82C2* repression by MYB51 is direct. These results suggest that MYB51 is a dual functional regulator of the 4OH-ICN pathway, directly activating *CYP79B2* and *CYP79B3* expression to increase flux of IAOx to ICN, and directly repressing *CYP82C2* expression to decrease flux of ICN to 4OH-ICN.

**Figure 5.**
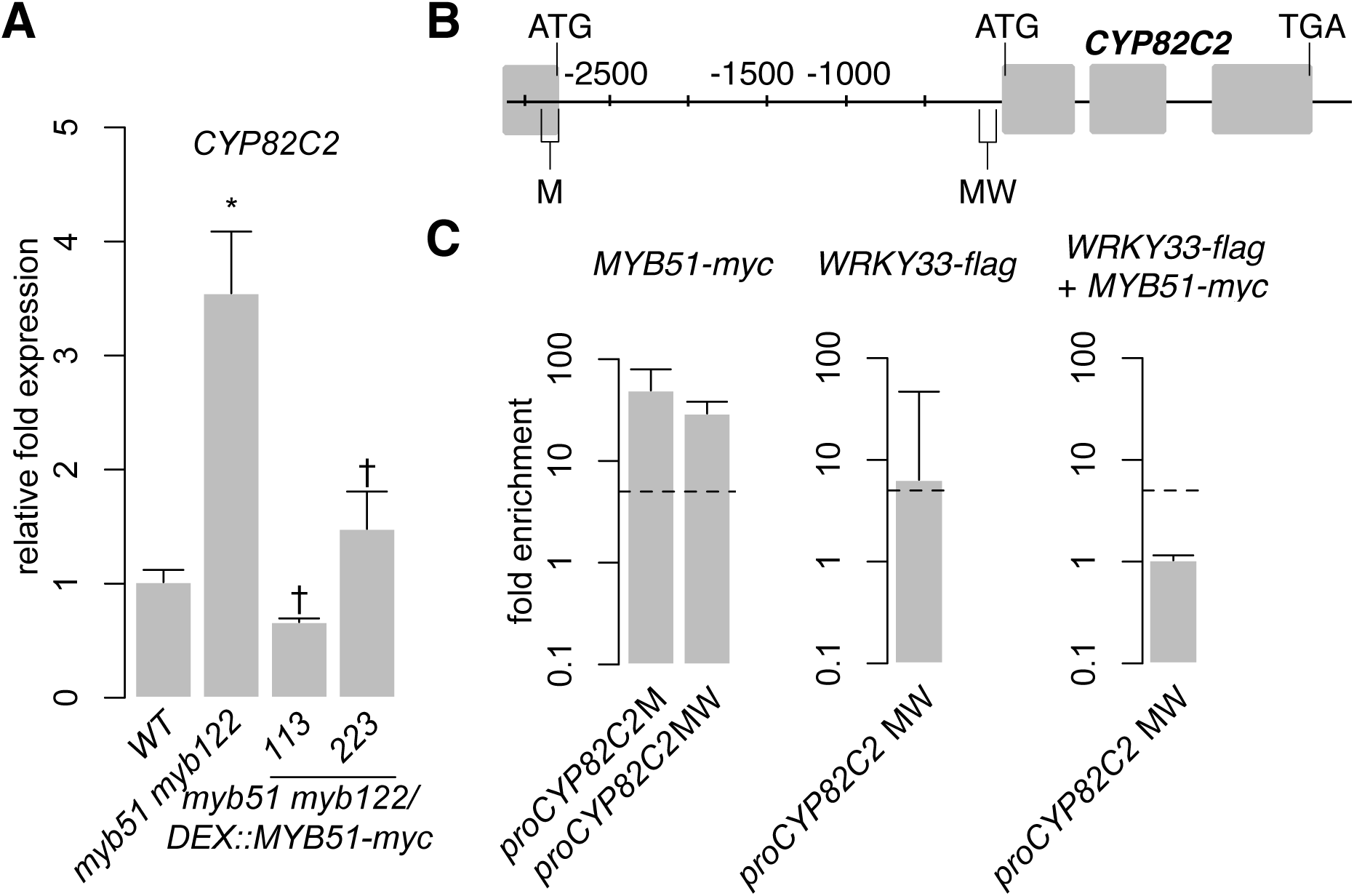
MYB51 directly represses SMRE-containing *CYP82C2* promoter. **(A)** qPCR analysis of *CYP82C2* in 9-day-old seedlings co-elicited with 20 μM Dex and *Psta* for 12 hr. Data represent the mean ± SE of four replicates of 13-17 seedlings each. Expression values were normalized to that of the housekeeping gene *EIF4A1* and relative to those of WT plants. Asterisks and daggers denote statistically significant differences compared to wild-type and *myb51 myb122*, respectively (*P* < 0.05, two-tailed *t* test). Experiment was performed twice, producing similar results.
**(B-C)** Nucleotide positions **(B)**, ChIP-PCR analysis (**C**, left and middle), and sequential ChIP-PCR analysis (**C**, right) of SMRE (M), W-box (W), and SMRE and W-box (MW)-containing *CYP82C2* promoter regions bound by MYB51-myc and/or WRKY33-flag in 9-day-old seedlings co-elicited with 20 μM Dex or mock solution (0.5% DMSO) and *Psta* for 9 hr. Dashed lines represent the 5-fold cutoff between weak and strong TF-DNA interactions. Data in **(C)** represent the median ± SE of 3 (right) and 4 (left and middle) biological replicates, each containing approximately 200 seedlings. Experiments in **(A,C)** were performed twice, producing similar results.

### WRKY33 and MYB51 do not co-localize on *CYP79B3* and *CYP82C2* promoters

In the course of mapping the *in vivo* binding sites of WRKY33 and MYB51, we observed two overlapping localization patterns. The first one involves the WRKY33-bound W and the MYB51-bound M1 promoter regions of *CYP79B3* (Figure 3B). A WRKY33-specific motif (TTTGAAT) in the WRKY33-bound W region is 50-nt from a SMRE-2 motif (ACCAACT) in the MYB51-bound M1 region (Supplemental File 1). The second involves the *CYP82C2* promoter region MW (Figure 5B), which is bound strongly by MYB51 in *myb51 myb122/DEX:MYB51-myc* plants (Figure 5C; Supplemental Figure 5A) and bound moderately (6.4-fold enrichment) by WRKY33 in *wrky33/DEX:WRKY33*-flag plants (Figure 5C; Supplemental Figure 6A). The WRKY33 and MYB51-bound MW region contains a SMRE-3 motif (ACCAAAC) that is 113-nt from one of two W-boxes (TTGACC) (Supplemental File 1). These observations suggest that WRKY33 and MYB51 could form a transcriptional complex in response to ETI-eliciting pathogens. To determine whether ETI triggers co-localization of WRKY33 and MYB51 at the same *CYP79B3* and *CYP82C2* promoter regions, we performed sequential ChIP-PCR on 9-hr *Psta*-infected, mock and Dex-treated seedlings containing both *DEX:MYB51-myc* and *DEX:WRKY33-flag* transgenes (Supplemental Figure 3C). We observed enrichment of neither the *CYP79B3* region M1W (encompassing SMRE-1 in M1 and WRKY33 motif in W) (Figure 3B, 3D; Supplemental Figure 3D) nor the *CYP82C2* region M1 amplicons from sequential ChIP of MYB51-myc followed by WRKY33-flag (Figure 5B-5C; Supplemental Figure 6B). These results indicate that ETI does not trigger stable co-localization of MYB51 and WRKY33 at *CYP79B3* and *CYP82C2* promoter regions, and suggest that WRKY33 and MYB51 likely alternate in binding to the *CYP79B3* M1W and *CYP82C2* M1 promoter regions. However it is also possible that due to lower yields of immunoprecipitated chromatin from the second IP compared to the first IP (Mendoza-Parra *et al*., 2012), transient or weak interactions such as those derived through competitive binding could be missed with this methodology.

### WRKY33 and MYB51 form a hierarchical TF cascade to control Trp-derived defense metabolism

We observed two overlapping regulatory functions of WRKY33 and MYB51; both TFs activate *CYP79B2/CYP79B3* in response to *Psta* (Figure 3A), while WRKY33 activates and MYB51 represses *CYP82C2* expression (Figure 5A) (Barco *et al.,* 2019b). The overlapping regulatory functions of WRKY33 and MYB51 suggest that these TFs form a regulatory hierarchy in response to ETI-eliciting pathogens. To determine whether ETI triggers a hierarchical regulatory interaction between WRKY33 and MYB51, we compared *Psta*-induced *WRKY33* and *MYB51* expression in WT, *myb51 myb122*, *myb51 myb122/DEX:MYB51-myc*, *wrky33*, and *wrky33/DEX:WRKY33-flag* plants. *WRKY33* expression was modestly increased in *myb51 myb122* relative to WT, and further increased in at least one *MYB51-myc* line post-elicitation with *Psta* (Figure 6A). In addition, these increases in *MYB51-myc* did not rise proportionally with the 4.5- and 50-fold increases in *MYB51-myc* expression (Figure 2A), but instead peaked at ∼ 2.5-fold relative to WT level (Figure 6A). These results indicate that MYB51 does not directly regulate *WRKY33* expression in ETI. By contrast, *MYB51* expression was reduced in *wrky33* relative to WT, and restored to greater than WT level upon induced expression of *WRKY33-flag* at both 9 and 12 hr post-elicitation (Figure 6A). Furthermore, the fold increases in *MYB51* expression in *wrky33/DEX:WRKY33-flag* and in similar transgenics expressing a C-terminal c-myc epitope were proportional to the previously reported fold increases in *WRKY33* expression (Figure 6A) (Barco *et al*., 2019b). These results indicate that WRKY33 is necessary and sufficient to activate *MYB51* expression in ETI.

**Figure 6.**
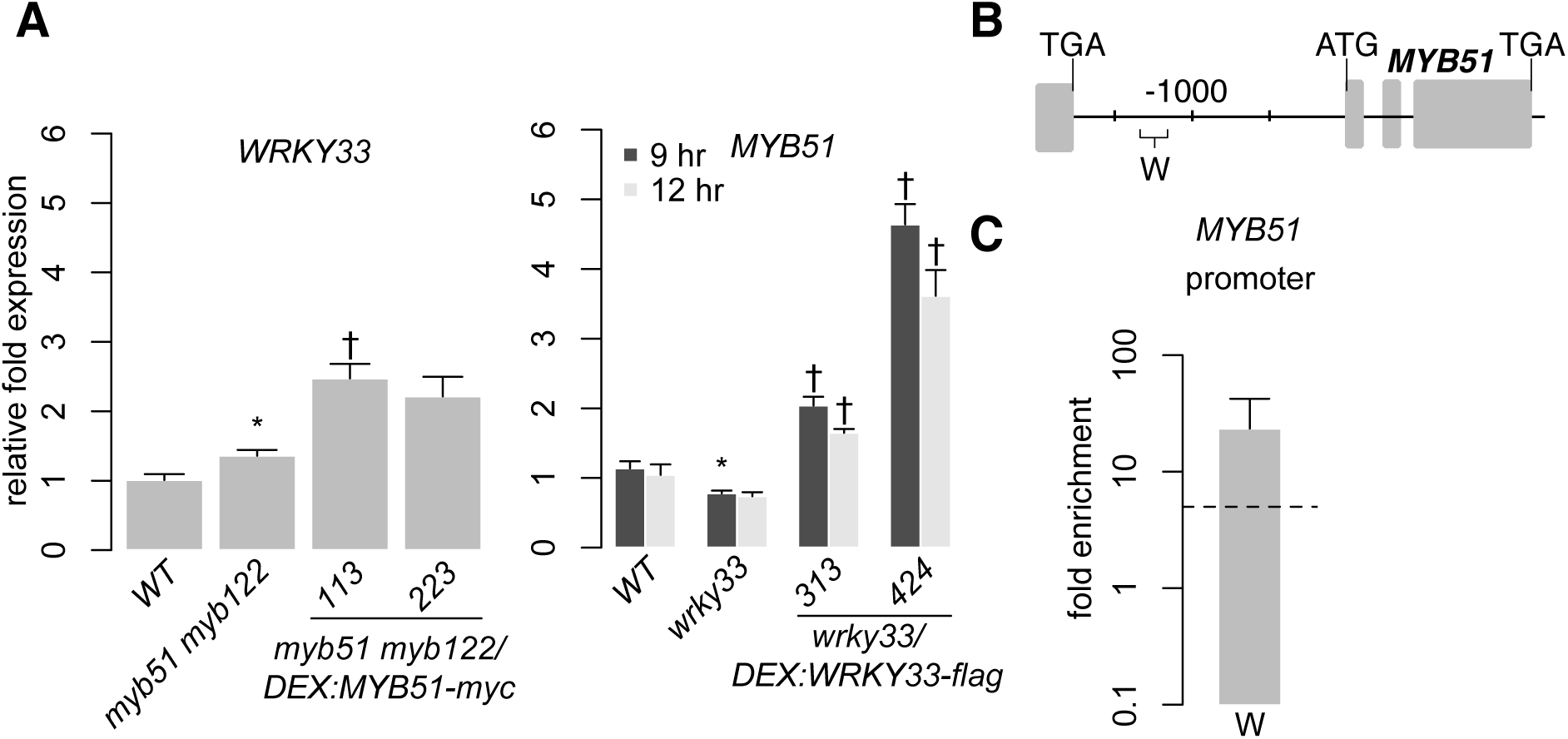
WRKY33 directly activates W-box-containing *MYB51* promoter. **(A)** qPCR analysis of *WRKY33* (left) and *MYB51* (right) in 9-day-old seedlings co-elicited with 20 μM Dex and *Psta* for 12 hr (left) or 9 and 12 hr (right). Data represent the mean ± SE of four replicates of 13-17 seedlings each. Expression values were normalized to that of the housekeeping gene *EIF4A1* and relative to those of WT plants. Asterisks and daggers denote statistically significant differences compared to wild-type and *myb51 myb122* (left) or *wrky33* (right) respectively (*P* < 0.05, two-tailed *t* test). Experiments were performed at least twice including in WRKY33-myc (Barco et al., 2019b), producing similar results.
**(B)** and **(C)** Nucleotide positions **(B)** and ChIP-PCR analysis **(C)** of W-box-containing *MYB51* promoter region W bound by WRKY33-flag in *wrky33/DEX:WRKY33-flag* plants co-elicited with 20 μM Dex or mock solution (0.5% DMSO) and *Psta* for 9 hr. The *WRKY33* promoter lacks SMRE motifs. Dashed line represents the 5-fold cutoff between weak and strong TF-DNA interactions. Data in **(C)** represent median ± SE of 4 biological replicates, each containing approximately 200 seedlings. Experiment was performed twice, producing similar results.

We then performed ChIP-PCR analysis in *wrky33/DEX:WRKY33-flag* lines of the *MYB51* promoter region W, which contains a 50-nt stretch of three W-boxes (Figure 6B; Supplemental File 1). We observed strong WRKY33 binding (>10-fold enrichment) in response to *Psta* (Figure 6C; Supplemental Figure 6), indicating that WRKY33 directly activates the *MYB51* promoter in ETI. This finding is consistent with a previous report of WRKY33 interaction with (but non-regulation of) the *MYB51* locus in response to the fungal pathogen *Botrytis cinerea* (Liu *et al.,* 2015). Since WRKY33 does not contribute to 4M-I3M biosynthesis (Figure 1C), these results indicate that a hierarchical TF cascade regulates camalexin and 4OH-ICN biosynthesis in ETI.

### *CYP79B2* and *CYP79B3* display coherent feedforward loop connectivity to WRKY33 and MYB51

The regulatory interactions between WRKY33, MYB51, and *CYP79B2* (and *CYP79B3*) resemble those of a coherent type 1 FFL circuit (C1-FFL) with OR-gate logic (Figure 7A) (Mangan *et al*., 2003a; Alon, 2007), in which WRKY33 activates the target genes *CYP79B2* and *CYP79B3* as well as their activator *MYB51,* and either WRKY33 or MYB51 is sufficient to directly activate *CYP79B2* and *CYP79B3* expression in response to *Psta* (Figures 3, 6; Supplemental Figures 3, 7). The presence of a second transcriptional activator (MYB51) in the indirect regulatory path from the first activator (WRKY33) to the target gene is responsible for a built-in time delay between when the offset signal from the direct path and the offset signal from the indirect path arrive at the target gene, resulting in delayed inactivation (continued activation) of target gene response at the offset of WRKY33 activity (Figure 7A) (Mangan *et al*., 2003a; Kalir *et al.,* 2005).

**Figure 7.**
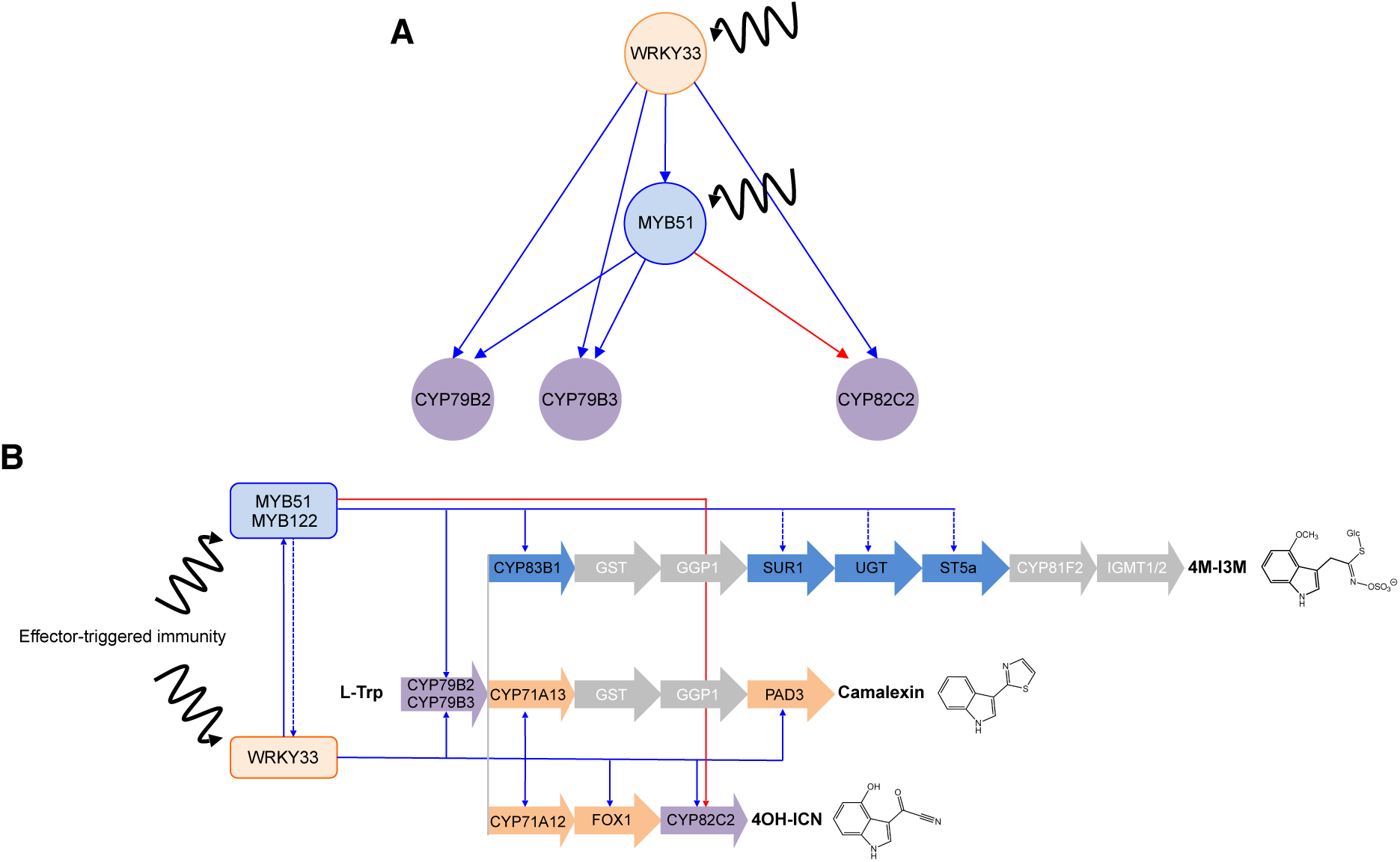
Interlinked coherent and incoherent type 1 feed-forward loops in ETI-responsive specialized metabolism. **(A-B)** Three FFLs identified in this study form a composite hierarchical network motif **(A)** that is likely responsible for the expression dynamics and metabolic flux changes through the camalexin and 4OH-ICN pathways **(B).** Blue and red directed lines indicate activating and inhibitory direct (solid) and indirect (dashed) interactions, respectively, between TFs (orange and blue circles in **A**, rectangles in **B**) and pathway genes (purple circles in **A**, arrows in **B**).

To confirm the hierarchical TF cascade and coherent configuration of the FFL, we compared the temporal dynamics of *MYB51*, *CYP79B2* and *CYP79B3* expression in two independent lines of *wrky33/DEX:WRKY33-flag* in response to induced expression of *WRKY33-flag* at 9 and 12 hours post-elicitation (Barco *et al*., 2019b). As WRKY33 expression decreased during that time period (Barco et al., 2019b), *MYB51* and *CYP79B2* expression remained steady (Figures 3A, 6A), and *CYP79B3* expression increased in one line (Figure 3A). This result indicates continued activation of *CYP79B2* and *CYP79B3* expression in the face of decreasing WRKY33 activity, and confirms their C1-FFL connectivity to WRKY33 and MYB51 in ETI.

### *CYP82C2* displays incoherent feedforward loop connectivity to WRKY33 and MYB51

The regulatory interactions between WRKY33, MYB51, and *CYP82C2* resemble those of an incoherent type 1 FFL circuit (I1-FFL) with OR-gate logic (Figure 7A) (Milo *et al.,* 2002; Mangan *et al*., 2003a), in which WRKY33 activates both the target *CYP82C2* and its repressor *MYB51,* and either WRKY33 or MYB51 is sufficient to directly regulate *CYP82C2* expression in response to *Psta* (Figures 5-6; Supplemental Figures 6-7). The I1-FFL motif was shown to have, among other dynamical functions, the ability to produce a non-monotonic (first increasing and then decreasing) target gene response to increasing expression of the activator TF (Basu *et al.,* 2005; Entus *et al.,* 2007; Kaplan *et al.,* 2008). A necessary condition to this non-monotonic behavior (i.e. a concentration-dependent, bell-shaped expression pattern) is that the repressor TF promoter remains responsive to a high level of activator TF activity, regardless of whether the activator and repressor can simultaneously bind the target promoter region (Kaplan *et al.,* 2008).

To confirm the hierarchical TF cascade and incoherent configuration of the FFL, we compared *MYB51* and *CYP82C2* expression between WT and *wrky33/DEX:WRKY33-flag*. *MYB51* expression exceeded WT levels by greater than 2-fold, and was proportional to fold increases in *WRKY33* expression reported in Barco *et al*. (2019b) (Figure 6A), indicating that the *MYB51* promoter responds monotonically to high (greater than WT) levels of WRKY33 activity. By contrast, *CYP82C2* expression was restored to WT levels by 9 hr post-elicitation with *Psta* and either remained steady or decreased to below WT levels by 12 hr post-elicitation, and thus was not proportional to fold increases in *WRKY33* expression at both time points reported in Barco *et al*. (2019b) (Figure 5A). These results indicate that the *CYP82C2* promoter responds non-monotonically to high (greater than WT) level of WRKY33 activity, as predicted by its I1-FFL connectivity to WRKY33 and MYB51 in ETI.

### Predominant role of camalexin and 4OH-ICN in bacterial resistance

Camalexin and 4OH-ICN have been shown to contribute non-redundantly to basal immunity against *Pst*, with WRKY33 as a major regulator (Figure 4) (Qiu *et al.,* 2008; Rajniak *et al.,* 2015; Barco *et al.,* 2019b). 4M-I3M also contributes to basal immunity against *Pst*, with MYB51/MYB122 as major regulators (Figure 1) (Clay *et al.,* 2009). To determine the extent of redundancy between 4M-I3M and camalexin/4OH-ICN in bacterial resistance, we used the TF mutants described in this study as proxies for mutants in defense-induced camalexin/4OH-ICN and 4M-I3M biosynthesis. Specifically, we compared bacterial growth of *Pst* in adult leaves of WT, *wrky33, myb51 myb122*, and *wrky33 myb51 myb122.* Consistent with previous reports, surface-inoculated leaves of *wrky33* and the 4OH-ICN biosynthetic mutant *cyp82C2* mutant showed increased susceptibility to *Pst* relative to WT (Figure 8) (Rajniak *et al.,* 2015; Barco *et al.,* 2019b). The *wrky33 myb51 myb122* mutant also showed increased susceptibility to *Pst* relative to WT and comparable to *wrky33* and *cyp82C2* (Figure 8). By contrast, the bacterial resistance of *myb51 myb122* was intermediate between WT and *wrky33*, such that it was not significantly different from either line (Figure 8). These results suggest a predominant role of camalexin and 4OH-ICN in bacterial resistance.

**Figure 8.**
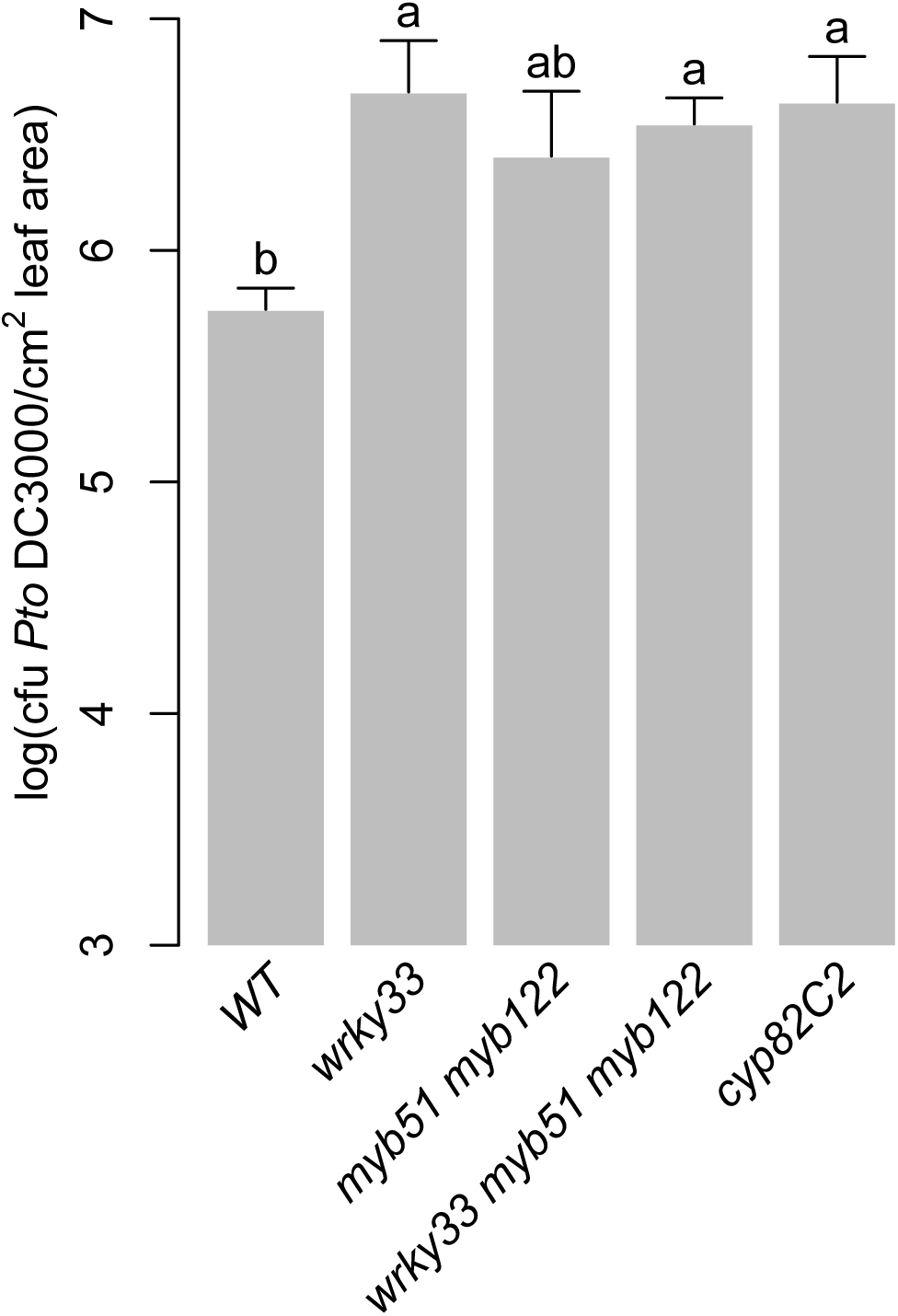
Predominant role of camalexin and 4OH-ICN metabolism in bacterial resistance. Bacterial growth analysis of *Pst* in surface-inoculated five-week-old leaves. Data represent the mean ± SE of 8-12 biological replicates with one leaf each. Different letters denote statistically significant differences (*P* < 0.05, one-factor ANOVA coupled to Tukey’s test). CFU, colony-forming units. Experiments were performed at least twice, producing similar results.

## Discussion

A signaling node between cell surface receptors and genes underlying specialized metabolism, TFs are also core components of network motifs, whose topological features are often independent from detailed reaction mechanisms and kinetic parameters (Alon, 2007; Gutenkunste *et al.,* 2007) and thus can be easily tuned or adjusted to produce the desired expression dynamics and metabolic output. Despite successes in mapping and modeling GRNs for a wide variety of model systems, only a handful of case studies have linked network architecture in its natural context to its dynamic performance and physiological relevance. Here, we identified two C1-FFLs and one I1-FFL that form a composite hierarchical transcription network motif, with WRKY33 as the condition-dependent master regulator and MYB51 as the dual functional regulator (Figure 7A). This composite motif is likely responsible for the observed gene expression dynamics and subsequent metabolic fluxes through the CYP79B2/CYP79B3- and CYP82C2-catalyzed steps in the camalexin and/or 4OH-ICN pathways in ETI (Figure 7B). One of the two motifs may have also facilitated the evolution of the 4OH-ICN pathway in *A. thaliana* via regulatory capture of the newly evolved *CYP82C2* gene into the MYB51 regulon (see below).

### Hierarchical regulatory architecture of ETI-responsive specialized metabolism

The GRNs of *E. coli, C. elegans,* human cells, and *A. thaliana* are characterized by an FFL-enriched hierarchical modularity that is predicted or demonstrated to closely overlap with known environmental stress responses or biological functions (Dobrin *et al.,* 2004; Ma *et al.,* 2004; Tang *et al.,* 2012; Defoort *et al.,* 2018). In addition, most FFLs in the simplest GRN (*E. coli*) contain global regulators, which often act together with local regulators to regulate functionally far-related genes, resulting in a network arrangement of different layers by regulatory depth, with global regulators in the top layers, target genes in the bottom layer, and FFLs spanning the layers (Ma *et al.,* 2004). Consistent with this finding, WRKY33 and MYB51 serve as the respective global and local regulators for the FFL circuits identified in this study, spanning a hierarchical regulatory architecture in the transcriptional network of ETI-responsive specialized metabolism. WRKY33 is characterized as a global regulator because it controls functionally diverse gene modules, such as hormone homeostasis in response to *B. cinerea* (Liu *et al.,* 2015), camalexin and 4OH-ICN biosynthesis in response to flg22, *Psta*, *B. cinerea* and/or *Pst* (Figure 4A) (Qiu *et al.,* 2008; Birkenbihl *et al.,* 2012, 2017; Barco *et al.,* 2019b), and immune receptor signaling in response to flg22 (Logemann *et al.,* 2013). By contrast, MYB51 is a local regulator, controlling only Trp-derived specialized metabolism in response to flg22, *Psta* and the fungal pathogen *Plectosphaerella cucumerina* (Figures 1C-1D, 4A-4B) (Clay *et al.,* 2009; Frerigmann *et al.,* 2016). FFLs in the transcription networks for lignocellulose metabolism in *A. thaliana* and phenolic metabolism in maize have been shown to link developmental regulators to metabolic regulators and their target biosynthetic genes (Zhong and Ye, 2012; Yang *et al.,* 2017), but to our knowledge, the FFLs identified in this study are the first to link two metabolic regulators in plant specialized metabolism.

Compared to development, defense-responsive metabolism has fewer, if any, irreversible regulatory switches, and a greater potential for fine-tuning through rapid interconversion between compounds (Samal and Jain, 2008; Chandrasekaran and Price, 2010; Watson and Walhout, 2014), thus calling into question on the necessity of a hierarchical regulatory architecture or fine-tuning through gene expression for defense-responsive metabolism (Li *et al.,* 2014). However, metabolic fluxes through a pathway are often regulated in response to external perturbations (Fell, 1997) through changes in the activities of its enzymes via gene expression and/or covalent modification (termed hierarchical regulation), and through changes in their interactions with other enzymes via allosteric regulation and/or (co-)substrate saturation (termed metabolic regulation) (ter Kuile and Westerhoff, 2001; Rossell *et al.,* 2005). Hierarchical regulation through proportionate increases in relevant gene expression has been demonstrated for enzymes of central carbon metabolism in response to environmental stresses in single-celled organisms (ter Kuile and Westerhoff, 2001; Even *et al.,* 2003; Rossell *et al.,* 2005; Rossell *et al.,* 2006; Daran-Lapujade *et al.,* 2007). However, to our knowledge, similar studies in defense-responsive plant specialized metabolism have been limited to identifying enzymes with high flux control (Olson-Manning *et al.,* 2013). Our findings suggest significant hierarchical regulation of fluxes through the camalexin and 4OH-ICN pathways through coherent feed-forward activation of the *CYP79B2/CYP79B3* promoters (Figures 3, 6; Supplemental Figure 3; Barco *et al.,* 2019b), and incoherent feed-forward regulation of the *CYP82C2* promoter in response to *Psta* (Figures 5-6; Supplemental Figure 6A). Future research is needed to determine whether the CYP82C2 enzyme has high flux control of the 4OH-ICN pathway and to what extent metabolic fluxes through individual enzymes with high flux control are regulated by gene expression or by metabolic regulation in ETI-responsive specialized metabolism. Increased understandings of flux control and regulation would promote systems biological approaches toward engineering plant specialized metabolism (Baghalian *et al.,* 2014).

### Physiological relevance of FFLs

The three branches of ETI-responsive Trp-derived specialized metabolism are distinctly and tightly controlled, both in timing and amplitude, despite sharing the CYP79B2 and CYP79B3-catalyzed first step (Mikkelsen *et al.,* 2000; Glawischnig *et al.,* 2004; Rajniak *et al.,* 2015). For instance, pathogen defense-responsive indole GSLs have been shown previously to be present at low levels in uninfected plants and accumulate to modest levels at the expense of the parent metabolite I3M in flg22- and *Psta*-inoculated plants (Clay *et al.,* 2009; Barco *et al.,* 2019b), whereas camalexin and 4OH-ICN are absent in uninfected plants, at low-to-undetectable levels in flg22-inoculated plants, and at high levels in *Psta*-inoculated plants (Figures 1C-1D, 4A-4B) (Rajniak *et al.,* 2015; Barco *et al.,* 2019b). The dynamical functions of three FFLs (two C1-FFLs and one I1-FFL) identified in this study likely contribute to these observed physiological responses.

First, C1-FFLs are proven mechanisms for delaying response times in transcription networks (Mangan *et al*., 2003a, 2003b; Kalir *et al.,* 2005). The C1-FFL connectivity of *CYP79B2* and *CYP79B3* to WRKY33 and MYB51 (Figure 7A) is likely responsible for the continued activation of *CYP79B2* and *CYP79B3* expression despite decreasing WRKY33 activity in ETI, which in turn may account for the high production of all defense-responsive Trp-derived metabolites at the onset of ETI, and the continued production of indole GSLs at the offset of ETI. Future experiments are needed to confirm that the duration of the time delay displayed by the two clustered C1-FFLs can be fine-tuned through changes in MYB51’s biochemical parameters (for example, its activation threshold for the target gene promoters) (Alon, 2007).

Second, I1-FFLs are proven mechanisms for non-monotonic time-responses in transcription networks that mediate tradeoffs in metabolism (Weickert and Adhya, 1993; Kaplan *et al.,* 2008; Ascensao *et al.,* 2016). For example, an I1-FFL was shown to regulate the non-monotonic target gene responses of galactose metabolism so that when cells are severely starved for glucose, galactose breakdown is reduced, and galactose is redirected towards cell wall synthesis (Weickert and Adhya, 1993).

Similarly, the I1-FFL connectivity of *CYP82C2* to WRKY33 and MYB51 (Figure 7A) is likely responsible for the initial increased and then decreased activation of *CYP82C2* expression in the face of increasing WRKY33 activity in ETI, so that when WRKY33 activity is high, 4OH-ICN synthesis is reduced, and IAOx is redirected towards indole GSL biosynthesis (Figure 7B). Indole GSL biosynthesis has been shown to be metabolically linked to auxin homeostasis; severe reductions in indole GSL biosynthesis by mutagenesis has been shown to result in overflow of IAOx to indole-3-acetic acid (auxin) and ultimately to high-auxin phenotypes, such as severe defects in plant growth and development (Boerjan *et al.,* 1995; Bak and Feyereisen, 2001; Mikkelsen *et al.,* 2004). Future experiments are needed to confirm that the concentration-dependent, bell-shaped target gene expression pattern can be fine-tuned through changes in MYB51 concentration and/or MYB51’s binding affinity for the target gene promoter (Entus *et al.,* 2007).

### WRKY33 is a condition-dependent master regulator

A hierarchical regulatory architecture of ETI-responsive specialized metabolism requires that a single master or key regulator is able to be both necessary and sufficient to drive biosynthesis (Li *et al.,* 2014). Our data shows that WRKY33 fulfills that requirement; its absence results in the near-absence of camalexin and 4OH-ICN biosynthesis in *Psta*-infected plants, comparable to the ETI-deficient *rpm1* mutant (Figure 4A), and its induced expression restores camalexin and 4OH-ICN biosynthesis in the *Psta*-infected *wrky33* mutant to levels that exceed WT (Barco *et al*., 2019b). Furthermore, WRKY33 initiates a feed-forward regulation of camalexin and 4OH-ICN biosynthetic gene responses via MYB51 (Figures 3, 5-6; Supplemental Figures 3, 6-7). However, WRKY33’s ability to initiate camalexin and 4OH-ICN biosynthetic gene responses occurs only in cells undergoing ETI-specific reprogramming (Figure 4A) (Barco *et al.,* 2019b). This condition-dependent regulation is consistent with recent ChIP-Seq and expression analysis revealing that WRKY33 binds to a large number of genomic loci in *B. cinerea* or flg22-elicited plants, yet only activates or represses transcription at a subset of these genes (Liu *et al*., 2015; Birkenbihl *et al*., 2017). Condition-dependent regulation has also been demonstrated for the classic master regulator in development, MyoD, which initiates skeletal muscle-specific gene expression only in cells conditioned to be permissive to myogenesis (Aziz *et al.,* 2010). Further protein interaction studies are needed to determine mechanistically what additional interactions are needed to activate WRKY33-bound promoters. The simplest hypothesis would be that two distinct classes of chromatin proteins are recruited to WRKY33-bound promoters, and that chromatin remodeling is needed for dynamic switching of gene expression during ETI.

### MYB51 is a dual functional regulator

Few plant TFs have been identified to act as transcriptional activators or repressors, depending on DNA-binding sequences or interactions with additional co-factors. They include WUSCHEL in stem cell regulation and floral patterning, WRKY53 in leaf senescence, WRKY6 and tomato Pti4 in pathogen defense, and WRKY33 in camalexin and ABA biosynthesis (Robatzek and Somssich, 2002; Miao *et al.,* 2004; González-Lamothe *et al.,* 2008; Ikeda *et al.,* 2009; Liu *et al.,* 2015). Our data shows that MYB51 also possesses dual functionality, acting as an activator or repressor in a manner dependent on promoter context. 4OH-ICN and camalexin profiles (Figure 4A-4B, Supplemental Figure 4) and *CYP79B2, CYP79B3* and *CYP82C2* transcript profiles (Figures 3A, 5A) in *MYB51* gain or loss of function lines – especially the variability in metabolism observed in *myb51 myb122 /MYB51-myc* – indicate complex regulatory control of 4OH-ICN and camalexin biosynthesis by MYB51, in the former case at both the first and last steps of biosynthesis (Figure 7B). By contrast, 4M-I3M metabolite and *CYP83B1* and *SUR1* transcript profiles in *myb51 myb122* and *myb51 myb122/DEX:MYB51-myc* indicate straightforward positive regulation of indole GLS biosynthesis by MYB51 (Figure 7B). Further protein interaction studies are needed to determine mechanistically how MYB51 exerts its dual regulatory functions. The simplest hypothesis would be that MYB51 is recruited to distinct repressor and activator complexes at defined promoter sites.

### Regulatory capture of newly duplicated gene *CYP82C2* into the MYB51 regulon

Recent phylogenetic decomposition analysis of the *A. thaliana* GRN indicated that novel genes are more likely to be regulated by conserved TFs in FFLs, as well as attach to gene modules with specific biological functions, instead of forming modules on their own (Defoort *et al.,* 2018). Few newly evolved plant specialized metabolic pathways have been shown to be regulated by existing TFs. They include the *A. thaliana-*specific benzoyloxy-GSL pathway and the Brassicaceae-specific SG25-type R2R3-MYBs MYB115 and MYB118 (Zhang *et al.,* 2015; Barco *et al.,* 2019a), the Brassicales-specific core GSL pathway and the plant lineage-specific SG3e-type MYCs MYC2–5 (Schweizer *et al.,* 2013; Frerigmann *et al.,* 2014b; Chezem *et al.,* 2016;), and the Brassicaceae-specific camalexin and *A. thaliana*-specific 4OH-ICN pathways and the plant lineage- specific WRKY33 (Bednarek *et al.,* 2011; Rinerson *et al.,* 2015; Schluttenhofer and Yuan, 2015; Barco *et al.,* 2019b). *MYB51/MYB122* and *CYP79B2/CYP79B3* are unique to the GSL-synthesizing plant order Brassicales (Fahey *et al.,* 2001; Bekaert *et al.,* 2012; Barco *et al.,* 2019a), whereas *CYP82C2* is a newly duplicated enzyme gene in the *A. thaliana*-specific 4OH-ICN biosynthetic pathway (Rajniak *et al.,* 2015; Barco *et al*., 2019b). Our data shows that the indole GSL regulator MYB51 interacts with the *CYP82C2* promoter at the M and MW regions (Figure 5B-5C; Supplemental Figure 6A) and negatively regulates its expression (Figure 5A). Further phylogenetic and syntenic analyses are needed to determine mechanistically how the *CYP82C2* gene was recruited into the MYB51 regulon. The simplest hypothesis would be that the *CYP82C2* promoter acquired one or more MYB51-binding sites through mutation and/or transposition (Wittkopp and Kalay, 2012).

## METHODS

### Plant Materials and Growth Conditions

For transcriptional and metabolomics analyses, seeds of *Arabidopsis thaliana* accession Columbia-0 (Col-0) were surface-sterilized in 20% (v/v) bleach/0.1% (v/v) Tween-20 aqueous solution for 5 min, washed three times with sterile water, stratified at 4°C for 2 days and sown in 12-well microtiter plates sealed with Micropore tape (3M, St. Paul, MN), each well containing 15 ± 2 seeds and 1 mL of filter-sterilized 1X Murashige and Skoog (MS; Murashige & Skoog, 1962) media (pH 5.7–5.8) (4.43 g/L MS basal medium with vitamins [Phytotechnology Laboratories, Shawnee Missions, KS], 0.05% MES hydrate, 0.5% sucrose). The plates were placed on grid-like shelves over water trays on a Floralight cart, and plants were grown under long-day conditions (16-hr light cycle [70-80 μE m^-2^ s^-1^ light intensity], 21°C, 60% relative humidity). For chromatin immunoprecipitation (ChIP) analyses, approximately 200 surface-sterilized seeds were sown in a 100-x 15-mm petri plate containing 20 mL of 1X MS media. Media were refreshed on day 9 prior to bacterial elicitation. 9-day-old seedlings were inoculated with *Psta* to OD_600_ of 0.013, and seedlings and/or ∼ 1 mL media were snap-frozen in liquid nitrogen 9 hr post-infection for ChIP analyses, 12 hr post-infection for qPCR analyses, and 24-to-48 hr post-infection for LC-DAD-FLD-MS analyses, prior to –80°C storage.

For bacterial infection assays, plants were grown on soil (3:1 mix of Farfard Growing Mix 2 [Sun Gro Horticulture, Vancouver, Canada] to vermiculite [Scotts, Marysville, OH]) at 22°C daytime/18°C nighttime with 60% humidity under a 12-hr light cycle (50 [dawn/dusk] and 100 [midday] μE m^-2^ s^-1^ light intensity).

The following homozygous Col-0 mutants and T-DNA/transposon insertion lines were obtained from the Arabidopsis Biological Resource Center (ABRC): *cyp82C2* (GABI_261D12; CS425008); *myb51* (SM_3_16332; CS104159), *myb122* (also referred to as *myb122-3*, SALK_022993); *pen2* (GABI_134C04), *rpm1* (CS67956) and *wrky33* (SALK_006603).

### Plant Binary Vector Construction and Transformation

To generate the *DEX:MYB51-myc* construct, the *MYB51* coding sequence was PCR-amplified from genomic DNA using the primers MYB51gXhoF (5’-AACTCGAGATGGTGCGGACACCGTG-3’) and MYB51gStuR (5’-AAGGCCTCCAAAATAGTTATCAATTTCGTC-3’), and subcloned into the *Xho*I/*Stu*I sites of pTA7002-6x c-Myc binary vector (Aoyama and Chua, 1997; Chezem *et al*., 2017). Construct was introduced into *myb51 myb122-3* plants via *Agrobacterium tumefaciens*-mediated floral dip method (Clough & Bent, 1998), and transformants were selected on agar media containing 15 μg/mL hygromycin B (Invitrogen, Carlsbad, CA).

### Extraction and LC-DAD-FLD-MS Analysis of Glucosinolates

Glucosinolates (GSLs) were extracted and analyzed as desulfo-GSLs as described in Barco *et al*. (2019b). Briefly, extracts were separated by reversed-phase chromatography on an Ultimate 3000 HPLC (Dionex, Sunnyvale, CA) system, using a 3.5-μm, 3 x 150-mm Zorbax SB-Aq column (Agilent, Santa Clara, CA). A coupled DAD-3000RS diode array detector (Dionex), FLD-311 fluorescence detector (Dionex), and MSQPlus mass spectrometer (Dionex) collected UV absorption spectra at 229-nm, fluorescence spectra at 275/350-nm (ex/em), and ESI mass spectra in positive/negative ion modes in the range of 100-1000 m/z, respectively.

### Extraction and LC-DAD-FLD-MS Analysis of Camalexin and 4OH-ICN

Camalexin and 4OH-ICN extraction from homogenates of lyophilized tissue and media samples was performed as described in Barco *et al*. (2019b) with the following modifications. For media samples, 2.5 µL of internal standard (IS; 200 μM 4-methoxyindole [Sigma-Aldrich] in 100% methanol) was added to extract per mg dry weight of accompanying seedling tissue sample and no formic acid was added to the mobile phases during extraction. ICN/ICN-ME, 4OH-ICA/4OH-ICA-ME and camalexin were quantified as described in Barco et al. (2019b).

### RNA extraction and qPCR analysis

Total RNA extraction and qPCR were performed as described in Chezem *et al*. (2017). The Pfaffl method (Pfaffl, 2001) and calculated primer efficiencies were used to determine the relative fold increase of the target gene transcript over *EIF4A1* (*AT3G13920*) housekeeping gene transcript for each biological replicate. Primer sequences and efficiencies are listed in Supplemental Table 2.

### Total protein extraction, SDS-PAGE and western blotting

Total protein extraction was performed as previously described (Barco et al., 2019b). 5 (DEX:MYB51-myc or DEX:WRKY33-flag) or 50 µL (DEX:MYB51-myc and DEX:WRKY33-flag) of extract was loaded onto a 10% SDS-PAGE gel, and the separated proteins were transferred to PVDF membrane (Millipore, Billerica, MA), stained with Ponceau S for labeling of total protein, and probed with either FLAG M2 (Sigma-Aldrich, cat# F1804) or c-Myc 9E10 (Santa Cruz Biotechnology, cat# sc-40) antibodies diluted 1:1000 in 1X PBS containing 5% (w/v) nonfat milk.

### Chromatin immunoprecipitation (ChIP) and PCR

ChIP was performed on *myb51/DEX:MYB51-myc* and *wrky33/DEX:WRKY33-flag* nuclear extracts as described in Barco *et al*. (2019b) with the following modification. Anti-FLAG M2 Affinity Gel (Sigma-Aldrich) and EZview^TM^ Red Anti-c-Myc Affinity Gel (Sigma-Aldrich) were used to immunoprecipitate chromatin-bound MYB51-myc and WRKY33-flag, respectively.

Sequential ChIP was performed on WT/*DEX:MYB51-myc, DEX:WRKY33-flag* nuclear extracts as described in Mendoza-Parra *et al*. (2012) with modifications. Initial ChIP was performed using EZview^TM^ Red Anti-c-Myc Affinity Gel. Chromatin was immunoprecipitated first with the c-Myc antibody and second with the FLAG antibody because of stronger chromatin binding with the c-Myc antibody. To avoid false positive IP readouts at the end of the assay (a common problem with sequential ChIP assays), the first antibody is covalently crosslinked to a solid substrate (Mendoza-Parra *et al.,* 2012). Following washes, beads were incubated in TE containing 10 mM DTT for 30 min at 37°C, and the released TF-DNA complexes were extracted into ChIP dilution buffer (1% Triton X-100, 1.2 mM EDTA, 16.7 mM Tris-Cl [pH 8], 167 mM NaCl, 1x Complete EDTA-free protease inhibitor cocktail [Roche]) and immunoprecipitated a second time using FLAG M2 antibody (Sigma-Aldrich) and Protein G magnetic beads (EMD Millipore, Burlington MA) pre-treated with 0.1% (w/v) non-fat milk in 1X PBS and 0.5 mg/mL sheared salmon sperm DNA.

PCR analysis was performed on nuclear extracts prior to antibody incubation (input) and after ChIP. PCR conditions were as follows: 95°C for 3 min; 40 cycles of 95°C for 15 sec, 53°C for 15 sec, and 72°C for 1 min; 72°C for 5 min. Primer sequences are listed in Supplemental Table 2. Densitometric determination of signal intensity in each ChIP and input sample was calculated using NIH ImageJ. Fold enrichment was determined by calculating the ratio of PCR product intensities in ChIP Dex/Mock to Input Dex/Mock. In cases where amplicons were absent, an arbitrary value of 10 was assigned.

### Bacterial Infection Assays and ETI Elicitations

*Pseudomonas syringae* pv. *tomato* DC3000 (*Pst*) and *Pst avrRpm1* (*Psta*) were used for bacterial infection assays and ETI elicitations. A single colony of *Pst* was grown in 2 mL of LB medium containing 25 μg/mL rifampicin (Sigma-Aldrich). A single colony of *Psta* from a freshly streaked 3-day-old agar plate was grown in 50 mL of LB medium containing 25 μg/ml rifampicin and 50 μg/mL kanamycin (IBI Scientific, Peosta, IA).

Both strains were grown to log phase, washed in sterile water twice or once, respectively, resuspended in sterile water to OD_600_ of 0.2, and incubated at room temperature with no agitation for 6 and ∼2.5 hr, respectively, prior to infection. Bacterial infection assays on 4-to-5-week-old adult leaves were performed as described in Barco *et al*. (2019b).

## ACKNOWLEDGEMENTS

We thank J.L. Celenza for *cyp79B2 cyp79B3*, E.S. Sattely for ICN/ICN-ME, 4OH-ICA/4OH-ICA-ME and camalexin standards, and J.C. Miller for valuable feedback. This work was supported by T32-GM007499 (to B.B.) and Elsevier/Phytochemistry Young Investigator Award (to N.K.C.).

## AUTHOR CONTRIBUTIONS

B.B. and N.K.C performed pathogen assays, immunoblot assays, and ChIP experiments. B.B. performed all other experiments. B.B. and N.K.C. interpreted the results and wrote the paper.

## References

1. 1001 Genomes Consortium. (2016). 1,135 Genomes Reveal the Global Pattern of Polymorphism in *Arabidopsis thaliana*. Cell 166: 481–491.

2. Aharoni, A., and Galili, G. (2011). Metabolic engineering of the plant primary-secondary metabolism interface. Curr. Opin. Biotechnol. 22: 239–244.

3. Alon, U. (2007). Network motifs: theory and experimental approaches. Nat. Rev. Genet. 8: 450–461.

4. Aoyama, T., and Chua, N.H. (1997). A glucocorticoid-mediated transcriptional induction system in transgenic plants. Plant J. 11: 605–612.

5. Ascensao, J.A., Datta, P., Hancioglu, B., Sontag, E., Gennaro, M.L., and Igoshin, O.A. (2016). Non-monotonic response to monotonic stimulus: regulation of glyoxylate shunt gene-expression dynamics in *Mycobacterium tuberculosis*. PLoS Comput. Biol. 12: e1004741.

6. Austin, R.S., Hiu, S., Waese, J., Ierullo, M., Pasha, A., Wang, T.T., Fan, J., Foong, C., Breit, R., Desveaux, D., Moses, A., and Provart, N.J. (2016). New BAR tools for mining expression data and exploring *Cis*-elements in *Arabidopsis thaliana*. Plant J. 88: 490–504.

7. Aziz, A., Liu, Q.-C., and Dilworth, F.J. (2010). Regulating a master regulator: establishing tissue-specific gene expression in skeletal muscle. Epigenetics 5: 691–695.

8. Baghalian, K., Hajirezaei, M.-R., and Schreiber, F. (2014). Plant metabolic modeling: achieving new insight into metabolism and metabolic engineering. Plant Cell 26: 3847–3866.

9. Bak, S., and Feyereisen, R. (2001). The involvement of two p450 enzymes, CYP83B1 and CYP83A1, in auxin homeostasis and glucosinolate biosynthesis. Plant Physiol. 127: 108–118.

10. Bak, S., Tax, F.E., Feldmann, K.A., Galbraith, D.W., and Feyereisen, R. (2001). CYP83B1, a cytochrome P450 at the metabolic branch point in auxin and indole glucosinolate biosynthesis in Arabidopsis. Plant Cell 13: 101–111.

11. Barabási, A.-L., and Oltvai, Z.N. (2004). Network biology: understanding the cell’s functional organization. Nat. Rev. Genet. 5: 101–113.

12. Barco, B., and Clay, N.K. (2019a). Evolution of glucosinolate diversity via whole-genome duplications, gene rearrangements, and substrate promiscuity. Annu. Rev. Plant Biol. 8: 32.

13. Barco, B., Kim, Y., and Clay, N.K. (2019b). Expansion of a core regulon by transposable elements promotes Arabidopsis chemical diversity and pathogen defense. Nat Commun. In press. https://doi.org/10.1101/368340

14. Barco, B., Zipperer, L., and Clay, N.K. (2019c). Catalytic promiscuity potentiated the divergence of new cytochrome P450 enzyme functions in cyanogenic defense metabolism. bioRxiv 398503.

15. Barthole, G., To, A., Marchive, C., Brunaud, V., Soubigou-Taconnat, L., Berger, N., Dubreucq, B., Lepiniec, L., and Baud, S. (2014). MYB118 represses endosperm maturation in seeds of *Arabidopsis*. Plant Cell 26: 3519–3537.

16. Basu, S., Mehreja, R., Thiberge, S., Chen, M.T., and Weiss, R. (2004). Spatiotemporal control of gene expression with pulse-generating networks. Proc. Natl. Acad. Sci. USA 101: 6355–6360.

17. Basu, S., Gerchman, Y., Collins, C.H., Arnold, F.H., and Weiss, R. (2005). A synthetic multicellular system for programmed pattern formation. Nature 434: 1130–1134.

18. Bednarek, P., Piślewska-Bednarek, M., Svatoš, A., Schneider, B., Doubský, J., Mansurova, M., Humphry, M., Consonni, C., Panstruga, R., Sanchez-Vallet, A., and Molina, A. (2009). A glucosinolate metabolism pathway in living plant cells mediates broad-spectrum antifungal defense. Science 323: 101–106.

19. Bednarek, P., Piślewska-Bednarek, M., van Themaat, E.V.L., Maddula, R.K., Svatoš, A., and Schulze-Lefert, P. (2011). Conservation and clade-specific diversification of pathogen-inducible tryptophan and indole glucosinolate metabolism in *Arabidopsis thaliana* relatives. New Phytol. 192: 713–726.

20. Bekaert, M., Edger, P.P., Hudson, C.M., Pires, J.C., and Conant, G.C. (2012). Metabolic and evolutionary costs of herbivory defense: systems biology of glucosinolate synthesis. New Phytol. 196: 596–605.

21. Birkenbihl, R.P., Diezel, C., and Somssich, I.E. (2012). Arabidopsis WRKY33 is a key transcriptional regulator of hormonal and metabolic responses toward *Botrytis cinerea* infection. Plant Physiol. 159: 266–285.

22. Birkenbihl, R.P., Kracher, B., Rocarro M, and Somssich IE. (2017). Induced genome-wide binding of three Arabidopsis WRKY transcription factors during early MAMP-triggered immunity. Plant Cell 29: 20–38.

23. Bisgrove, S.R., Simonich, M.T., Smith, N.M., Sattler, A., and Innes, R.W. (1994). A disease resistance gene in Arabidopsis with specificity for two different pathogen avirulence genes. Plant Cell 6: 927–933.

24. Boerjan, W., Cervera, M.T., Delarue, M., Beeckman, T., Dewitte, W., Bellini, C., Caboche, M., Van Onckelen, H., Van Montagu, M., and Inzé, D.. (1995). Superroot, a recessive mutation in Arabidopsis, confers auxin overproduction. Plant Cell 7: 1405–1419.

25. Bohman, S., Staal, J., Thomma, B.P.H.J., Wang, M., and Dixelius, C. (2004). Characterisation of an *Arabidopsis-Leptosphaeria maculans* pathosystem: resistance partially requires camalexin biosynthesis and is independent of salicylic acid, ethylene and jasmonic acid signaling. Plant J. 37: 9–20.

26. Bonawitz, N.D., and Chapple, C. (2013). Can genetic engineering of lignin deposition be accomplished without an unacceptable yield penalty? Curr. Opin. Biotech. 24: 336–343.

27. Böttcher, C., Westphal, L., Schmotz, C., Prade, E., Scheel, D., and Glawischnig, E. (2009). The multifunctional enzyme CYP71B15 (PHYTOALEXIN DEFICIENT3) converts cysteine-indole-3-acetonitrile to camalexin in the indole-3-acetonitrile metabolic network of *Arabidopsis thaliana*. Plant Cell 21: 1830–1845.

28. Boyer, L.A., Lee, T.I., Cole, M.F., Johnstone, S.E., Levine, S.S., Zucker, J.P. Guenther, M.G., Kumar, R.M., Murray, H.L., Jenner, R.G., Gifford, D.K., Melton, D.A., Jaenisch, R., and Young, R.A. (2005). Core transcriptional regulatory circuitry in human embryonic stem cells. Cell 122: 947–956.

29. Brady, S.M., Zhang, L., Megraw, M., Martinez, N.J., Jiang, E., Yi, C.S., Liu, W., Zeng, A., Taylor-Teeples, M., Kim, D., Ahnert, S., Ohler, U., Ware, D., Walhout, A.J.M., and Benfey, P.N. (2011). A stele-enriched gene regulatory network in the Arabidopsis root. Mol. Syst. Biol 7: 459.

30. Celenza, J.L., Quiel, J.A., Smolen, G.A., Merrikh, H., Silvestro, A.R., Normanly, J., Bender, J. (2005). The Arabidopsis ATR1 Myb transcription factor controls indole glucosinolate homeostasis. Plant Physiol. 137: 253–262.

31. Chae, L., Kim, T., Nilo-Poyanco, R., and Rhee, S.Y. (2014). Genomic signatures of specialized metabolism in plants. Science 344: 510–513.

32. Clay, N.K., Adio, A.M., Denoux, C., Jander, G., and Ausubel, F.M. (2009). Glucosinolate metabolites required for an *Arabidopsis* innate immune response. Science 323: 95–101.

33. Clarke, D.B. (2010). Glucosinolates, structures and analysis in food. Anal. Methods 2: 301–416.

34. Chen, H., Wang, J.P., Liu, H., Li, H., Lin, Y.-C.J., Shi, R., Yang, C., Gao, J., Zhou, C., Li, Q, Sereroff, R.R., Li, W., and Chiang, V.L. (2019). Hierarchical transcription factor and chromatin binding network for wood formation in *Populus trichocarpa*. Plant Cell.

35. Cheng, C., Yan, K.K., Hwang, W., Qian, J., Bhardwaj, N., Rozowsky, J., Lu, Z.J., Niu, W., Alves, P., Kato, M, Snyder, M., and Gerstein, M. (2011). Construction and analysis of an integrated regulatory network derived from high-throughput sequencing data. PloS Comput. Biol. 7: e1002190.

36. Chepyala, S.R., Chen, Y.-C., Yan, C.-C.S., Lu, C.-Y.D., Wu, Y.-C., and Hsu, C.-P. (2016). Noise propagation with interlinked feed-forward pathways. Sci. Rep. 6: 23607.

37. Chezem, W.R., and Clay, N.K. (2016). Regulation of plant secondary metabolism and associated specialized cell development by MYBs and bHLHs. Phytochemistry 131: 26–43.

38. Chezem, W.R., Memon, A., Li, F.S., Weng, J.K., Clay, N.K. (2017). SG2-Type R2R3-MYB Transcription Factor MYB15 Controls Defense-Induced Lignification and Basal Immunity in Arabidopsis. Plant Cell 29: 1907–1926.

39. Clough, S.J., and Bent, A.F. (1998). Floral dip: a simplified method for *Agrobacterium*-mediated transformation of *Arabidopsis thaliana*. Plant J. 16: 735–743.

40. Colón, A.M., Sengupta, N., Rhodes, D. Dudareva, N., and Morgan, J. (2010). A kinetic model describes metabolic responses to perturbations and distribution of flux control in the benzenoid network of *Petunia hybrida*. Plant J. 62: 64–76.

41. Consonni, C., Bednarek, P., Humphry, M., Francocci, F., Ferrari, S., Harzen, A., van Themaat, E.V., and Panstruga, R. (2010). Tryptophan-derived metabolites are required for antifungal defense in the Arabidopsis *mlo2* mutant. Plant Physiol, 152: 1544–1561.

42. Cournac, A., and Sepulchre, J.A. (2009). Simple molecular networks that respond optimally to time-periodic stimulation. BMC Syst. Biol. 3: 29.

43. Daran-Lapujade, P. Jansen, M.L.A., Daran, J.-M., van Gulik, W., de Winde, J.H., and Pronk, J.T. (2004). Role of transcriptional regulation in controlling fluxes in central carbon metabolism of *Saccharomyces cerevisiae.* A chemostat culture study. J. Biol. Chem. 279: 9125–9138.

44. de Ronde, W.H., Tostevin, F., and ten Wolde, P.R. (2012). Feed-forward loops and diamond motifs lead to tunable transmission of information in the frequency domain. Phys. Rev. E. 86: 021913.

45. Defoort, J., Van de Peer, Y., and Vermeirssen, V. (2018). Function, dynamics and evolution of network motif modules in integrated gene regulatory networks of worm and plant. Nucleic Acids Res. 46: 6480–6503.

46. Dellaporta, S.L., Wood, J., and Hicks, J.B. (1983). A plant DNA minipreparation: version II. Plant Mol. Biol. Rep. 1: 19–21.

47. Denoux, C., Galletti, R., Mammarella, N., Gopalan, S., Werck, D., De Lorenzo, G., Ferrari, S., Ausubel, F.M., and Dewdney, J. (2008). Activation of defense response pathways by OGs and Flg22 elicitors in Arabidopsis seedlings. Mol. Plant 1: 423–445.

48. Dixon, R.A., and Strack, D. (2003). Phytochemistry meets genome analysis and beyond. Phytochemistry 62: 815–816.

49. Dobrin, R., Berg, Q.K., Barabási, A.-L., and Oltvai, Z.N. (2004). Aggregation of topological motifs in the *Escherichia coli* transcriptional regulatory network. BMC Bioinformatics 5: 10.

50. Dubos, C., Stracke, R., Grotewold, E., Weisshaar, B., Martin, C., and Lepiniec, L. (2010). MYB transcription factors in *Arabidopsis*. Trends Plant Sci. 15: 573–581.

51. Entus, R., Aufderheide, B., and Sauro, H.M. (2007). Design and implementation of three incoherent feed-forward motif based biological concentration sensors. Syst. Synth. Biol. 1: 119–128.

52. Even, S., Lindley, N.D., and Cocaign-Bousquet, M. (2003). Transcriptional, translational and metabolic regulation of glycolysis in *Lactococcus lactis* subsp. *cremoris* MG1363 grown in continuous acidic cultures. Microbiology 149: 1935–1944.

53. Fahey, J.W., Zalemann, A.T., and Talalay, P. (2001). The chemical diversity and distribution of glucosinolates and isothiocyanates among plants. Phytochemistry 56: 5–51.

54. Fan, M., Bai, M.-Y., Kim, J.-G., Wang, T., Oh, E., Chen, L., Park, C.H., Son, S.-H., Kim, S.-K., Mudgett, M.B., and Wang, Z.-Y. (2014). The bHLH transcription factor HBI1 mediates the trade-off between growth and pathogen-associated molecular pattern-triggered immunity in *Arabidopsis*. Plant Cell 26: 828–841.

55. Fell, D. (1997). Understanding the Control of Metabolism. (London: Portland Press).

56. Ferrari, S., Galletti, R., Denoux, C., De Lorenzo, G., Ausubel, F.M., and Dewdney, J. (2003). Resistance to *Botrytis cinerea* induced in Arabidopsis by elicitors I independent of salicylic acid, ethylene, or jasmonate signaling but requires *PHYTOALEXIN DEFICIENT3*. Plant Physiol. 144: 367–379.

57. Frerigmann, H., and Gigolashvili, T. (2014a). MYB34, MYB51, and MYB122 distinctly regulate indolic glucosinolate biosynthesis in *Arabidopsis thaliana*. Mol. Plant 7: 814–828.

58. Frerigmann, H., Berger, B., and Gigolashvili, T. (2014b). bHLH05 is an interaction partner of MYB51 and a novel regulator of glucosinolate biosynthesis in *Arabidopsis*. Plant Physiol. 166: 349–369.

59. Frerigmann, H., Glawischnig, E., and Gigolashvili, T. (2015). The role of MYB34, MYB51 and MYB122 in the regulation of camalexin biosynthesis in *Arabidopsis thaliana*. Front. Plant Sci. 6: 654.

60. Frerigmann, H., Piślewska-Bednarek, M., Sánchez-Vallet, A., Molina, A., Glawischnig, E., Gigolashvili, T., and Bednarek, P. (2016). Regulation of pathogen-triggered tryptophan metabolism in *Arabidopsis thaliana* by MYB transcription factors and indole glucosinolate conversion products. Mol. Plant 9: 682–695.

61. Gachon, C.M., Langlois-Meurinne, M., Henry, Y., and Saindrenan, P. (2005). Transcriptional co-regulation of secondary metabolism enzymes in *Arabidopsis*: functional and evolutionary implications. Plant Mol. Biol. 58: 229–245.

62. Gao, Z., Chen, S., Qin, S., and Tang, C. (2018). Network motifs capable of decoding transcription factor dynamics. Sci. Rep. 8: 3594.

63. Gerstein, M.B., Lu, Z.J., Van Nostrand, E.L., Cheng, C., Arshinoff, B.I., Liu, T., Yip, K.Y., Robilotto, R., Rechtsteiner, A., Ikegami, K., Alves, P., Chateigner, A., Perry, M., Morris, M., Auerbach, R.K., Feng, X., Leng, J., Vielle, A., Niu, W., Rhrissorrakrai, K., Agarwal, A., Alexander, R.P., Barber, G., Brdlik, C.M., Brennan, J., Brouillet, J.J., Carr, A., Cheung, M.S., Clawson, H., Contrino, S., Dannenberg, L.O., Dernburg, A.F., Desai, A., Dick, L., Dose, A.C., Du, J., Egelhofer, T., Ercan, S., Euskirchen, G., Ewing, B., Feingold, E.A., Gassmann, R., Good, P.J., Green, P., Gullier, F., Gutwein, M., Guyer, M.S., Habegger, L., Han, T., Henikoff, J.G., Henz, S.R., Hinrichs, A., Holster, H., Hyman, T., Iniguez, A.L., Janette, J., Jensen, M., Kato, M., Kent, W.J., Kephart, E., Khivansara, V., Khurana, E., Kim, J.K., Kolasinska Zwierz, P., Lai, E.C., Latorre, I., Leahey, A., Lewis, S., Lloyd, P., Lochovsky, L., Lowdon, R.F., Lubling, Y., Lyne, R., MacCoss, M., Mackowiak, S.D., Mangone, M., McKay, S., Mecenas, D., Merrihew, G., Miller, D.M., 3rd, Muroyama, A., Murray, J.I., Ooi, S.L., Pham, H., Phippen, T., Preston, E.A., Rajewsky, N., Ratsch, G., Rosenbaum, H., Rozowsky, J., Rutherford, K., Ruzanov, P., Sarov, M., Sasidharan, R., Sboner, A., Scheid, P., Segal, E., Shin, H., Shou, C., Slack, F.J., Slightam, C., Smith, R., Spencer, W.C., Stinson, E.O., Taing, S., Takasaki, T., Vafeados, D., Voronina, K., Wang, G., Washington, N.L., Whittle, C.M., Wu, B., Yan, K.K., Zeller, G., Zha, Z., Zhong, M., Zhou, X., mod, E.C., Ahringer, J., Strome, S., Gunsalus, K.C., Micklem, G., Liu, X.S., Reinke, V., Kim, S.K., Hillier, L.W., Henikoff, S., Piano, F., Snyder, M., Stein, L., Lieb, J.D., and Waterston, R.H. (2010). Integrative analysis of the *Caenorhabditis elegans* genome by the modENCODE project. Science 330: 1775–1787.

64. Gigolashvili, T., Berger, B., Mock, H.P., Müller, C., Weisshaar, B., and Flügge, U.I. (2007a). The transcription factor HIG1/MYB51 regulates indolic glucosinolate biosynthesis in *Arabidopsis thaliana*. Plant J. 50: 886–901.

65. Gigolashvili, T., Yatusevich, R., Berger, B., Møller, C., and Flügge, U.I. (2007b). The R2R3 MYB transcription factor HAG1/MYB28 is a regulator of methionine derived glucosinolate biosynthesis in *Arabidopsis thaliana*. Plant J. 51: 247–261.

66. Gigolashvili, T., Engqvist, M., Yatusevich, R., Møller, C., and Flügge, U.I. (2008). HAG2/ MYB76 and HAG3/MYB29 exert a specific and coordinated control on the regulation of aliphatic glucosinolate biosynthesis in *Arabidopsis thaliana*. New Phytol. 177: 627–642.

67. Glawischnig, E., Hansen, B.G., Olsen, C.E., and Halkier, B.A. (2004). Camalexin is synthesized from indole-3-acetaldoxime, a key branching point between primary and secondary metabolism in *Arabidopsis*. Proc. Natl. Acad. Sci. USA 101: 8245–8250.

68. Goentoro, L., Shoval, O., Kirschner, M.W., and Alon, U. (2009). The incoherent feedforward loop can provide fold-change detection in gene regulation. Mol. Cell 36: 894–899.

69. González-Lamothe, R., Boyle, P., Dulude, A., Roy, V., Lezin-Doumbou, C., Kaur, G.S., Bouarab, K., Després, C., and Brisson, N. (2008). The transcriptional activator Pti4 is required for the recruitment of a repressosome nucleated by repressor SEBF at the potato *PR-10a* gene. Plant Cell 20: 3136–3147.

70. Grotewold, E. (2005). Plant metabolic diversity: a regulatory perspective. Trends Plant Sci. 10: 57–62.

71. Grubb, C.D., Masuno, M.N., Molinski, T.F., and Abel, S. (2004). Arabidopsis glucosyltransferase UGT74B1 functions in glucosinolate biosynthesis and auxin homeostasis. Plant J. 40: 893–908.

72. Gutenkunst, R.N., Waterfall, J.J., Casey, F.P., Brown, K.S., Myers, C.R., and Sethna, J.P. (2007). Universally sloppy parameter sensitivities in system biology models. PLoS Comput. Biol. 3: 1871–1878.

73. Hartmann, T. (2007). From waste products to ecochemicals: fifty years research of plant secondary metabolism. Phytochemistry 68: 2831–2846.

74. Heyndrickx, K.S., Van de Velde, J., Wang, C., Weigel, D., and Vandepoele, K. (2014). A functional and evolutionary perspective on transcription factor binding in *Arabidopsis thaliana*. Plant Cell 26: 3894–3910.

75. Hiratsu, K., Matsui, K., Koyama, T., and Ohme-Takagi, M. (2003). Dominant repression of target genes by chimeric repressors that include the EAR motif, a repression domain, in *Arabidopsis*. Plant J. 34: 733–739.

76. Hiruma, K., Onozawa-Komori, M., Takahashi, F., Asakura, M., Bednarek, P., Okuno, T., Schulze-Lefert, P., and Takano, Y. (2010). Entry mode-dependent function of an indole glucosinolate pathway in Arabidopsis for nonhost resistance against anthracnose pathogens. Plant Cell 22: 2429–2443.

77. Hogge, L.R., Reed, D.W., Underhill, E.W., and Haughn, G.W. (1988). HPLC separation of glucosinolates from leaves and seeds of *Arabidopsis thaliana* and their identification using thermospray liquid chromatography-mass spectrometry. J. Chromatog. Sci. 26: 551–556.

78. Humphry, M., Bednarek, P., Kemmerling, B., Koh, S., Stein, M., Göbel, U., Stüber, K., Pislewska-Bednarek, M., Loraine, A., Schulze-Lefert, P., Somerville, S., and Panstruga, R. (2010). A regulon conserved in monocot and dicot plants defines a functional module in antifungal plant immunity. Proc. Natl. Acad. Sci. USA 107: 21896–21901.

79. Ikeda, M., Mitsuda, N., and Ohme-Takagi, M. (2009). Arabidopsis WUSCHEL is a bifunctional transcription factor that acts as a repressor in stem cell regulation and as an activator in floral patterning. Plant Cell 21: 3493–3505.

80. Jaeger, K.E., Pullen, N., Lamzin, S., Morris, R.J., and Wigge, P.A. (2013). Interlocking feedback loops govern the dynamic behavior of the floral transition in *Arabidopsis*. Plant Cell 25: 820–833.

81. James, A.M., Ma, D., Mellway, R., Gesell, A., Yoshida, K., Walker, V., Tran, L., Stewart, D., Reichelt, M., Suvanto, J., Salminen, J.P, Gershenzon, J., Séguin, A., and Constabel, C.P. (2017). Poplar MYB115 and MYB134 transcription factors regulate proanthocyanidin synthesis and structure. Plant Physiol. 174: 154–171.

82. Jin, H., Cominelli, E., Bailey, P., Parr, A., Mehrtens, F., Jones, J., Tonelli, C., Weisshaar, B., and Martin, C. (2000). Transcriptional repression by AtMYB4 controls production of UV-protecting sunscreens in *Arabidopsis*. EMBO J. 19: 6150–6161.

83. Jin, J., He, K., Tang, X., Li, Z., Lv, L., Zhao, Y., Luo, J., and Gao, G. (2015). An *Arabidopsis* transcriptional regulatory map reveals distinct functional and evolutionary features of novel transcription factors. Mol. Biol. Evol. 32: 1767–1773.

84. Joanito, I., Chu, J.-W., Wu, S.-H., and Hsu, C.-P. (2018). An incoherent feed-forward loop switches the Arabidopsis clock rapidly between two hysteretic states. Sci. Rep. 8: 13944.

85. Jones, J.D.G., and Dangl, J.L. (2006). The plant immune system. Nature 444: 323–329.

86. Kalir, S., Mangan, S., and Alon, U. (2005). A coherent feed-forward loop with a SUM input function prolongs flagella expression in *Escherichia coli*. Mol. Syst. Biol. 1: 2005.0006.

87. Kaplan, S., Bren, A., Dekel, E., and Alon, U. (2008). The incoherent feed-forward loop can generate non-monotonic input functions for genes. Mol. Syst. Biol. 4: 203.

88. Klein, A.P., Anarat-Cappillino, G., and Sattely, E.S. (2013). Minimum set of cytochromes P450 for reconstituting the biosynthesis of camalexin, a major *Arabidopsis* antibiotic. Angew. Chem. Int. Ed. Engl. 52: 13625–13628.

89. Kliebenstein, D.J., Kroymann, J., Brown, P., Figuth, A., Pedersen, D., Gershenzon, J., and Mitchell-Olds, T. (2001). Genetic control of natural variation in Arabidopsis glucosinolate accumulation. Plant Physiol. 126: 811–825.

90. Kliebenstein, D.J., and Osbourn, A. (2012). Making new molecules - evolution of pathways for novel metabolites in plants. Curr. Opin. Plant Biol. 15: 415–423.

91. Lahrmann, U., Strehmel, N., Langen, G., Frerigmann, H., Leson, L., Ding, Y., Scheel, D., Herklotz, S., Hilbert, M., and Zuccaro, A. (2015). Mutualistic root endophytism is not associated with the reduction of saprotrophic traits and requires a noncompromised plant innate immunity. New Phytol. 207: 841–857.

92. Lavenus, J., Goh, T., Guyomarc’h, S., Hill, K., Lucas, M., Voβ, U., Kenobi, K., Wilson, M.H., Farcot, E., Hagen, G., Guilfoyle, T.J., Fukaki, H., Laplaze, L., and Bennett, M.J. (2015). Inference of the Arabidopsis lateral root gene regulatory network suggests a bifurcation mechanism that defines primordial flanking and central zones. Plant Cell 27: 1368–1388.

93. Lee, R.E.C., Walker, S.R., Savery, K., Frank, D.A., Gaudet, S. (2014). Nuclear NF-kB determines TNF-induced transcription in single cells. Mol. Cell 53: 867–879.

94. Lee, T.I., Rinaldi, N.J., Robert, F., Odom, D.T., Bar-Joseph, Z., Gerber, G.K., Hannett, N.M., Harbison, C.T., Thompson, C.M., Simon, I., Zeitlinger, J., Jennings, E.G., Murray, H.L., Gordon, D.B., Ren, B., Wyrick, J.J., Tagne, J.B., Volkert, T.L., Fraenkel, E., Gifford, D.K., and Young, R.A. (2002). Transcriptional regulatory networks in *Saccharomyces cerevisiae*. Science 298: 799–804.

95. Lewis, L.A., Polanski, K., de Torres-Zabala, M., Jayaraman, S., Bowden, L., Moore, J., Penfold, C.A., Jenkins, D.J., Hill, C., Baxter, L., Kulasekaran, S., Truman, W., Littlejohn, G., Prusinska, J., Mead, A., Steinbrenner, J., Hickman, R., Rand, D., Wild, D.L., Ott, S., Buchanan-Wollaston, V., Smirnoff, N., Beynon, J., Denby, K., and Grant, M. (2015). Transcriptional dynamics driving MAMP-triggered immunity and pathogen effector-mediated immunosuppression in Arabidopsis leaves following infection with *Pseudomonas syringae* pv tomato DC3000. Plant Cell 27: 3038–3064.

96. Li, B., Gaudinier, A., Tang, M., Taylor-Teeples, M., Nham, N.T., Ghaffari, C., Benson, D.S., Steinmann, M., Gray, J.A., Brady, S.M., and Kliebenstein, D.J. (2014). Promoter-based integration in plant defense regulation. Plant Physiol. 166: 1803–1820.

97. Li, G., Meng, X., Wang, R., Mao, G., Han, L., Liu, Y., and Zhang, S. (2012). Dual-level regulation of ACC synthase activity by MPK3/MPK6 cascade and its downstream WRKY transcription factor during ethylene induction in Arabidopsis. PLoS Genet. 8: e1002767.

98. Lin, Y.C., Li, W., Sun, Y.H., Kumari, S., Wei, H., Li, Q., Tunlaya-Anukit, S., Sederoff, R.R., and Chiang, V.L. (2013). SND1 transcription factor-directed quantitative functional hierarchical genetic regulatory network in wood formation in *Populus trichocarpa*. Plant Cell 25: 4324–4341.

99. Lipka, V., Dittgen, J., Bednarek, P., Bhat, R., Wiermer, M., Stein, M., Landtag, J., Brandt, W., Rosahl, S., Scheel, D., and Llorente, F. (2005). Pre-and postinvasion defenses both contribute to nonhost resistance in *Arabidopsis*. Science 310: 1180–1183.

100. Liu, S., Kracher, B., Ziegler, J., Birkenbihl, R.P., and Somssich, I.E. (2015). Negative regulation of ABA signaling by WRKY33 is critical for *Arabidopsis* immunity towards *Botrytis cinerea* 2100. eLife 4: e07295.

101. Logemann, E., Birkenbihl, R.P., Rawat, V., Schneeberger, K., Schmelzer, E., and Somssich, I.E. (2013). Functional dissection of the PROPEP2 and PROPEP3 promoters reveals the importance of WRKY factors in mediating micobe-associated molecular pattern-induced expression. New Phytol. 198: 1165–1177.

102. Lozano-Durán, R., Macho, A.P., Boutrot, F., Segonzac, C., Somssich, I.E., and Zipfel, D (2013). The transcriptional regulator BZR1 mediates trade-off between plant innate immunity and growth. Elife 2: e00983.

103. Ma, H.-W., Buer, J., and Zeng, A.-P. (2004). Hierarchical structure and modules in the Escherichia coli transcriptional regulatory network revealed by a new top-down approach. BMC Bioinformatics 5: 199.

104. Ma, W., Trusina, A., El-Samad, H., Lim, W.A., and Tang, C. (2009). Defining network topologies that can achieve biochemical adaptation. Cell 138: 760–773.

105. Ma’ayan, A., Jenkins, S.L., Neves, S., Hasseldine, A., Grace, E., Dubin-Thaler, B., Eungdamrong, N.J., Weng, G., Ram, P.T., Rice, J.J., Kershenbaum, A., Stolovitzky, G.A., Blitzer, R.D., and Iyengar, R. (2005). Formation of regulatory patterns during signal propagation in a mammalian cellular network. Science 309: 1078–1083.

106. Malinovsky, F.G., Batoux, M., Schwessinger, B., Youn, J.H., Stransfeld, L., Win, J., Kim, S.K., and Zipfel, C. (2014). Antagonistic regulation of growth and immunity by the Arabidopsis basic helix-loop-helix transcription factor HOMOLOG OF BRASSINOSTEROID ENHANCED EXPRESSION2 INTERACTING WITH INCREASED LEAF INCLINATION1 BINDING BHLH1. Plant Physiol. 164: 1443–1455.

107. Malitsky, S., Blum, E., Less, H., Venger, I., Elbaz, M., Morin, S., Eshed, Y., and Aharoni, A. (2008). The transcript and metabolite networks affected by the two clades of Arabidopsis glucosinolate biosynthesis regulators. Plant Physiol. 148: 2021–2049.

108. Mangan, S., and Alon, U. (2003a). Structure and function of the feed-forward loop network motif. Proc. Natl. Acad. Sci. USA 100: 11980–11985.

109. Mangan, S., Zaslaver, A., and Alon, U. (2003b). The coherent feedforward loop serves as a sign-sensitive delay element in transcriptional networks. J. Mol. Biol. 334: 197–204.

110. Mangan, S., Itzkovitz, S., Zaslaver, A., and Alon, U. (2006). The incoherent feed-forward loop accelerates the response-time of the gal system of *Escherichia coli*. J. Mol. Biol. 356: 1073–1081.

111. Martin, C., Ellis, N., and Rook, F. (2010). Do transcription factors play special roles in adaptive variation? Plant Physiol. 154: 506–511.

112. Mendoza-Parra, M.A., Pattabhiraman, S., and Gronemeyer, H. (2012). Sequential chromatin immunoprecipitation protocol for global analysis through massive parallel sequencing (reChIP-seq). Protocol Exchange 10.

113. Miao, Y., Laun, T., Zimmermann, P., and Zentgraf, U. (2004). Targets of the WRKY53 transcription factor and its role during leaf senescence in *Arabidopsis*. Plant Mol. Biol. 55: 853–867.

114. Mikkelsen, M.D., Hansen, C.H., Wittstock, U., and Halkier, B.A. (2000). Cytochrome P450 CYP79B2 from *Arabidopsis* catalyzes the conversion of tryptophan to indole-3-acetaldoxime, a precursor of indole glucosinolates and indole-3-acetic acid. J. Biol. Chem. 275: 33712–33717.

115. Mikkelsen, M.D., Naur, P., and Halkier, B.A. (2004). *Arabidopsis* mutants in the C-S lyase of glucosinolate biosynthesis establish a critical role for indole-3-acetaldoxime in auxin homeostasis. Plant J. 37: 770–777.

116. Milo, R., Shen-Orr, S., Kashtan, N., Chklovskii, D., and Alon, U. (2002). Network motifs: simple building blocks of complex networks. Science 298: 824–827.

117. modEncode Consortium, Roy, S., Ernst, J., Kharchenko, P.V., Kheradpour, P., Negre, N., Eaton, M.L., Landolin, J.M., Bristow, C.A., Ma, L., Lin, M.F., Washietl, S., Arshinoff, B.I., Ay, F., Meyer, P.E., Robine, N., Washington, N.L., Di Stefano, L., Berezikov, E., Brown, C.D., Candeias, R., Carlson, J.W., Carr, A., Jungreis, I., Marbach, D., Sealfon, R., Tolstorukov, M.Y., Will, S., Alekseyenko, A.A., Artieri, C., Booth, B.W., Brooks, A.N., Dai, Q., Davis, C.A., Duff, M.O., Feng, X., Gorchakov, A.A., Gu, T., Henikoff, J.G., Kapranov, P., Li, R., MacAlpine, H.K., Malone, J., Minoda, A., Nordman, J., Okamura, K., Perry, M., Powell, S.K., Riddle, N.C., Sakai, A., Samsonova, A., Sandler, J.E., Schwartz, Y.B., Sher, N., Spokony, R., Sturgill, D., van Baren, M., Wan, K.H., Yang, L., Yu, C., Feingold, E., Good, P., Guyer, M., Lowdon, R., Ahmad, K., Andrews, J., Berger, B., Brenner, S.E., Brent, M.R., Cherbas, L., Elgin, S.C., Gingeras, T.R., Grossman, R., Hoskins, R.A., Kaufman, T.C., Kent, W., Kuroda, M.I., Orr-Weaver, T., Perrimon, N., Pirrotta, V., Posakony, J.W., Ren, B., Russell, S., Cherbas, P., Graveley, B.R., Lewis, S., Micklem, G., Oliver, B., Park, P.J., Celniker, S.E., Henikoff, S., Karpen, G.H., Lai, E.C., MacAlpine, D.M., Stein, L.D., White, K.P., and Kellis, M. (2010). Identification of functional elements and regulatory circuits by *Drosophila* modENCODE. Science 330: 1787–1797.

118. Moreno-Risueno, M.A., Busch, W., and Benfey, P.N. (2010). OmicsMurashige, T., and Skoog, F. (1962). A revised medium for rapid growth and bio assays with tobacco tissue cultures. Physiol. Plant. 15: 473–497.

119. Nafisi, M, Goregaoker, S., Botanga, C.J., Glawischnig, E., Olsen, C.E., Halkier, B.A., and Glazebrook, J. (2007). *Arabidopsis* cytochrome P450 monoxygenase 71A13 catalyzes the conversion of indole-3-acetaldoxime in camalexin synthesis. Plant Cell 19: 2039–2052.

120. Navarro, L., Zipfel, C., Rowland, O., Keller, I., Robatzek, S., Boller, T., and Jones, J.D.G. (2004). The transcriptional innate immune response to flg22. Interplay and overlap with Avr gene-deendent defense responses and bacterial pathogenesis. Plant Physiol. 135: 1113–1128.

121. Nelson, D., and Werck-Reichhart, D. (2011). A P450-centric view of plant evolution. Plant J. 66: 194–211.

122. Niu, W., Lu, Z.J., Zhong, M., Sarov, M., Murray, J.I., Brdlik, C.M., Janette, J., Chen, C., Alves, P., Preston, E., Slightham, C., Jiang, L., Hyman, A.A., Kim, S.K., Waterston, R.H., Gerstein, M., Snyder, M., and Reinke, V. (2011). Diverse transcription factor binding features revealed by genome-wide ChIP-seq in *C. elegans*. Genome Res. 21: 245–254.

123. Odom, D.T., Zizlsperger, N., Gordon, D.B., Bell, G.W., Rinaldi, N.J., Murray, H.L., Volkert, T.L., Schreiber, J., Rolfe, P.A., Gifford, D.K., Fraenkel, E., Bell, G.I., and Young, R.A. (2004). Control of pancreas and liver gene expression by HNF transcription factors. Science 303: 1378–1381.

124. Ohta, M., Matsui, K., Hiratsu, K., Shinshi, H., and Ohme-Takagi, M. (2001). Repression domains of class II ERF transcriptional repressors share an essential motif for active repression. Plant Cell 13: 1959–1968.

125. Olson-Manning, C.F., Lee, C.-R., Rausher, M.D., and Mitchell-Olds, T. (2013). Evolution of flux control in the glucosinolate pathway. Mol. Biol. Evol. 30: 14–23.

126. Omranian, N., Kleessen, S., Tohge, T., Klie, S., Basler, G., Mueller-Roeber, B., Fernie, A.R., Nikoloski, Z. (2015). Differential metabolic and coexpression networks of plant metabolism. Trends Plant Sci. 20: 266–268.

127. Pandey, S.P., Roccaro, M., Schön, M., Logemann, E., and Somssich, I.E. (2010). Transcriptional reprogramming regulated by WRKY18 and WRKY40 facilitates powdery mildew infection of Arabidopsis. Plant J. 64: 912–923.

128. Parcy, F., Nilsson, O., Busch, M.A., Lee, I., and Weigel, D. (1998). A genetic framework for floral patterning. Nature 395: 561–566.

129. Penn, B.H., Bergstrom, D.A., Dilworth, F.J., Bengal, E., and Tapscott, S.J. (2004). A MyoD-generated feed-forward circuit temporally patterns gene expression during skeletal muscle differentiation. Genes Dev. 18: 2348–2353.

130. Pfaffl, M.W. (2001). A new mathematical model for relative quantification in real-time RT-PCR. Nucleic Acids Res. 29: e45.

131. Pfalz, M., Vogel, H., and Kroymann, J. (2009). The gene controlling the *Indole Glucosinolate Modifier1* quantitative trait locus alters indole glucosinolate structures and aphid resistance in *Arabidopsis*. Plant Cell 21: 985–999.

132. Preston, J., Wheeler, J., Heazlewood, J., Li, S.F., and Parish, R.W. (2004). AtMYB32 is required for normal pollen development in *Arabidopsis thaliana*. Plant J. 40: 979–995.

133. Pullen, N., Jaeger, K.E., Wigge, P.A., and Morris, R.J. (2013). Simple network motifs can capture key characteristics of the floral transition in *Arabidopsis*. Plant Signal. Behav. 8: 11.

134. Qiu, J.L., Fiil, B.K., Petersen, K., Nielsen, H.B., Botanga, C.J., Thorgrimsen, S., Palma, K., Suarez-Rodriguez, M.C., Sandbech-Clausen, S., Lichota, J., and Brodersen, P. (2008). *Arabidopsis* MAP kinase 4 regulates gene expression through transcription factor release in the nucleus. EMBO J. 27: 2214–2221.

135. Raes, J., Rohde, A., Christensen, J.H., Van de Peer, Y., and Boerjan, W. (2003). Genome-wide characterization of the lignification toolbox in Arabidopsis. Plant Physiol. 133: 1051–1071.

136. Rajniak, J., Barco, B., Clay, N.K., and Sattely, E.S. (2015). A new cyanogenic metabolite in Arabidopsis required for inducible pathogen defence. Nature 525: 376–379.

137. Rinerson, C.I., Rabara, R.C., Tripathi, P., Shen, Q.J., and Rushton, P.J. (2015). The evolution of WRKY transcription factors. BMC Plant Biol. 15: 66.

138. Robatzek, S., and Somssich, I.E. (2002). Targets of *At*WRKY6 regulation during lant senescence and pathogen defense. Genes Dev. 16: 1139–1149.

139. Romero, I., Fuertes, A., Benito, M.J., Malpica, J.M., Leyva, A., and Paz-Ares, J. (1998). More than 80 *R2R3-MYB* regulatory genes in the genome of Arabidopsis thaliana. Plant J. 14: 273–284.

140. Rossell, S., van der Weijden, C.C., Kruckeberg, A.L., Bakker, B.M., and Westerhoff, H.V. (2005). Hierarchical and metabolic regulation of glucose influx in starved *Saccharomyces cerevisiae*. FEMS Yeast Res. 5: 611–619.

141. Rossell, S., van der Weijden, C.C., Lindenbergh, A., van Tuijl, A., Francke, C., Bakker, B.M., and Westerhoff, H.V. (2006). Unraveling the complexity of flux regulation: a new method demonstrated for nutrient starvation in *Saccharomyces cerevisiae*. Proc. Natl. Acad. Sci. USA 103: 2166–2171.

142. Rushton, P.J., Somssich, I.E., Ringler, P., and Shen, Q.J. (2010). WRKY transcription factors. Trends Plant Sci. 15: 247–258.

143. Saddic, L.A., Huvermann, B., Bezhani, S., Su, Y., Winter, C.M., Kwon, C.S., Collum, R.P., and Wagner, D. (2006). The LEAFY target LMI1 is a meristem identity regulator and acts together with LEAFY to regulate expression of *CAULIFLOWER*. Development 133: 1673–1682.

144. Sakuraba, Y., Kim, Y.-S., Han, S.-H., Lee, B.-D., and Paek, N.-C. (2015). The Arabidopsis transcription factor NAC016 promotes drought stress responses by repressing AREB1 transcription through a trifurcate feed-forward regulatory loop involving NAP. Plant Cell 27: 1771–1787.

145. Sanchez-Vallet, A., Ramos, B., Bednarek, P., López, G., Piślewska-Bednarek, M., Schulze-Lefert, P., and Molina, A. (2010). Tryptophan-derived secondary metabolites in *Arabidopsis thaliana* confer non-host resistance to necrotrophic *Plectosphaerella cucumerina* fungi. Plant J. 63: 115–127.

146. Schluttenhofer, C., and Yuan, L. (2015). Regulation of specialized metabolism by WRKY transcription factors. Plant Physiol. 167: 295–306.

147. Schuhegger, R., Nafisi, M., Mansourova, M., Petersen, B.L., Olsen, C.E., Svatoš, A., Halkier, B.A., and Glawischnig, E. (2006). CYP71B15 (PAD3) catalyzes the final step in camalexin biosynthesis. Plant Physiol. 141: 1248–1254.

148. Schlaeppi, K., Abou-Mansour, E., Buchala, A., and Mauch, F. (2010). Disease resistance of Arabidopsis to *Phytophthora brassicae* is established by the sequential action of indole glucosinolates and camalexin. Plant J. 62: 840–851.

149. Schranz, M.E., Edger, P.P., Pires, J.C., van Dam, N.M., and Wheat, C.W. (2011). Comparative genomics in the Brassicales: ancient genome duplications, glucosinolate diversification and Pierinae herbivore radiation. In: Genetics, genomics and breeding of oilseed brassicas (Edwards JBD, Parkin I, Kole C., eds) Jersey, UK: Science Publishers, pp. 206–218.

150. Schweizer, F., Fernández-Calvo, P., Zander, M., Diez-Diaz, M., Fonseca S., Glauser, G., Lewsey, M.G., Ecker, J.R., Solano, R., and Reymond, P. (2013). *Arabidopsis* basic helix-loop-helix transcription factors MYC2, MYC3, and MYC4 regulate glucosinolate biosynthesis, insect performance, and feeding behavior. Plant Cell 25: 3117–3132.

151. Semsey, S., Krishna, S., Sneppan K., and Adhya, S. (2007). Signal integration in the galactose network of *Escherichia coli*. Mol. Microbiol. 65: 465.

152. Sen, S., Kim, J., and Murray, R.M. (2014). Designing robustness to temperature in a feedforward loop circuit, in 53^rd^ IEEE Conference on Decision and Control, pp. 4629-4634.

153. Shen-Orr, S.S., Milo, R., Mangan, S., and Alon, U. (2002). Network motifs in the transcriptional regulation network of *Escherichia coli*. Nature Genet. 31: 64–68.

154. Sontag, E.D. (2009). Remarks on feedforward circuits, adaptation, and pulse memory. IET Systems Biol. 4: 39–51.

155. Stahl, E., Bellwon, P., Huber, S., Schlaeppi, K., Bernsdorff, F., Vallat-Michel, A., Mauch, F., and Zeier, J. (2016). Regulatory and functional aspects of indolic metabolism in plant systemic acquired resistance. Mol. Plant 9: 662–681.

156. Sugawara, S., Hishiyama, S., Jikumaru, Y., Hanada, A., Nishimura, T., Koshiba, T., Zhao, Y., Kamiya, Y., and Kasahara, H. (2009). Biochemical analyses of indole-3-acetaldoxime-dependent auxin biosynthesis in Arabidopsis. Proc. Natl. Acad. Sci. USA 106: 5430–5435.

157. Takeda, K., Shao, D., Adler, M., Charest, P.G., Loomis, W.F., Levine, H., Groisman, A., Rappel, W.J., and Firtel, R.A. (2012). Incoherent feedforward control governs adaptation of activated ras in a eukaryotic chemotaxis pathway. Sci. Signal. 5: ra2.

158. Tang, B., Hsu, H.-K., Hsu, P.-Y., Bonneville, R., Chen, S.-S., Huang, T.H.-M., and Jin, V.X. (2012). Hierarchical modularity in ERalpha transcriptional network is associated with distinct functions and implicates clinical outcomes. Sci. Rep. 2: 875.

159. Tao, Y., Xie, Z., Chen, W., Glazebrook, J., Chang, H.-S., Han, B., Zhu, T., Zou, G., and Katagiri, F. (2003). Quantitative nature of Arabidopsis responses during compatible and incompatible interactions with the bacterial pathogen *Pseudomonas syringae*. Plant Cell 15: 317–330.

160. Taylor-Teeples, M., Lin, L., de Lucas, M., Turco, G., Toal, T.W., Gaudinier, A., Young, N.F., Trabucco, G.M., Veling, M.T., Lamothe, R., Handakumbura, P.P., Xiong, G., Wang, C., Corwin, J., Tsoukalas, A., Zhang, L., Ware, D., Pauly, M., Kliebenstein, D.J., Dehesh, K., Tagkopoulos, I., Breton, G., Pruneda-Paz, J.L., Ahnert, S.E., Kay, S.A., Hazen, S.P., and Brady, S.M. (2015). An Arabidopsis gene regulatory network for secondary cell wall synthesis. Nature 517: 571–575.

161. ter Kuile, B.H., and Westerhoff, H.V. (2001). Transcriptome meets metabolome: hierarchical and metabolic regulation of the glycolytic pathway. FEBS Lett. 500: 169–171.

162. Thomma, B.P.H.J., Nelissen, I., Eggermont, K., and Broekaert, W.F. (1999). Deficiency in phytoalexin production causes enhanced susceptibility of *Arabidopsis thaliana* to the fungus *Alternaria brassicicola*. Plant J. 19: 163–171.

163. Tohge, T., and Fernie, A.R. (2012). Co-expression and co-responses: within and beyond transcription. Front. Plant Sci. 3: 248.

164. Tohge, T., Scossa, F., and Fernie, A.R. (2015). Integrative approaches to enhance understanding of plant metabolic pathway structure and regulation. Plant Physiol. 169: 1499–1511.

165. Toufighi, K., Brady, S.M., Austin, R., Ly, E., and Provart, N.J. (2005). The botany array resource: e-northerns, expression angling, and promoter analyses. Plant J. 43: 153–163.

166. Vermeirssen, V., De Clercq, I., Van Parys, T., Van Breusegem, F., and Van de Peer, Y. (2014). Arabidopsis ensemble reverse-engineered gene regulatory network discloses interconnected transcription factors in oxidative stress. Plant Cell 26: 4656–4679.

167. Waddington, C.H. (1942). Canalization of development and the inheritance of acquired characters. Nature 150: 563–565.

168. Wagner, D., Sablowski, R.W., and Meyerowitz, E.M. (1999). Transcriptional activation of APETALA1 by LEAFY. Science 285: 582–584.

169. Wang, X., Niu, Q.W., Teng, C., Li, C., Mu, J., Chua, N.H., and Zuo, J. (2009). Overexpression of PGA37/MYB118 and MYB115 promotes vegetative-to-embryonic transition in Arabidopsis. Cell Res, 19: 224.

170. Weickert, M.J., and Adhya, S. (1993). The galactose regulon of *Escherichia coli*. Mol. Microbiol. 10: 245–251.

171. Weng, J.K., Philippe, R.N., and Noel, J.P. (2012). The rise of chemodiversity in plants. Science 336: 1667–1670.

172. William, D.A., Su, Y., Smith, M.R., Lu, M., Baldwin, D.A., and Wagner, D. (2004). Genomic identification of direct target genes of LEAFY. Proc. Natl. Acad. Sci. USA 101: 1775–1780.

173. Wink, M. (2003). Evolution of secondary metabolites from an ecological and molecular phylogenetic perspective. Phytochemistry 64: 3–19.

174. Wittkopp, P.J., and Kalay, G. (2012). *Cis*-regulatory elements: molecular mechanisms and evolutionary processes underlying divergence. Nat. Rev. Genet. 13: 59–69.

175. Wright, K.M., and Rausher, M.D. (2010). The evolution of control and distribution of adaptive mutations in a metabolic pathway. Genetics 198: 483–502.

176. Xu, J., Meng, J., Meng, X., Zhao, Y., Liu, J., Sun, T., Liu, Y., Wang, Q., and Zhang, S. (2016). Pathogen-responsive MPK3 and MPK6 reprogram the biosynthesis of indole glucosinolates and their derivatives in Arabidopsis immunity. Plant Cell 28: 1144–1162.

177. Yang, F., Li, W., Jiang, N., Yu, H., Morohashi, K., Ouma, W.Z., Morales-Mantilla, D.E., Gomez-Cano, F.A., Mukundi, E., Prada-Salcedo, L.D., Velazquez, R.A., Valentin, J., Mejia-Guerra, M.K., Gray, J., Doseff, A.I., and Grotewold, E. (2017). A maize gene regulatory network for phenolic metabolism. Mol. Plant 10: 498–515.

178. Zhan, J., Li, G., Ryu, C.H., Ma, C., Zhang, S., Lloyd, A., Hunter, B.G., Larkins, B.A., Drews, G.N., Wang, X., and Yadegari, R. (2018). Opaque-2 regulates a complex gene network associated with cell differentiation and storage functions of maize endosperm. Plant Cell 30: 2425–2446.

179. Zhang, Y., Li, B., Huai, D., Zhou, Y., and Kliebenstein, D.J. (2015). The conserved transcription factors, MYB115 and MYB118, control expression of the newly evolved benzoyloxy glucosinolate pathway in *Arabidopsis thaliana*. Front. Plant Sci. 6: 343.

180. Zhao, Y., Hull, A.K., Gupta, N.R., Goss, K.A., Alonso, J., Ecker, J.R., Normanly, J., Chory, J., and Celenza, J.L. (2002). Trp-dependent auxin biosynthesis in Arabidopsis: involvement of cytochrome P450s CYP79B2 and CYP79B3. Genes Dev. 16: 3100–3112.

181. Zheng, Y., Ren, N., Wang, H., Stromberg, A.J., and Perry, S.E. (2009). Global identification of targets of the *Arabidopsis* MADS domain protein AGAMOUS-Like15. Plant Cell 21: 2563–2577.

182. Zheng, Z., Qamar, S.A., Chen, Z., and Mengiste, T. (2006). Arabidopsis WRKY33 transcription factor is required for resistance to necrotrophic fungal pathogens. Plant J. 48: 592–605.

183. Zhong, R., and Ye, Z.-H. (2012). MYB46 and MYB83 bind to the SMRE sites and directly activate a suite of transcription factors and secondary wall biosynthetic genes. Plant Cell Physiol. 53: 368–380.

184. Zhou, J., Lee, C., Zhong, R., and Ye, Z.-H. (2009). MYB58 and MYB63 are transcriptional activators of the lignin biosynthetic pathway during secondary cell wall formation in *Arabidopsis*. Plant Cell 21: 248–266.

185. Zipfel, C., and Felix, G. (2005). Plants and animals: a different taste for microbes? Curr. Opin. Plant Biol. 8: 353–360.

