## Supplementary Figures, Tables, Files for "Hierarchical and dynamic regulation of defense-responsive specialized metabolism by WRKY and MYB transcription factors"

Affiliation:
(1) Department of Molecular, Cellular & Developmental Biology, Yale University, Kline Biology Tower 734, 219 Prospect St., New Haven, CT 06511

Present address:
(2) Seeds Research, Syngenta Crop Protection, 9 Davis Dr., Durham, NC 27703

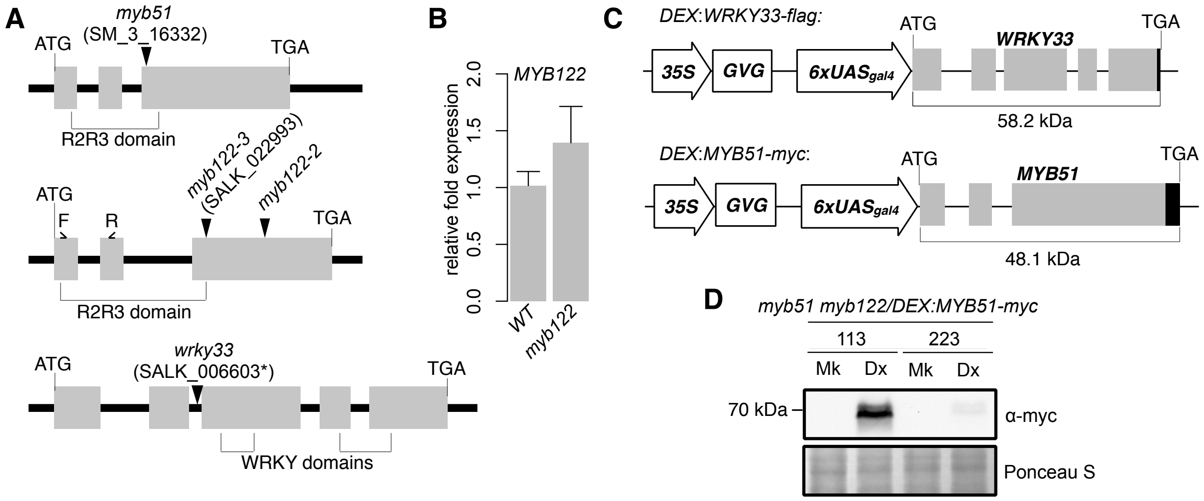


**Supplemental Figure 1.** MYB51 and MYB122 are necessary for 4OH-I3M and 4M-I3M biosynthesis in ETI (supports Figure 1).

**(A)** Schematic representation of the *MYB51*, *MYB122*, and *WRKY33* genes (top to bottom). Indicated in brackets are regions corresponding to the R2R3-MYB DNA-binding domain. The *myb122-2* insertion (Frerigmann *et al.*, 2014a) is shown in relation to the *myb122-3* insertion line used in this study. Boxes indicate exons; triangles, positions of T-DNA insertions. Arrows in *MYB122* indicate forward (F) and reverse (R) primers used for qPCR analysis in **(B)**. Asterisk denotes insertion position of an additional tandem copy of *WRKY33* and two T-DNAs (Zheng *et al.*, 2006).

**(B)** qPCR analysis of *MYB122* transcript in seedlings elicited with 1 µM flg22 for 6 hr. Expression values were normalized to that of *EIF4A1* and relative to that of WT.

**(C)** Schematic representation of the *DEX:WRKY33-flag* and *DEX:MYB51-myc* constructs. Arrows indicate promoter elements, white box indicates the constitutively expressed glucocorticoid-regulated transcription factor (GVG), gray boxes indicate gene exons, and black boxes at 3' ends of genes indicate the 1x *flag* and 6x *c-Myc* epitope for *WRKY33* and *MYB51*, respectively. Not drawn to scale.

**(D)** Immunoblot analysis of MYB51-myc protein expression in 9-day-old *myb51 myb122/DEX:MYB51-myc* seedlings co-elicited with 20 µM Dex (Dx) or mock (0.5% DMSO) (Mk) and *Psta* for 6 hr. Immunoblot analysis of WRKY33-flag protein expression in 9-day-old *wrky33/DEX:WRKY33-flag* plants was reported in Barco *et al.* (2019b).

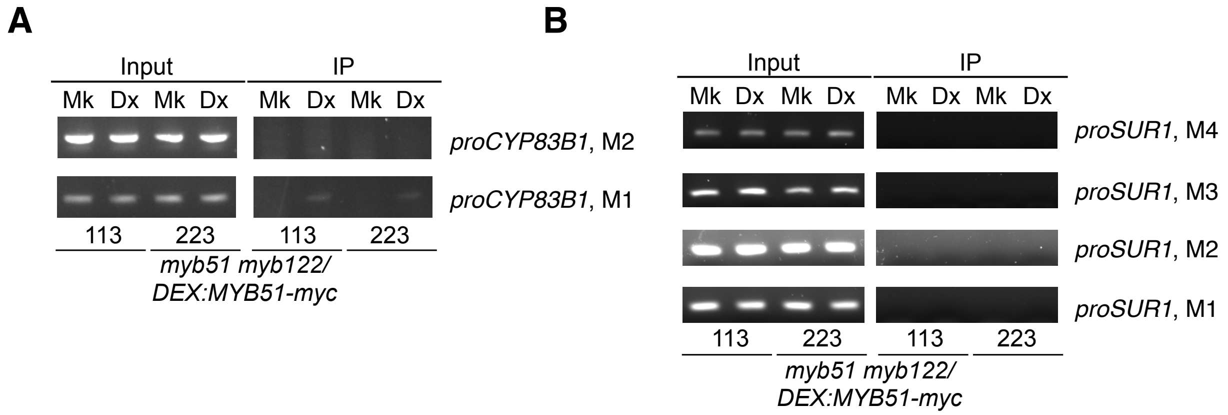


**Supplemental Figure 2.** MYB51 directly activates SMRE-containing *CYP83B1* promoter (supports Figure 2).

**(A)** and **(B)** ChIP Gel images of amplicons of *CYP83B1* (**A**) and *SUR1* (**B**) promoter (pro) regions bound by MYB51-myc in 9-day-old *myb51 myb122/DEX:MYB51-myc* seedlings co-elicited with 20 µM Dex (Dx) or mock solution (0.5% DMSO) (Mk) and *Psta* for 9 hr.

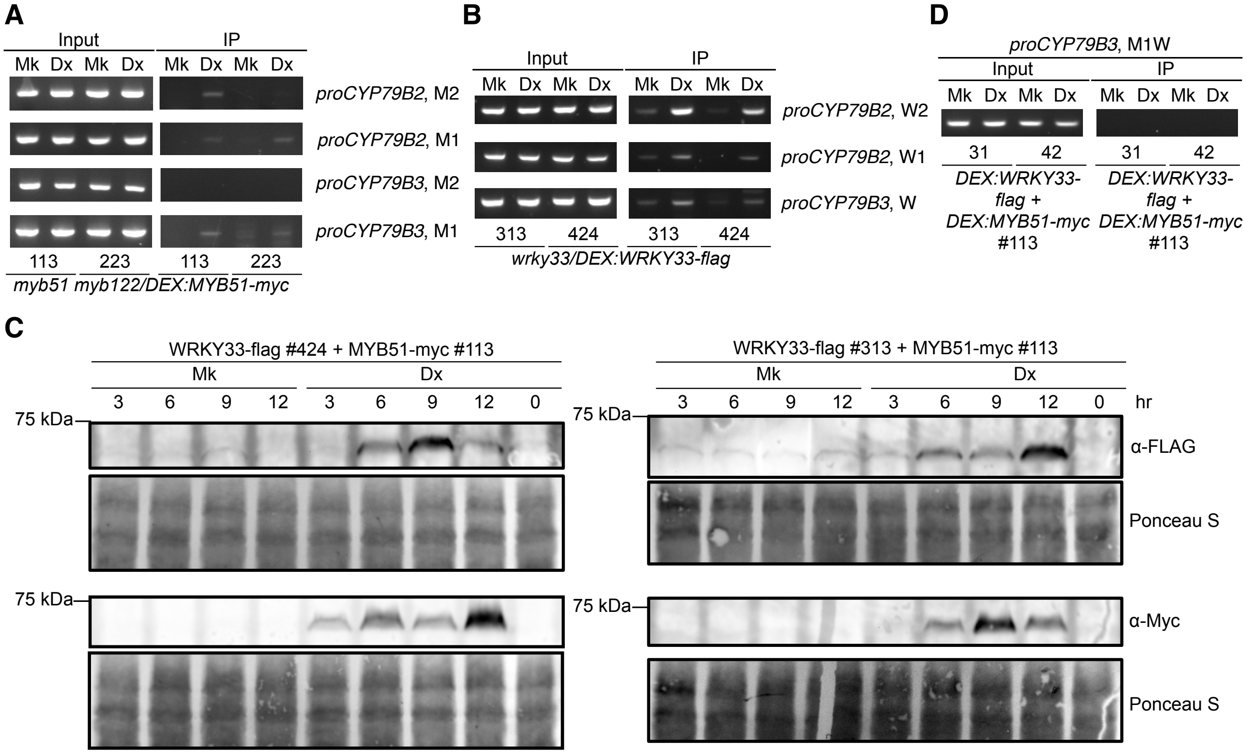


**Supplemental Figure 3.** MYB51 and WRKY33 directly activate SMRE- and W-box-containing *CYP79B2* and *CYP79B3* promoters (supports Figure 3).

**(A)**, **(B)**, and **(D)** Single ChIP (**A**) and (**B**) or sequential ChIP (**D**) gel images of amplicons of *CYP79B2* and *CYP79B3* promoter (pro) regions bound by MYB51-myc **(A)**, WRKY33-flag **(B)**, or both **(D)** in 9-day-old seedlings co-elicited with 20 µM Dex (Dx) or mock solution (0.5% DMSO) (Mk) and *Psta* for 9 hr.

**(C)** Immunoblot time-course analysis of WRKY33-flag (top) and MYB51-myc proteins (bottom) in 9-day-old seedling sequential ChIP lines containing both *DEX:WRKY33-flag* (#424 or #313) and *DEX:MYB51-myc* (#113) transgenes, and co-elicited with 20 µM Dex or mock (0.5% DMSO) and *Psta* for the timepoints indicated.

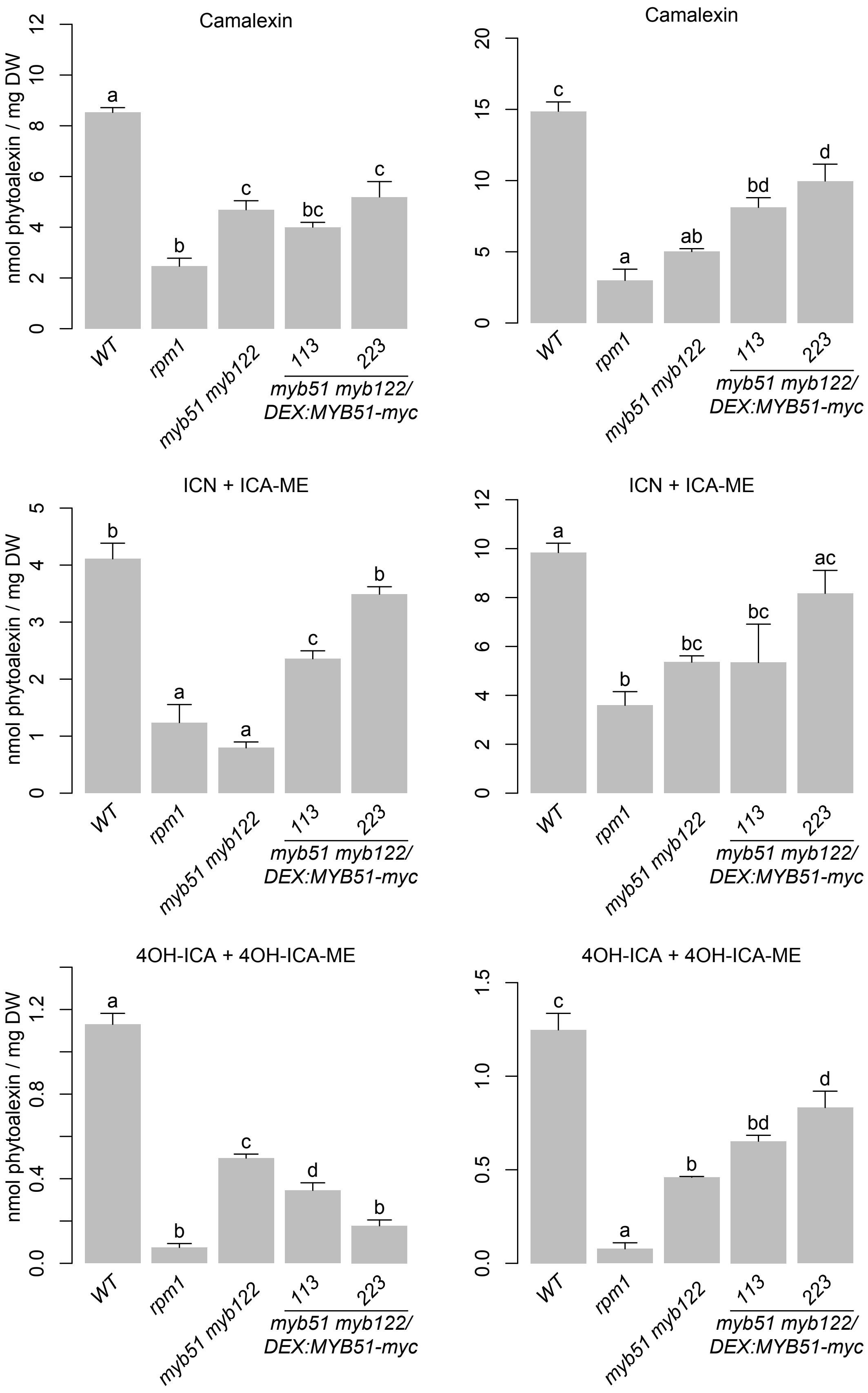
Camalexin
nmol phytoalexin / mg DW
0
2
4
6
8
10
12
a
b
c
bc
c
WT
rpm1
myb51 myb122
113
223
myb51 myb122/
DEX:MYB51-myc
Camalexin
0
5
10
15
20
c
a
ab
bd
d
WT
rpm1
myb51 myb122
113
223
myb51 myb122/
DEX:MYB51-myc
ICN + ICA-ME
nmol phytoalexin / mg DW
0
1
2
3
4
5
b
a
a
c
b
WT
rpm1
myb51 myb122
113
223
myb51 myb122/
DEX:MYB51-myc
ICN + ICA-ME
0
2
4
6
8
10
12
a
b
bc
bc
ac
WT
rpm1
myb51 myb122
113
223
myb51 myb122/
DEX:MYB51-myc
4OH-ICA + 4OH-ICA-ME
nmol phytoalexin / mg DW
0.0
0.4
0.8
1.2
a
b
c
d
b
WT
rpm1
myb51 myb122
113
223
myb51 myb122/
DEX:MYB51-myc
4OH-ICA + 4OH-ICA-ME
0.0
0.5
1.0
1.5
c
a
b
bd
d
WT
rpm1
myb51 myb122
113
223
myb51 myb122/
DEX:MYB51-myc

**Supplemental Figure 4.** Variable levels of camalexin and 4OH-ICN observed in *MYB51-myc* lines (supports Figure 4).

LC-DAD analysis of camalexin (top), ICN (center), and 4OH-ICN (bottom) in 9-day-old seedlings co-elicited with 20 µM Dex and *Psta* for 24 hr. Data represent mean ± SE of four replicates of 13-17 seedlings each. The *rpm1* mutant is ETI-deficient when elicited with *Psta* (Bisgrove *et al.,* 1994; Barco *et al*., 2019b). DW, dry weight. Different letters denote statistically significant differences ($P < 0.05$, one-factor ANOVA coupled to Tukey's test).

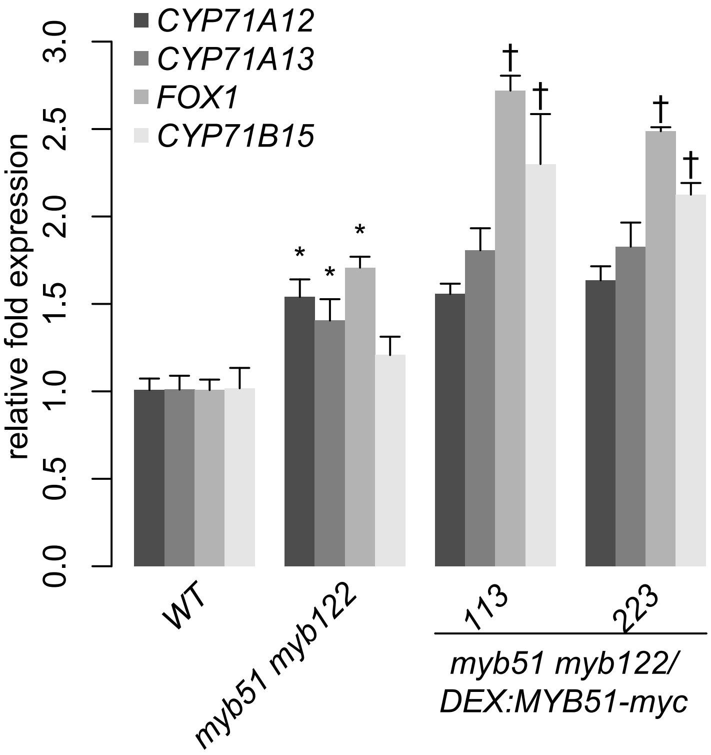


**Supplemental Figure 5.** MYB51 and MYB122 contribute to camalexin and 4OH-ICN biosynthesis (supports Figure 4).

qPCR analysis of camalexin and ICN biosynthetic genes *CYP71A12/CYP71A13*, ICN biosynthetic gene *FOX1*, and camalexin biosynthetic gene *CYP71B15* in 9-day-old seedlings co-elicited with 20 µM Dex and *Psta* for 12 hr. Data represent the mean ± SE of four replicates of 13-17 seedlings each. Expression values were normalized to that of *EIF4A1* and relative to that of WT. Asterisks and daggers denote statistically significant differences compared to wild-type and myb51 myb122, respectively ($P < 0.05$, two-tailed $t$ test).

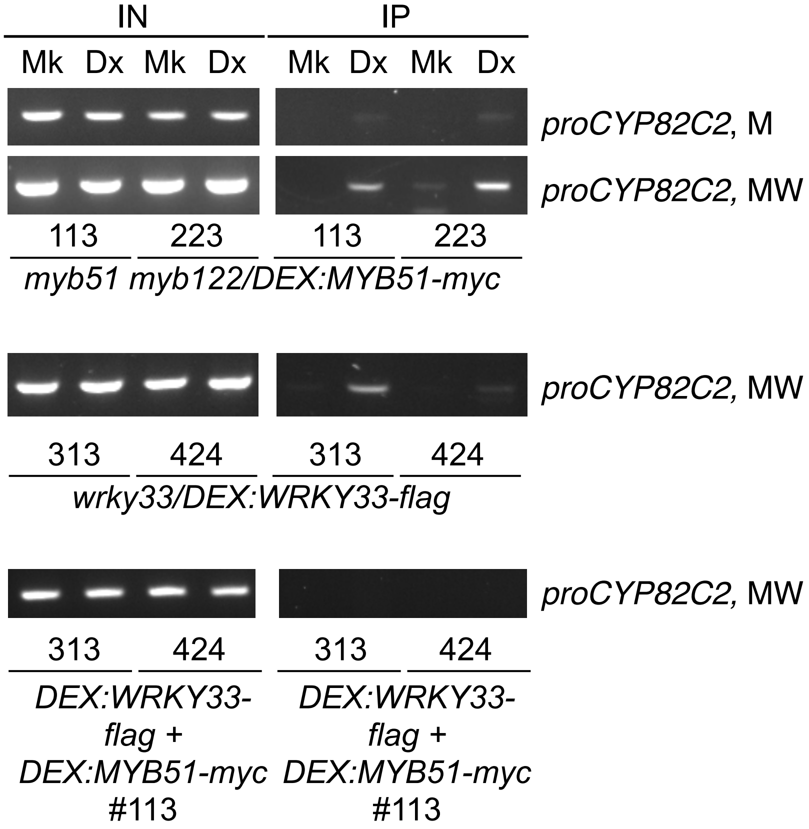


**Supplemental Figure 6.** WRKY33 and MYB51 do not co-localize to W-box and SMRE-containing *CYP82C2* promoter region (supports Figure 5).

Sequential ChIP gel images of amplicons of *CYP82C2* promoter (pro) regions bound by MYB51-myc (top), WRKY33-flag (middle), or both (bottom) in 9-day-old seedlings co-treated with 20 µM Dex (Dx) or mock solution (0.5% DMSO) (Mk) and *Psta* for 9 hr.

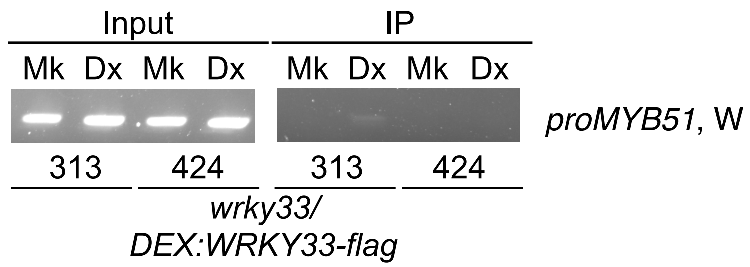


**Supplemental Figure 7**. WRKY33 binds W-box-containing *MYB51* promoter (Supports Figure 6).

Gel images of amplicons of W-box-containing *MYB51* promoter (pro) region W bound by WRKY33-flag in 9-day-old *wrky33/DEX:WRKY33-flag* seedlings co-treated with 20 µM Dex (Dx) or mock solution (0.5% DMSO) (Mk) and *Psta* for 9 hr.

| | details | Denoux et al., | Denoux et al., | AtGen_B-34_ | AtGen_B-41_ | AtGen_A-41_ | AtGen_A-45_ | AtGen_A-50_ |
|---|---|---|---|---|---|---|---|---|
| | tissue | seedling | seedling | leaf | leaf | leaf | leaf | leaf |
| | treatment | Flg22 | Flg22 | flg22 | flg22 | Psy phaseo | Psy phaseo | Psy phaseo |
| | time point (hr) | 1 | 3 | 1 | 4 | 2 | 6 | 24 |
| MYB34 | Fold Change | -3.476 | -5.446 | 0.65497275 | 0.63168124 | 1.02764538 | 0.47731239 | 0.3575506 |
| | P-value | 0.00008 | 2.34E-08 | 0.10551144 | 0.07098809 | 0.95426352 | 0.04786825 | 0.00211221 |
| MYB51 | Fold Change | 47.14094 | 23.83079 | 2.04692388 | 1.81469649 | 2.3128655 | 4.72981248 | 1.58467928 |
| | P-value | 1.21E-28 | 8.92E-33 | 0.06113077 | 0.10131137 | 0.0060054 | 0.00775587 | 0.00013798 |
| MYB122 | Fold Change | 3.23323 | 13.76 | 0.24338624 | 2.21568627 | 4.77142857 | 1.8984127 | 7.78 |
| | P-value | 0.15382 | 8.30E-10 | 0.34631054 | 0.31711367 | 0.17470999 | 0.55714933 | 0.15493082 |
| WRKY33 | Fold Change | 23.453 | 11.841 | 2.82124053 | 2.74755927 | 2.19262736 | 3.2795076 | 2.31010374 |
| | P-value | 1.03E-27 | 4.81E-18 | 0.00070675 | 0.00655808 | 0.00114146 | 0.00949111 | 0.0067449 |

| AtGen_A-19_ | AtGen_A-24_ | AtGen_A-25_ | 2530_Psy-ES | 2529_Psy-ES | 2791_Psy-ES | 2788_Psy-ES | 2525_Psy-ES | 2796_Psy-ES |
|---|---|---|---|---|---|---|---|---|
| leaf | leaf | leaf | leaf | leaf | leaf | leaf | leaf | leaf |
| Psy avrRpm1 | Psy avrRpm1 | Psy avrRpm1 | Psm | Psm | Psm | Psm | Psm | Psm avrRpt2 |
| 2 | 6 | 24 | 4 | 8 | 16 | 24 | 48 | 4 |
| 0.54146806 | 2.59205934 | 0.73326414 | 1.07845157 | 0.85291707 | 0.70794421 | 0.73462239 | 0.63225458 | 0.79323923 |
| 0.20422927 | 0.05865677 | 0.39650357 | 0.88062399 | 0.82935231 | 0.07459958 | 0.5338396 | 0.20989059 | 0.70129971 |
| 3.1031746 | 1.32759281 | 1.38901707 | 0.69241825 | 1.26021374 | 1.94502099 | 2.66834747 | 1.81109017 | 1.791178 |
| 0.09551394 | 0.50825441 | 0.20932129 | 0.10288338 | 0.1394821 | 0.11064329 | 0.03636376 | 0.46353279 | 0.18876567 |
| 8.08571429 | 4.71746032 | 36.22 | 2.16575592 | 6.50877193 | 1.46428571 | 0.95544193 | 0.49234973 | 0.34146341 |
| 0.38940782 | 0.00760971 | 0.02623886 | 0.0009021 | 0.01976948 | 0.6218737 | 0.94155251 | 0.22346208 | 0.05168994 |
| 2.19164956 | 2.7188028 | 3.28236355 | 1.18682779 | 1.18168181 | 1.99746173 | 2.54292216 | 1.74664892 | 1.20710287 |
| 0.00337224 | 0.00820789 | 0.02255407 | 0.23178077 | 0.00610072 | 0.13172359 | 0.00644129 | 0.44402462 | 0.07381242 |

| 2793_Psy-ES | 2508_Psy-ES | 2509_Psy-ES | 2784_Psy-ES4326avrRpt2-48 [ATGE_ExpID_168] Pseudomonas half leaf injection le |
|---|---|---|---|
| leaf | leaf | leaf | leaf |
| Psm avrRpt2 | Psm avrRpt2 | Psm avrRpt2 | Psm avrRpt2 |
| 8 | 16 | 24 | 48 |
| 0.77356382 | 0.49714979 | 0.67044977 | 0.73667745 |
| 0.38270176 | 0.10438865 | 0.04540329 | 0.28107588 |
| 1.34405024 | 1.12047001 | 0.85188685 | 0.93090946 |
| 0.06145988 | 0.00633511 | 0.13205375 | 0.27219847 |
| 0.19901168 | 1.19105691 | 0.24617737 | 0.7891232 |
| 0.30716202 | 0.64233625 | 0.17206966 | 0.09972893 |
| 0.93765349 | 0.76800801 | 0.83610537 | 1.02457985 |
| 0.0569519 | 0.07612223 | 0.02863455 | 0.26841229 |

**Supplemental Table 1.** Published transcriptional responses to PTI elicitors (Denoux et al., 2008) and ETI elicitors (Toufighi *et al.,* 2005; Austin *et al.,* 2016).

| type | TAIR ID | primers | sequence | alias | approximate upstream location (bp) |
|---|---|---|---|---|---|
| ChIP-PCR | AT4G31500 | 83B1pr1F | atggaggttctggt | M2 | -1900 |
| ChIP-PCR | AT4G31500 | 83B1pr1R | acggacgacgaa | M2 | -1900 |
| ChIP-PCR | AT4G31500 | 83B1pr2F | tgtcacctgtcaaa | M1 | -100 |
| ChIP-PCR | AT4G31500 | 83B1pr2R | tacttaagtctttgtg | M1 | -100 |
| ChIP-PCR | AT2G20610 | SUR1pr4F | agctttcagtaaat | M4 | -1500 |
| ChIP-PCR | AT2G20610 | SUR1pr4R | tctacatttgcatctt | M4 | -1500 |
| ChIP-PCR | AT2G20610 | SUR1pr3F | atcgtcctcccagc | M3 | -1300 |
| ChIP-PCR | AT2G20610 | SUR1pr3R | actttttgaactaca | M3 | -1300 |
| ChIP-PCR | AT2G20610 | SUR1pr2F | cactgacgcccat | M2 | -300 |
| ChIP-PCR | AT2G20610 | SUR1pr2R | tgataagaatagtt | M2 | -300 |
| ChIP-PCR | AT2G20610 | SUR1pr1F | ccttcacaaagcc | M1 | -200 |
| ChIP-PCR | AT2G20610 | SUR1pr1R | aagttttgaattatg | M1 | -200 |
| ChIP-PCR | AT4G39950 | 79B2pr1F | agattcaaaatgtt | W2 | -2300 |
| ChIP-PCR | AT4G39950 | 79B2pr1R | aaaagtcaatttcc | W2 | -2300 |
| ChIP-PCR | AT4G39950 | 79B2pr3F | gtactgtcttgaaa | M2 | -2000 |
| ChIP-PCR | AT4G39950 | 79B2pr3R | gattggtttgtaata | M2 | -2000 |
| ChIP-PCR | AT4G39950 | 79B2pr2F | ccttgatttggaaa | W1 | -1200 |
| ChIP-PCR | AT4G39950 | 79B2pr2R | ctgatgtgggacg | W1 | -1200 |
| ChIP-PCR | AT4G39950 | 79B2pr4F | gcgacaatactag | M1 | -600 |
| ChIP-PCR | AT4G39950 | 79B2pr4R | ggaaatacgcaa | 1M | -600 |
| ChIP-PCR | AT2G22330 | 79B3p2F | tatgaagataaatc | M2 | -500 |
| ChIP-PCR | AT2G22330 | 79B3p2R | ggtagtctctttatc | M2 | -500 |
| ChIP-PCR | AT2G22330 | 79B3pr1F | gatttttggcaggtc | W | -300 |
| ChIP-PCR | AT2G22330 | 79B3pr1R | aacttggtctctag | W | -300 |
| ChIP-PCR | AT2G22330 | 79B3p4F | gtacatttttgaaa | M1W | -250 |
| ChIP-PCR | AT2G22330 | 79B3p4R | ggaagcagtgtgg | M1W | -250 |
| ChIP-PCR | AT2G22330 | 79B3p3F | catttcacacgtac | M1 | -100 |
| ChIP-PCR | AT2G22330 | 79B3GW1R (g | GGGGACCAC | M1 | -100 |
| ChIP-PCR | AT4G31970 | 82C2pr9F | gttttgagtcttcgtt | M | -3000 |
| ChIP-PCR | AT4G31970 | 82C2pr9R | attggaagcttctc | M | -3000 |
| ChIP-PCR | AT4G31970 | 82C2pr8F | taggcgtatgtctc | MW | -100 |
| ChIP-PCR | AT4G31970 | 82C2pr8R | cgtgcaaaagag | MW | -100 |
| ChIP-PCR | AT1G18570 | MYB51pr2F | attcattctgtcatg | W | -1150 |
| ChIP-PCR | AT1G18570 | MYB51pr2R | aaaaaagctttatt | W | -1150 |

| type | TAIR ID | primers | sequence | efficiency |
|---|---|---|---|---|
| qPCR | AT3G13920 | EIF4AF | TCTGCACCA | 100 |
| qPCR | AT3G13920 | EIF4A2R | TCATAGGAT | 100 |
| qPCR | AT5G60890 | MYB34_1F | ATGGTGAGG | 85.9 |
| qPCR | AT5G60890 | MYB34_1R | ACATCTCTT | 85.9 |
| qPCR | AT1G18570 | 513F | ACAAATGGT | 73.1 |
| qPCR | AT1G18570 | 512R | CTTGTGTGT | 73.1 |

| | | | | |
|---|---|---|---|---|
| qPCR | AT1G74080 | MYB122_FB | GCATGGACT | 100 |
| qPCR | AT1G74080 | MYB122_RB | GTCAGGTCT | 100 |
| qPCR | AT2G38470 | WRKY33_1F | GAAGCAAAG | 99.4 |
| qPCR | AT2G38470 | WRKY33_1R | CTACGATTC | 99.4 |
| qPCR | AT4G31500 | 83B1_1R | TCACGCCAT | 87.3 |
| qPCR | AT4G31500 | 83B1_2F | TGGACGTCA | 87.3 |
| qPCR | AT2G20610 | SUR1_F | TGTAACACC | 91.5 |
| qPCR | AT2G20610 | SUR1_R | CCTGTATTG | 91.5 |
| qPCR | AT4G39950 | 79B2_1R | GTAACTTCG | 95 |
| qPCR | AT4G39950 | 79B2_2F | TCGCCGGAT | 95 |
| qPCR | AT2G22330 | 79B3_1R | AGTCACTTC | 92.8 |
| qPCR | AT2G22330 | 79B3_2F | TCGCAGGTT | 92.8 |
| qPCR | AT4G31970 | 82C2_L1 | CAAGCATGT | 91.6 |
| qPCR | AT4G31970 | 82C2_R1 | GCATCTTCA | 91.6 |
| qPCR | AT2G30750 | 71A12R | GATTATCAC | 93.1 |
| qPCR | AT2G30750 | 71A12F | CCACTAATA | 93.1 |
| qPCR | AT2G30770 | 71A13F | GCTTCGGTT | 87.5 |
| qPCR | AT2G30770 | 71A13R | ATTGATTAT | 87.5 |
| qPCR | AT1G26380 | FOX1_1F | ATGATGGAT | 98.5 |
| qPCR | AT1G26380 | FOX1_2R | ACTCAGGTT | 98.5 |
| qPCR | AT3G26830 | PAD3_L1 | ACGAGCATC | 78.4 |
| qPCR | AT3G26830 | PAD3_R1 | TCGGTCATT | 78.4 |

**Supplemental Table 2.** qPCR and ChIP-PCR primer sequences and efficiencies.

**W2, M2, W1, M1 promoter regions of *CYP79B2***

Agttagaatcgaattaatccaaaaggaattattgtggaattaactaaagcaatccatacaaatc
acaacaaaagctcgtcaagtatatataatcagaat**agtcaa(Wbox)**taatttagc**tttgaat(**
**WRKY33)**gccgaac**ttgact(Wbox)**aaaactcagacatgttatagtaccaaaatatttgtaat
ttagggaatatcattgctaatacaatataaaaaagatcatagctag**attcaaa(WRKY33)**atg
ttgaacatgacctacgacatt**gttgaat(WRKY33)**tcattgatctggtcttgctaaaacttt
aaaattgatgagttcaacatcttcaaatgcatgataacgggtccaacggaaa**ttgact(Wbox)**
tttttttcatgctcctgatatataataatatctaacgattacgggttccactaattgtcattac
tcattaacattcctatttaaaagttgtgatagttttagggttttacgtagtcgtgtcatatagc
gattaactacgtacttgtagatttatcaattacttctgttgtttacgagaacctaaaaaaaaga
agcagatgcctagtttatagagcacgtgtactgtcttgaaaactta**ggtaggt(SMRE8/ACI)**
tggtaaggttaccaaaagaccttaaaggaatataaagttactaattaacttaagtaa**agttggt**
**(SMRE2)**attgcttatatattgcaaagtattacaaaccaatcccctctgtatattgttttaaac
catagatttttttacaattaagtttatgatcaatcaattatttcaccatttctattaaattatg
taaaaagaaaaggatatatatatatatatataattaaataagaataaatcaaaataccgaaatt
ttttattatccattctttgtggacatcgcccctaatatataaaaaaaaaaaactttcgtataac
tgatttatattttttgtaaaaacttaaaggaagcctaagaaatatcttgtgatattttgaca
aaatgtggtatatatcttttataatatcatttataaagaaaatattgattacatggtgaaaaa
cattttgctagcgatcaacaaaattaaataggcacatgttaactgatctcatacgaccttgaaa
ttttaatctttgtgtcgagagaccgatctttatgcaaattatgaaactacacatggtttatgca
cggaagatcacattgcatgtataccatattataaaccaaaaatgatcaagaagaaggcgaaaac
atttgggtaaattttaaatttcgatcatgcgatttttttagctcatcatcaacagacaagaaact
atcttttgtactgtaaatactaaatacaaaataaaatcttcatcatttttttgcatgtgtcaaat
aaattacgcgaacttttttttttttatcgactattaatagagaaacctgtttttatttgccttgat
ttggaaaaatggagaaa**ttgact(Wbox)**taagacttagtctcggtcacatcggcaacaacgga
gcttaaacggcgtccgcaacatggaaactcaagccacgaatctgatata**ttgact(Wbox)**ata
gaagtagtaagtaact**ttgact(Wbox)**cgtccCacatcagtttcaatttccacgagggtattt
ggcaggtgaactctctacgtacccaaaacataatggctattttatttcataactgatatttagc
aattaattattcgtcctttttaaaccaatttctatagttgggaaaataatcaatttttacactt
tcaatgtatacgttacagatttttttttattagtcatgcacatattttcaattttttacactttc
aatgtaaacaatcgattcttaattgttaaaaataggtttacgtaaggaattaaagatttgttta
aaatatgttccggccggtctaataatttacttgacgttaatttcttaaacacttttagatagga
ggctttgtttatcccaaatgattttgtaccactgcgacaatactagctagacataaaatgttaa
taaattttttattaagtaatataatcgaagtattagatcaatgtagtagaca**gttaggt(SMRE7**
**/ACIII)**taactaaaacaagagtaaacactttttttttttcttttcaggataggtaaaacaaatt
tcacactattttgcgtatttccttaaatttgttgttcgttttctcagcaaagatgaatatttttg
tttcatagtaattcacaagtataaactcgccagaactcctcaaacagtgaaatataatatagct
tttaactgtttttcggctggaccgggtttttaagtgcatatataacacgaggaattttggcagg
tcaccaacaaaactttaaaaatattaaaaattcccatcaagaatagaaattaataaacaatga
tatctctaataatatagatattttgaaacgttaggaataatcgtaataatgttcaacgttggtg
gtggtactcaagatggaccctccctcccacattttcctcactccttcgtaagtcctttccacgc
ataagggtattatagtcatttcacataaactaacgactactagacttgtatataaataggaagg
tgaagctctctctttatccatgcagagacaacagaaaccacaacaaaaactttgagtcctcttc
ttctctatacacaaac**ATG(TSS of CYP79B2)**

**M2, W, *M1W*, and M1 promoter regions of *CYP79B3***

atataatttcaaatttacacattgtataacgatatgagatttacataaaatattttcatatatt
tgttcattttaagagttcacaccacggataaattatcgatatataggaaagacttggctataac
tacaaaaagctgaagggacatttattattatccccaaaaaaagcaacatcatgttcgtattga
ttattgaatatgacatattgtagagacgaatcatttatgaagataaatcaaacg**acctacc(SM
RE8/ACI)**aaaataagatatactcaaaagagatataatgtaagttcaatcagttaaagctacta
ttatggctctgttagatatgagagtagtgctaagataaagagactaccaattcatattaccgag
aaaaaaaaaaactgtcttgttagtttagagtacatatataacacgaggattttggcaggtcac
caacaaaataattttaaaatatattatgaatgaattattatacatgattattatcttttat*gta
cattttttgaaagttagccctacaaa****attcaaa(WRKY33)****atatggtgacactcctcactaga
gacca****agtt****ggt(SMRE2)**ccaactctagctataccttccacactgcttcc*tcccccaccacaa
aaggggtactatcgtcatttcacacgt**acctacc(SMRE8/ACI)**tgtaattaatgtgacatta
taaataacaacgtgaaagaaggatccatgcagaaacaattagcatatattacaaaacacatact
aattgtttctccttctccttcttccttgcaa**ATG(TSS of CYP79B3)**

**M and MW promoter region of *CYP82C2***
aggcctttgtctttgttactcacgacactgtcgttttgagtcttcgtttcgacttctccaaagg
gatcttgaagctgtctggattcagatcttgaaggtgatgtcggagcagtggaggtcatgctaat
atcaaca**accaaac(SMRE3)**ctgtaagcatatgagaagcttccaataaaacattcttaaaatt
tattttcaacaactgccaatgtattaaaaaaaagcaatctcatgaattgcctacacgctaatta
ggcgggcgaaatttgttttatgtgtatgtggtatggtatttgtttttcctcgccatcctttcct
ggtaaaaccaataatttttatatcacatataaagttacgtgctgaacgaaacaaccgattatt
attaagaatccatgttgaaatagagaaaatggattgtgtgaatagaaaaacaaaaacgaaaagt
cgacaaaaaaaacaaaaaacagaaaaacccactaa**agtcaa(Wbox)**gcgttttgtttcaacca
tgttatagtttagaactttataatcagaaattaattccacgaaaatgaataaatgtcagc**agtc
aa(Wbox)**accagaccattaaaagaaacaattaccattaaaagatata**ttgact(Wbox)**aaca
aaatcaatacaaacctttgcgtccatctctgttatggacacagttgacaattacaaaaaccctc
catcttcctgtctctctctatcatccatgactttttttttttctcaacaacaactaattattatc
caaagttcttagataatgagattaatacatgctatttcttgcccaactaggcacaatagaataa
agcttaggatcata**attcaaa(WRKY33)**aatgaaagagactcccttgatactctgccaatgtg
tgccctctcttggtatgaacacaaac**ttgacc(Wbox)**tcttcaaattgggggagcaacctctg
cagatcatccatgacttaaactgcacaataattaaaataggaaacttaagttcatcaagagaga
ccaaaaatatcacaaaactcttatttataccatttctattggcaatatttaatagtttttata
attaaaataataaaataaaagagaaaaaaagtaaaaaaattctcgcttgaatttttttgaaata
ctctatacaactcatagtgacatttttttttttcaattggttcaatttttttttactaaaaaaaa
aattttaattactttttttagaaattctttggggttctccttgatggaaaaagatataccaaaat
aggtaaagattcttgaacaatggtgtaccatctacgacttcaccgaagtgagctaaagcatccc
tttaacgttggtaatcagttacccatgacacacactggacatttaccttcatcactctggtcgt
cctgagacttgaacagtttcttgtttctgttatcatttacaacataaggcccacttcttgccca
cacacagtttattactcatcaatttctctaatatactattactctataacaacaaaaaagtaac
attgtaatttgtacacatagcgtttgatctagctagttctatatgtagaattgatgttagcaag
tgaatcaaaattttgcggagatgatgtttgaacatgggtttgttcattgtgttacccacacgaa
gccaatttgattgttcgtccagatccttgtaaaaagaacacatatggatcatttagtcgcaaca

ataaagaagatgtatcagagctacatatgtatgtttcacagtatttctcctctcgattaacaatgaactggtgcaaggatcacccgattcgttcttgagggaagtatcttgggaagcagatcgatttagaggcaaagagggctcaaaagaaatcgttgatgaaatcgctcaaggtctcgcttgaaggttcgcagtttcttcactaatcgtgagcctttttaaagttgcagtaatcgttgtatgaaacaagcgtgcacggtcatagtcatgctcgttttctcgaggcagcttgacaagagttaaattatggtgatggataattagtgtatttttgaggctgtctcaactgaactttcatttaatgacttggctagcgatacagaaccgtctagccacaaaagagagattattaaagtggaatggcgaggcaaatggtggtagtgtgttttgtcaaacggaaatagaaacaagagatcacctatttttctcttgttcatactctcatgcttggtcagctcttactatgaaatatatgctcggctctcgcttcacgacttgctgtgattctcttgttgccctatctacgacctttcctctccacaacttcatttgtttgtgtttagatatgtctttcaactcactatacacaatctatggaaagaattaaatgccccataactcagagctcccacttctttttaaatcttgttctattaagttttcaaaatttctcggttttat**gttgaat(WRKY33)**ttattgtaaaaccgtt**tttgaat(WRKY33)**tttaaactcatacaaaaacaaaaaaatataattagtgtattttaaattatttgtgttttaatatatttttttattttatcttttaaaacatgtttttttatttgagttattattatgaattcagtattataaagtcatatcttcatttcaatttttt**tttgaat(WRKY33)**ataatgttatataatatttctaaacacaagtagataacgtt**ggtcaa(Wbox)**t**atttggt(SMRE1)**taagataaatggtggaaaaatattcagaaatgttcaaaaatgttcgaccatttttttttatttcaaaatgtacgtcagtaactatcgattttttt**ttgacc(Wbox)**atatacaatttgcgacccccgcctttcgacgacttgctttt**ggtcaa(Wbox)**acagcagtaagataggcgtatgtctcatgcttacatggtattgaaccgataatatgtgtgtgtatatatagagagacagactatactttttaatc**attcaaa(WRKY33)**actagaaatc**accaaac(SMRE3)**acacatctcttttgcacgctcaaaccact**ATG(TSS of CYP82C2)**

**M2 and M1 promoter regions of *CYP83B1***

aaacaaaggcttcaagaattctaccggaatcaATGGAGGTTCT**GGTAGGT(SMRE8/ACI)**TCTACGTTTAGGGATCGTAGCAGCGTCACGACGCATGATCAGGCCGTACCGGCGTCTCTGTCTAGCCGGATTGGTTTGAGGAGATGTGGTAGATCTCCTCCGCCGGAGAGTTCGTCGTCCGTTGGAGAAACCAGCGAAAACGAGGAAGATGAAGACGACGCCGTTTCGTCATCACAAGGAAGATGGCTCAATTCGTTTAGTTCATCTTTGGAAGATTCTCTTCCGATTAAgtacgtttctttattaattatcttctaattattctgttatctacacctgaatatattcgatcgatttattttattacaagaataaagtaaatcattagattctcttatgttcttgggtttgattttattgatgaatctgtaagatgttgaaattgtttcttacttattagattctcagtttaaatgttattactacatgttgtgtttttgtttagtcatgtggtttaggttttggtttttgttacagGAGAGGATTATCAAACCATTACATAGGGAAATCGAAATCGTTTGGGAATTTAATGGAAGCGAGTAACACAAACG**ATTTGGT(SMRE1)**GAAAGTGGAGAGTCCATTGAACAAGAGAAGAAGATTGCTTATTGCTAATAAGCTAAGAAGAAGATCGTCCTTGTCTTCTTTCAGCATCTACACTAAAATTAACCCTAATTCTATGCCGTTGCTTGCATTGCAAGAATCTGACAACGAGGATCATAAACTCAATGATGATGATGATGATGATGATAGTAGCAGTGATGATGAAACTAGTAAGCTGAAAGAGAAGAGGATGAAGATGACGAACCATAGAGATTTTATGGTTCCTCAGACTAAGAGCTGTTTTAGCTTAACTAGTTTTCAAGATGATGATGATCGATGAtcgatgatcattcttcttcttttgtccaagagtagataaagtttcgttgacaaataagttggaggaagatttgacagaaaccaatttacgtgtaattgtaataaattaatttgactgtttgtatattttgaatttctcaacgaagtagactttttatttttttgtcacacttcttgtcaaccttaacatttgttagtaagcccatgagattggttctataaattttaatgtccgcatgagaacaaagtcgttaatattgacggcttagtaatttcaatagcggctgcgaagtcaatagggacatctttaggctcgttctccgaaatactctttttt tttttctctaccgagttgaatttatgtacgattcatcttagattgacgaagtagaatatttgtctcactgtg**acctact(SMRE6)**gttcaactgcaaaaatatgaacatatttttctacaaaaattgactacagagtaattagaaaacttaaaaactgagagtttagtttttggttacgtatacttttttttctttcagtttacgttatagcataggagaaaatgatcgcttcaaatttcataggtatttttaggaatttttctaaagtaccatttttatgctttgtttttaaatcgatcatatcagttcaaaaagaaaaaaatcaagaatgatgtctaaatgaacaaaagtttatgagatttataaaaaacaagttgatgatattttattacttcataatatacttactgttctatagtatattttagaaaacaaaaccaagaggttatattcttatccaatatcttcacattttcaactcttcatcgatctcgctttcagtattattgtgtaacccaagttacatttttaattccaaaaaccagtagtcc**acctact(SMRE6)**aacatcacgttttaaaaacaagggaaaattctgtcacctgtcaaaaaatccaaaacaa**accaacc(SMRE4/ACII)**aataaaactttttgtcttgctatataaaccacatcatcattcaaagtagaaaagtatccgaacacaaagacttaagtaagtcaaacagaaaaaa**ATG(TTS of CYP83B1)**

**M4, M3, M2 and M1 promoter regions of *SUR1***

TTCCTTATCAAGAGTGTTTCTCTTATCTAGACTTGCAAAATCAATCAAATCTTGAGGAGAACAGAGTGTATCATCACGTCCATCAAGAAGCCTTATAACTCGTATGAGTTCACTCACAACAATGGCCACCACACATCTTACAAAACACACATAAGAAAAAAACAAAAGGAGTTAAAAAAGTGAAGAAAATATTTCAATATCATATCTTATCTATATTACAATTAGAGTAGTTGTATTGTAAACTTTTCAATATTGTTTTAAAGATTCTCATATGTACTTCATATATATGTCTCTTTATATTGCAGCATATAATAATCTTTTCTCTACAAAATATTCATTCATATGTCCATTCCTTTTTCTCTTGCAACTTAACTGTGTTCTTTCTGATCTCTTAAAGCTTTCATTATTATCCTTCAATCAGCTTTCAGTAAATCTGGGATTTTGAAATAAAACTACAAGAACAATAATAAATTGTTAG**ACCAAAC(SMRE3)**TTGTTTCATAGAGAAAAAAAAAGAATCTGATACACTT**ACCTAAC(SMRE7/ACIII)**ATGTTGACCAAGATGCAAATGTAGATTATTTAATCGTCCTCCCAGCTCTCAAAAAAAAACCCATATCTTTAGCAATTTCTTTCAAAACATGGAGAATATTTTTTCGAGCATCACTTAT**ACCAACC(SMRE4/ACII)**AGTGATTCAAGAAGCTGGTGTAGTTCAAAAAGTATAGCCCTCACTCGTCGACGAACTTTAAATGAAAGTAATGAAAATCTGGTATTAACTTGAAACTTTCACACTTCTCTTCCACTTTTATCCTTTCACACTAGAAAATTACAATGTGCACAATTATAGCCGTTTCACACTTCTCTTCTACAATTTTTTTTTCACATCAAAAAATTATTTCAGGTTTTTGACACATATTCCCAACAATTTATTTTTCTTAATAATTAATATAATTTTTAAAATATATCAATGTACAAAACATAGGCTCCAAATGGATGACAAATAATTGAATATTTGGGTGCAAATGGTTGCACAATTTAAAAGGTTGAAGAGGTTAAAAACATTATTCAATTAATCTTCAGCATTTTAGTTGAAAAAGTTGAAAAAATTATTCAATTCATTCAACTTAAAAAGTTGAAAAATGAACAAATTATAAAGGAAAAGTGTTCAACTTCTTGATCCACATTTTTGGAGATGATCATTCAACCCCAATTTTGTTCA**ACCTAAT(SMRE5)**AATTTCAACTCATTCAATTTTGAAAAACCATTTGTACAAGTTGAAGATCTTTCAACTTATTCCTCTACAAAAAGGTTAAACAATTTTTCTTTATAACTTTTTTGGTTGAATGGATGAAATTAGTTCTTAGTTATTTATCTCAAATAGTAAAAACATAAGATGAATGATCATTTAACTTTTTATAACCTCTTATTTTCAAACAACATTTGCACCCATAGTCTAAAAATAGAAATTTGTTGACCAAGAAGA**ATTTGGT(SMRE1)**ATCAAATGAATATTTTTTGTGGCTCAAACTTTTATTCTCTCTTTAAGTAGCTGAAAAATTAACGTTGGAAAGTGGGGTCTACAGACCTTTACGCCCATTTATTTACGCTACAATGGTTTTCTAATCACATCCTTAATCTTACAATTTGTTTGATAATAAATCTACTAATTAAATTTGACGTCACAAATCACAATGCACTGACGCCCATTAGCATGTTAGTGACAATGTA**ACCAAAC(SMRE3)**TGATTCATCACATTTCAAACTGAACTCATTCCGGTTGACTAATCGGTTGGGTTTTAACTCGCCAACTATTCTTATCAGGCCAGGCCCAAACATTCCTTCACAAAGCCCTGCCAATTTGATTAATTTTTGTGGCACGTGATTTTAGTTTTGT**GTTAGGT(SMRE7/ACIII)**CTCACCAAGAT

TATTGATCACGTTCCATAATTCAAAACTTTGCATTATATACCAAAAATCCAATCCGTATTGAAA
CTTCATTTGTAAAGTTTTTTTTTTTCCCAAATAGTACTTTTTTTTTGTTTGGCCTTCTTATAGA
AAGAAGAACTCAAAGCACAGAGAGAAG**ATG(TTS of SUR1)**

**W promoter region of *MYB51***

GattatttttTagtatttgtactaaagaactactgtaatttagttgtcatacttttagcatgaa
tcattgaatttagagttgggttgatgtaaaccaagctggttagctttcttcataaccgtatgta
ctttcatatcatattagagagaaaaaaaaacacaatg**ttgact(Wbox)**aattaatgggttctg
tctaactagaaaagtggaaattagttgtacgaaaaaagaaaatagcttatctctttctttgtac
cttagattttccttggtataatggatatcgatatgtccaagtccagtgtttgagtgaccaagt
tgttgctttaatttaatctagttcaaatacgactactttgtgttgtacctctctaccttacct
tcctcgaattttcaattcgtgtgctagtttgccaattgtttctttataattttgtcgtctctat
gattgtttaagcatatttaaatatttgagagaaaaaaatactaagagttcagtcgtattttgaa
aagtcctcgcactccaagttttaaa**gttgaat(WRKY33)**tcattctgtcatggaaacgtacct
ttgtggtttttcatttcgatagacatttttaactcttctgataa**attcaaa(WRKY33)**atgt
tttttacattggttaaaatttgttgccaatttcaataaagcttttttttactcgttataaaa
ttaaaaacttgctgagaaattatagtttcatatatactttggaccgac**ggtcaa(Wbox)**tacc
aaaaaactaaaataaaatttt**ggtcaa(Wbox)**gaatcg**ggtcaa(Wbox)**tacaaaatatctc
caaagatcgaagagaaagttcaacaggattttaatttaattggtgttccggacctgccggcgcc
gtattataattcttcaataaattttatatttggattgcataatcaaaactatttgcatagcag
acatagatttcaaaataaattgatacgtaattagctatttacataggcaagacaattttacgtt
acatattttgcatgcttgatattcattctttttttttagcaacctttgctatattcattctatt
ttttcccctttttttgctactatcaatatcatatcagatgtggcggttcttatgcatgggatta
tgaaaaatgatattatataatagatgtcttcgtagtattttttttattgacaatcaaaataaaat
aaattagtttaagaaagatcaaaagtctagtgtcaaaatgtccactgacaaaagttagcagttt
ttcttttatgtaataatccgattataaaaaattgcgttaaggaatcataatgagccatgtgtca
gtcatatttaaacgacgacgttggcagcagccacacgtacagcctcaggcaagcataaatagag
atgggctctttcctataagtttcccactttgcaaagtgtaacacaaacagttctctttagagaa
aaaaagattctctctataaaccttctctttgtgtctctctttgtggtaagaaaacagagcaac
tagatctctcttctcctctgctcgttcaatgttataacgatctcgagtatcacaaaagaacaag
atgtctattcgaagaatctaccgatgtctgaattctttaatcaaaagatcttgtgctcaatctc
tatgatcatacgccccatttcacactttgtttcacctctcagttttttttttgtgttttttg
tccctctttgtttcacttgagaacaacccccttttgaactcgatcaagaaagctaagtttgaaga
atcaaga**ATG(TTS of MYB51)**

**Supplemental File 1.** Promoter regions amplified in ChIP-PCR experiments.
